# Therapeutic-induced chromatin remodeling via AP-1/SWI-SNF enhances BK Polyomavirus replication in urothelial cells

**DOI:** 10.64898/2026.09.11.750935

**Authors:** Subhajit Chatterjee, Lucy Langenberg, Jason T. Blackard, Stella Davies, Benjamin Laskin, Gabriel J. Starrett

**Affiliations:** Laboratory of Cellular Oncology, Center for cancer research, National Cancer Institute, National Institute of Health, Building 37, Room 4128, Bethesda, MD 20892-4263, USA; Division of Bone Marrow Transplantation and Immune Deficiency, Cincinnati Children’s Hospital Medical Center, USA; Division of Infectious Diseases, University of Cincinnati College of Medicine, Cincinnati, Ohio 45219, USA; Division of Nephrology, The Children’s Hospital of Philadelphia, USA

**Author notes:** To whom correspondence should be addressed Gabriel J. Starrett, Laboratory of Cellular Oncology, Center for Cancer Research, National Cancer Institute, National Institute of Health, Building 37, Room 4128, Bethesda, MD 20892-4263, USA.

**Keywords:** Hemorrhagic cystitis, BK polyomavirus, Chemotherapy, Immune conditioning, Transplantation, Chromatin remodeling

## Abstract

Hemorrhagic cystitis (HC) caused by BK polyomavirus (BKPyV) reactivation is a serious complication in immunocompromised patients especially in hematopoietic stem cell transplant recipients, yet the effects of conditioning regimens on viral replication remain undefined. Current therapies for HC are largely supportive, and no specific antiviral drugs are approved for BKPyV reactivation, underscoring an urgent need for new interventions. Using a cellular model of HC, we provide the first mechanistic evidence that physiologic concentrations of genotoxic chemotherapy, immune-conditioning, and antiviral exposures commonly enhance BKPyV replication by reprogramming host chromatin remodeling machinery independently of immune suppression. Transcriptomic and epigenomic profiling of urothelial cells revealed that these treatments induce host and virus chromatin remodeling, specifically, the SWI/SNF ATP-dependent remodeling complex was activated and created a regulatory landscape enriched for AP-1 transcription factor motifs, implicating the AP-1/SWI-SNF axis as a key regulator of viral replication. Functional inhibition of SWI/SNF with AU-15330 and FHD-286 consistently suppressed BKPyV replication across all tested therapeutic conditions. Patient urine samples and cellular experiments demonstrated that these effects were independent of viral genomic changes. These findings suggest chromatin remodeling as the mechanistic driver of therapy-associated BKPyV reactivation and propose SWI/SNF inhibition as a translational therapeutic strategy to mitigate hemorrhagic cystitis.

## Introduction

Hemorrhagic cystitis (HC) is a painful and potentially life-threatening urological syndrome characterized by bladder mucosal injury, vascular compromise, and frank hematuria [1]. In hematopoietic stem cell transplant (HSCT) recipients, HC develops in approximately 7-54% of patients and is associated with significant morbidity, including urinary tract obstruction, prolonged hospitalization, and increased healthcare costs [2, 3]. HSCT patients receive a complex pharmacopeia to ablate their immune systems (immune-conditioning) prior to transplantation and to prevent complications from opportunistic infections, through high dose genotoxic agents and antivirals. Early-onset HC, occurring during or immediately after conditioning, is thought to be primarily driven by direct urothelial toxicity from drug treatment, such as alkylating agents like cyclophosphamide, ifosfamide, and busulfan [3]. On the other hand, late-onset HC, which typically arises post-engraftment, is linked to viral replication due to immunosuppression [3, 4]. Clinically, severe HC cases can involve massive bleeding and clot retention. In extreme cases bladder necrosis and hemorrhage may require surgical intervention and can be fatal. Studies show severe HC can cause 11% mortality in children [5]. Thus, HC represents a major clinical burden in transplant and oncology patients, driving greater length of stay, morbidity, and even mortality.

BK polyomavirus (BKPyV), a nonenveloped, circular double-stranded DNA virus of the Polyomaviridae family, infects most individuals during childhood and persists in an episomal form within urinary tract [6]. The BKPyV genome is packaged and delivered with host-derived histones into a minichromosome, providing a chromatinized template for viral transcription and replication. BKPyV minichromosomes carry diverse histone post-translational modifications, including extensive histone acetylation, indicating that the viral genome is subject to epigenetic regulation throughout the viral life cycle [7]. Central to this regulation is the noncoding control region (NCCR), which contains the viral origin of replication together with bidirectional promoter/enhancer elements that control early and late gene expression [8]. Alterations in NCCR architecture can profoundly affect BKPyV transcriptional activity and replication capacity, emphasizing the sensitivity of the viral regulatory region to both viral sequence and host regulatory inputs [9]. However, whether therapeutic stress alters host chromatin-regulatory machinery in a manner that changes the transcriptional competence of an otherwise sequence-stable BKPyV chromosome remains poorly understood.

While normally asymptomatic in immunocompetent hosts, BKPyV can replicate leading to high-level viruria, viremia, and direct cytolytic damage to urothelial tissue in immunosuppressed individuals, such as after HSCT or solid-organ transplantation, [6, 10]. In HSCT-associated late-onset HC, BKPyV is detected in the urine of 80% of affected patients, and prospective studies have demonstrated that rising BKPyV loads correlate with the onset and severity of HC [11]. More recently, early BKPyV viruria following allogeneic HCT was associated with an increased risk of subsequent BKPyV-associated HC, further emphasizing the importance of understanding the events that promote viral amplification during the peri-transplant period [12]. The prevailing model posits that conditioning regimens damage the urothelium, enabling BKPyV to replicate in regenerating epithelial cells, after which immune reconstitution triggers an inflammatory response that culminates in hemorrhage [13].

While HC most commonly occurs in transplant recipients, several case reports have also reported it in oncology patients receiving intensive chemotherapy, particularly alkylators and other urotoxic agents. For example, Cisplatin has been implicated as a pediatric series described a 15-year-old with medulloblastoma developed HC following cisplatin administration [1]. Additionally, children receiving high-dose cyclophosphamide exhibited persistent BKPyV viruria concomitant with prolonged hematuria, and clinical improvement followed cidofovir therapy [14]. Despite these observations, systematic investigations into BKPyV reactivation after chemotherapy, particularly in adult oncology populations, are lacking, resulting in a critical gap in our understanding of HC pathogenesis outside the transplant setting.

While BKPyV is a leading cause of complication in solid organ transplant recipient (SOTR) and HSCT recipients, current antiviral strategies lack any proven specificity or efficacy for BKPyV. Transplant protocols often include nucleoside analog antivirals such as acyclovir or ganciclovir to prevent herpesvirus complications, however, the lack of a viral thymidine kinase in the BKPyV genome renders these drugs ineffective against BKPyV replication [15]. Cidofovir has demonstrated in vitro inhibition of BKPyV DNA synthesis and has been used off-label in small case series to treat BKPyV-associated HC, sometimes with clinical benefit [16]. Nonetheless, no antiviral regimen has yet demonstrated efficacy in randomized trials. Importantly, many of these antivirals, chemotherapeutics, and immune conditioning agents are all eliminated through the kidneys and passed into the bladder still in an active form with up to 50% of the intravenous dose eliminated by this pathway within the first 24 to 48 hours [17–19].

Considering this, the bladder in particular may be exposed to more persistent and higher concentrations of these drugs compared to other organs, yet the effects of these individual drugs and in combination have not been studied in urothelial model systems nor in relation to BKPyV infection and disease.

To address this, we implemented a human urothelial cells (HBLAK)-based experimental systems (both monolayer and organotypic culture) that simulate chemotherapy-induced immunosuppression using representative chemotherapeutics (etoposide, cisplatin, and 5-flurouracil), immune-conditioning agents (cyclophosphamide, fludarabine, and busulfan), and antivirals (ganciclovir, acyclovir, and cidofovir). By quantifying rearranged and wildtype BKPyV replication under these conditions, our study aims to define how toxic drug exposures and combination regimens, mirroring clinical practice, affect BKPyV replication dynamics and contribute to HC pathogenesis in both transplant and nontransplant contexts.

In this study, we mechanistically investigate BKPyV reactivation following immune conditioning, chemotherapy, and antiviral-induced genotoxic stress, evaluating its potential contribution to urothelial damage and hemorrhagic cystitis. By investigating viral replication, host transcriptomic responses, and histopathological outcomes in in vitro and organotypic bladder models, our work provides mechanistic insights into BKPyV-associated bladder injury beyond traditional transplant contexts and underscores the need for virologic surveillance in patients at risk.

## Results

### Therapeutic conditioning promotes BKPyV replication

To model infection of the urothelium, spontaneously immortalized human urothelial cells (HBLAK) were infected with either 0.5 MOI archetype (wild type, strain Dik) or rearranged BKPyV (strain Gardner) at a multiplicity of infection (MOI) of 0.5. Rearranged BKPyV exhibited increased genome copies at day 3 post-infection, then increased markedly by day 5 (**Supplementary Fig. 1A**). In contrast, archetype BKPyV failed to detectably replicate by quantitative PCR (qPCR) (**Supplementary Fig. 1A**) even at doses up to 10 MOI over a 5-day period (**Supplementary Fig. 1B**). Because only rearranged BKPyV replicated efficiently in monolayer HBLAKs, most subsequent experiments employed the rearranged strain.

Next, to assess the effect of immune conditioning drugs on BKPyV replication, HBLAK monolayers were infected with rearranged BKPyV at 0.5 MOI and incubated for 3 days to establish viral replication. At 72 hours post-infection, cells were treated with cyclophosphamide at either IC_10_, IC_20_, or IC_30_, for 48 hours. The IC_10_ and IC_20_ concentrations, determined empirically in HBLAK cells (**Supplementary Fig. 2, Supplementary Table 1**), are similar to the IC_50_ doses reported for various cancerous cell lines, modeling how clinical regimens may inadvertently impact BKPyV replication in normal quiescent cells. qPCR conducted 48 hours after drug addition revealed that cyclophosphamide IC_10_ increased BK genome copy number to approximately 1.6-fold of untreated infected controls, whereas IC_20_ produced a 3.6-fold elevation (P = 0.0073) (**Fig. 1A**). Conversely, at IC_30_ concentrations BKPyV replication was found to be decreased (**Fig. 1A**).

**Figure 1.**
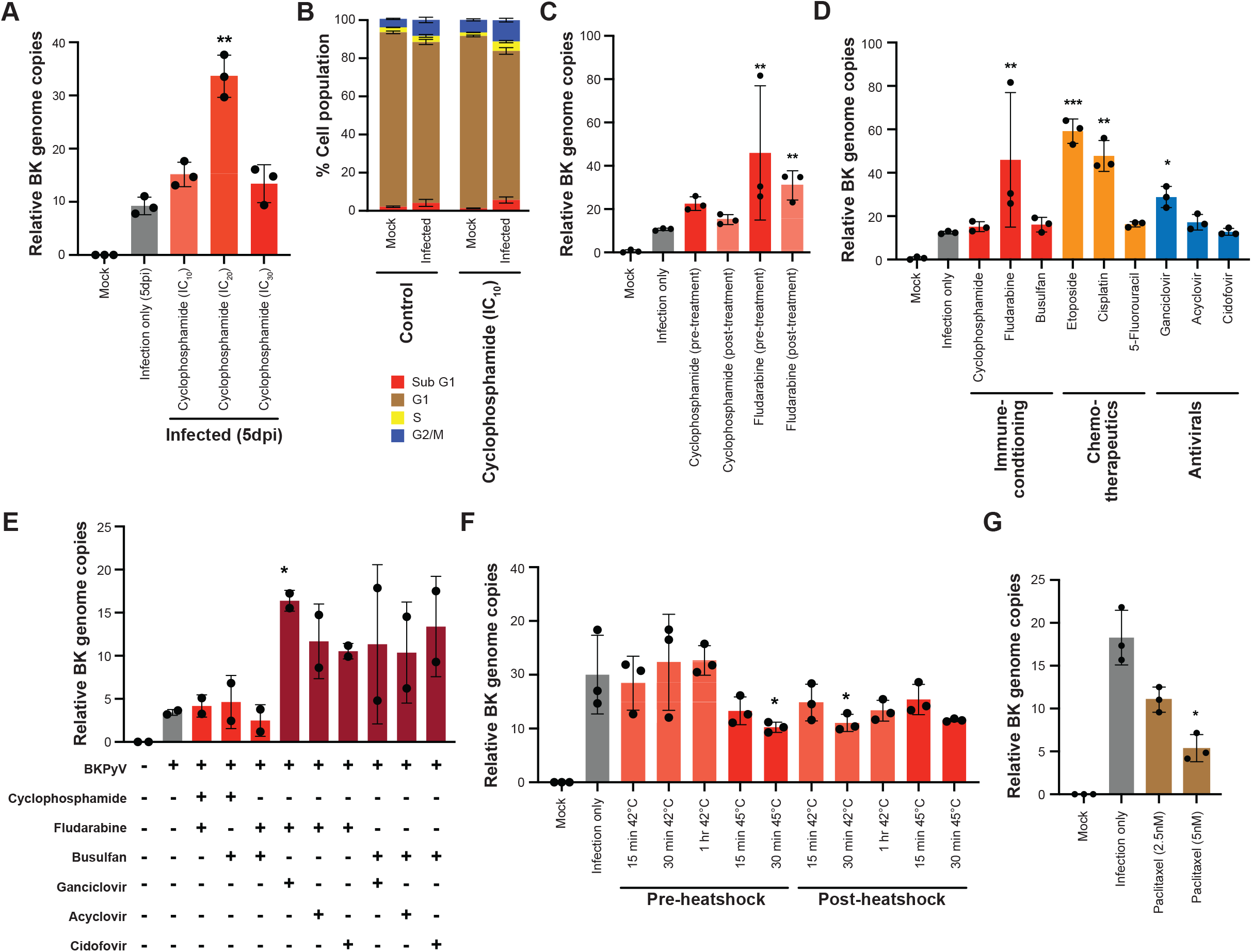
Sublethal therapeutic exposures enhance rearranged BKPyV replication in HBLAK cells. **(A)** HBLAK cells were infected with rearranged BKPyV (strain Gardner) at MOI 0.5 for 3 days and subsequently treated with cyclophosphamide at the indicated IC_10_, IC_20_, or IC_30_ concentration for 48 hours. BKPyV genome abundance was quantified by T-antigen-targeted qPCR at 5 days post-infection and normalized as described in Materials and Methods. **(B)** Cell-cycle distribution of mock or rearranged BKPyV-infected HBLAK cells untreated or treated with cyclophosphamide IC_10_. The percentages of cells in Sub-G1, G1, S, and G2/M were determined by propidium iodide staining and flow cytometry. **(C)** Comparison of pre-infection and post-infection exposure to cyclophosphamide or fludarabine. For pre-infection treatment, cells were exposed to drug for 6 hours, washed, and infected for 5 days. For post-infection treatment, cells were infected for 3 days and then treated for 48 hours. **(D)** BKPyV genome abundance following 6-hour IC_10_ pre-treatment with the indicated immune-conditioning, chemotherapeutic, or antiviral agent, washout, and infection for 5 days. **(E)** BKPyV genome abundance following 6-hour exposure to the indicated combination regimens, followed by washout and infection for 3 days. ‘Plus’ and ‘minus’ symbols indicate compounds included in each treatment. **(F)** BKPyV genome abundance following heat shock at the indicated temperature either before or after infection. **(G)** BKPyV genome abundance following treatment with the indicated paclitaxel concentration. Data represent the mean ± SD from three independent biological experiments. Statistical comparisons were performed using one- or two-way ANOVA (where applicable) using Prism v10 software (GraphPad).

Because DNA viruses rely on host cellular machinery for genome replication, now the question arise whether BKPyV infection or sublethal therapeutic exposures could alter host cell-cycle progression. Flow cytometric analysis revealed no significant differences in the distribution of cells across Sub-G1, G1, S, or G2/M phases between mock and infected conditions. Similarly, treatment with IC_10_ concentrations of Cyclophosphamide did not induce measurable cell-cycle perturbations. (**Fig. 1B**). Analogous experiments with etoposide further validated that there was of cell cycle profiling upon IC_10_ treatment concentrations in both BKPyV (archetype and rearranged) infected and mock conditions (**Supplementary Fig. 3A**). These findings indicate that enhanced BKPyV replication under these conditions occurs independently of overt cell-cycle dysregulation.

To determine the influence of exposure versus infection timing, HBLAKs underwent two regimens with IC_10_ cyclophosphamide or fludarabine: (1) ‘post-infection treatment’, in which cells were infected at 0.5 MOI BKPyV for 3 days and then treated with drug for 48 hours; and (2) ‘pre-infection treatment’, in which cells were treated with drug for 6 hours, washed, and then infected at 0.5 MOI BKPyV for 5 days without further drug exposure. By day 5, pre-treatment cyclophosphamide yielded an approximately 2.1-fold increase in BK genome copy relative to infected controls, compared to a 1.6-fold increase when cyclophosphamide was added post-infection. Similarly, fludarabine pre-treatment produced a 4.3-fold elevation (P = 0.0047), compared to a 2.9-fold for post-infection fludarabine (P = 0.0062) (**Fig. 1C**).

Analogous comparisons for etoposide (IC_10_) and cisplatin (IC_10_) confirmed that short genotoxic pre-infection exposure primes HBLAKs for BKPyV replication resulting in higher viral genome copies than post-infection treatment (**Supplementary Fig. 3B**).

To explore the effect of different therapeutic regimens, we next pre-treated HBLAKs for 6 hours with IC_10_ concentrations of immune-conditioning agents, chemotherapeutics, and antivirals, and then washed the cells and infected at 0.5 MOI for 5 days. At day 5, qPCR analysis demonstrated that several IC_10_ treatment significantly enhanced BKPyV replication relative to infected controls: etoposide (4.8-fold; P = 0.001), cisplatin (3.8-fold; P = 0.003), 5-flurouracil (1.3-fold; P = 0.1941), cyclophosphamide (1.2-fold; P = 0.3301), fludarabine (3.7-fold; P = 0.0054), busulfan (1.3-fold; P = 0.2862), ganciclovir (2.3-fold; P = 0.018), acyclovir (1.4-fold; P = 0.2279), and cidofovir (1.1-fold; P = 0.963) (**Fig. 1D**). These IC_10_ doses mimic the exposures that normal urothelial cells may experience in transplant and chemotherapy settings, underscoring how even low-level conditioning can potentiate BKPyV infection. Comparable experiments using IC_20_ concentrations for 6 hours followed by a very low infection concentration (0.1 MOI) for 3 days yielded parallel enhancement patterns (**Supplementary Fig. 3C**).

Because several of the replication-enhancing agents can induce DNA damage at higher concentrations, we next considered whether the sublethal IC_10_ exposures used in our model elicited detectable activation of the canonical ATM- or ATR-mediated DNA damage response. Western blotting was performed for total ATM/ATR and other DNA damaging markers like Chk-1, Chk-2, pChk-1, pChk-2, BAX, and BCL-XL following treatment under the same sublethal exposure conditions used in the replication experiments. Consistent with the absence of measurable cell-cycle redistribution, IC_10_ treatment did not produce a reproducible increase in ATM or ATR phosphorylation relative to the corresponding untreated controls (**Supplementary Fig. 3D**). Total ATM and ATR protein abundance also remained largely unchanged.

Together, these findings indicate that the replication-enhancing treatment conditions do not elicit sustained activation of the canonical ATM/ATR signaling pathways detectable at the time point examined. Thus, the increased BKPyV replication observed following sublethal therapeutic conditioning cannot be readily explained by overt cell-cycle redistribution or sustained activation of the canonical ATM/ATR DNA damage response.

Transplant recipients typically receive a multidrug regimen to fully eliminate immune cells prior to engraftment and prevent opportunistic viral infections. To model this, HBLAK cells were co-treated with IC_10_ immune-conditioning agents (Fludarabine+ Cyclophosphamide, Busulfan+ Cyclophosphamide, or Fludarabine+ Busulfan) and combined immune-conditioning and antivirals (Fludarabine+ Ganciclovir, Fludarabine+ Acyclovir, Fludarabine+ Cidofovir, Busulfan+ Ganciclovir, Busulfan + Acyclovir, and Busulfan + Cidofovir) for 6 hours. After washing, cells were infected at 0.5 MOI for 3 days. By day 3, qPCR revealed that most of the combination treatments produced an additive effect, particularly when cells were pre-treated with immune-conditioning drugs and antivirals (**Fig. 1E**). These results indicate that concurrent low-level conditioning and antiviral prophylaxis create an environment even more permissive for BKPyV replication than individual agents alone.

To ensure that the observed increase in BKPyV replication was not due to a nonspecific stress response, we subjected HBLAK monolayers to two non-genotoxic stressors, heat shock and paclitaxel exposure. For heat shock, HBLAKs were incubated at 42°C for 15 minutes, 30 minutes, and 1 hour or 45°C for 15 minutes and 30 minutes either before BKPyV infection (“pre-heat shock”) or at 3 days of post-infection (“post-heat shock”), then returned to 37°C. At 5 days post-infection (0.5 MOI), qPCR data revealed no significant increase in viral copy numbers as compared with infected controls maintained continuously at 37°C, indicating that acute thermal stress alone does not enhance BKPyV susceptibility or replication (**Fig. 1F**). Importantly, Western blot analysis showed induction of the cellular heat-shock response (HSP-90 and HSP-70) under these conditions (**Supplementary Fig. 3E**), indicating that the heat-shock treatment elicited a measurable stress response without promoting BKPyV replication. Likewise, HBLAKs treated with paclitaxel (2.5 and 5 nM for 6 hours) before initial BKPyV infection failed to show elevated viral genome copies at day 5 (**Fig. 1G**). Dose-response viability profiling confirmed that the paclitaxel concentrations used in these experiments were within a sublethal range, retaining approximately 84–89% cellular viability (**Supplementary Fig. 3E**).

To evaluate the effects of extended drug exposure, 48 hours BKPyV-infected HBLAKs were treated with cyclophosphamide or Fludarabine IC_10_ for 5 days without media replacement. At day 5 post-infection, BK replication was further noted higher as compared to when the cyclophosphamide or fludarabine-containing medium was replaced at day 2 and cells cultured for an additional 3 days in fresh, drug-free medium (**Supplementary Fig. 3F**). Analogous comparisons for representative chemotherapeutic agent, etoposide further validated the enhanced BKPyV replication during extended drug exposure (**Supplementary Fig. 3G**). Also, at IC_30_ for a 6 hours pre-treatment with etoposide, BKPyV replication decreased to 1.4-fold relative to the observed BKPyV replication under IC_20_ concentrations for a 6 hours pre-treatment, thereby validating the cytotoxicity limiting viral expansion (**Supplementary Fig. 3H**). To determine whether therapy-induced BKPyV activation was restricted to urothelial cells, we evaluated BKPyV replication in RPTECs, a physiologically relevant renal epithelial cell model that supports BKPyV replication. Similar to HBLAK cells, genotoxic stress conditions enhanced BKPyV replication in RPTECs, although the magnitude of induction varied across therapeutic exposures. Thus the therapy-mediated enhancement of BKPyV replication is not unique to a single epithelial lineage and may reflect a conserved host response to genotoxic stress (**Supplementary Fig. 3I**).

### Therapeutic conditioning increases cellular permissiveness to infection and facilitates archetype BKPyV replication in 2D and 3D models

To determine whether the drug-induced enhancement of BKPyV replication in HBLAK monolayers is due to increased replication rate and/or increased cellular permissiveness, we performed immunofluorescence for T antigen (TAg) and VP1 in monolayers following rearranged BKPyV infection. HBLAKs were pre-treated and then washed and infected at 0.5 MOI. After 3 days, cells were fixed and stained for TAg and VP1. Initial microscopic data indicated that genotoxic therapeutics could elevate urothelial cells susceptibility to BKPyV infection (**Fig. 2A**). However, quantification of TAg- and/or VP1-positive cells across replicates confirmed strong statistically significant increase in the percentage of infected cells under etoposide IC_10_ conditioning regimen compared with the untreated, infected control (p<0.0001). Etoposide produced the greatest shifts in permissiveness, while all other treatment conditions exhibited modest changes (**Fig. 2B**). These findings suggest that therapeutic conditioning can increase HBLAK susceptibility to BKPyV infection, however, this is not the sole reason behind elevated BKPyV replication.

**Figure 2.**
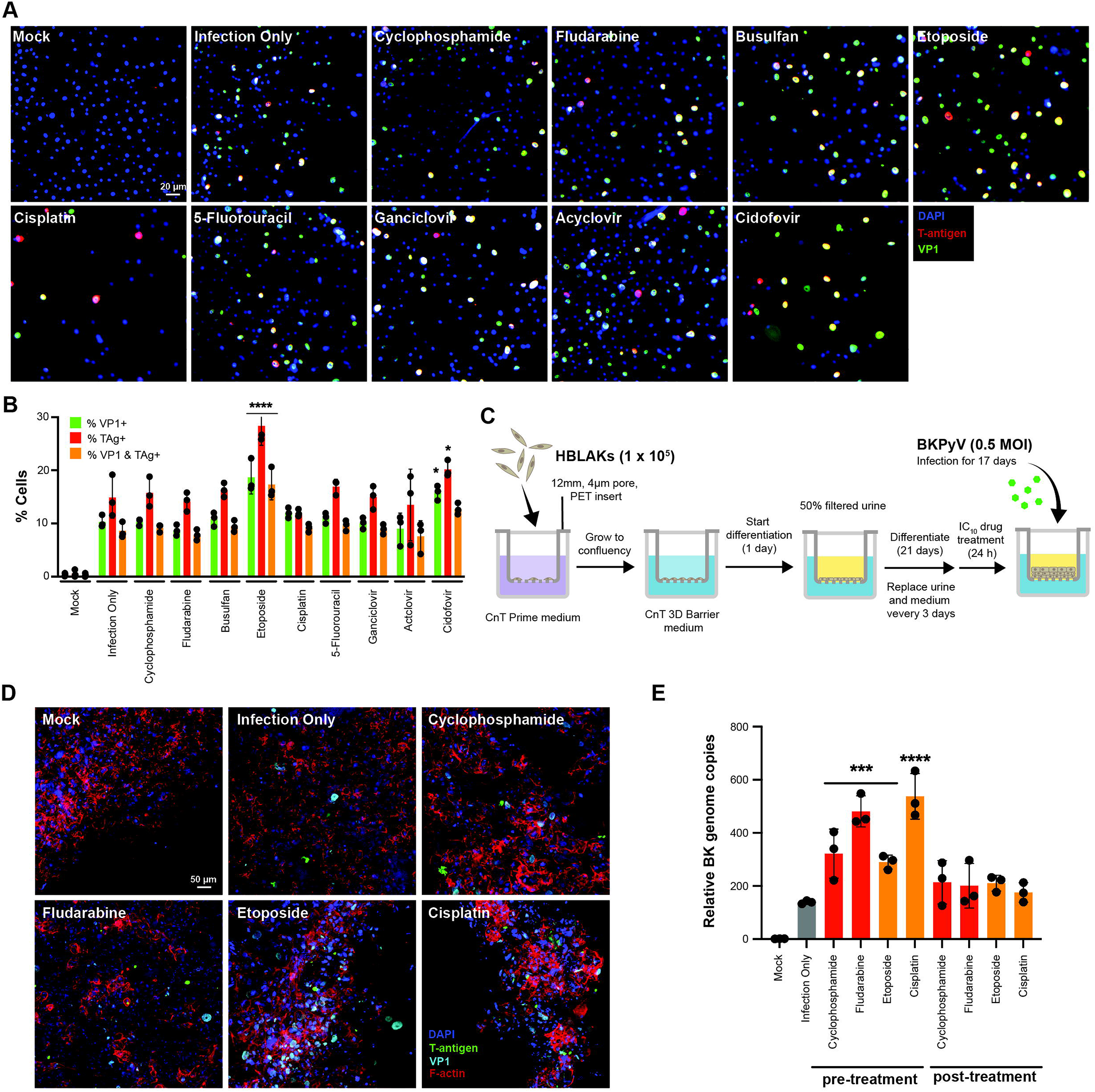
Therapeutic conditioning increases viral protein expression and supports archetype BKPyV replication in differentiated urothelium. **(A)** Representative immunofluorescence images of HBLAK cells pre-treated for 6 hours with the indicated IC_10_ agent, washed, and infected with rearranged BKPyV at MOI 0.5 for 3 days. Cells were stained for BKPyV large T antigen (TAg; red), VP1 (green), and nucleus (DAPI; blue). Scale bar is 20 μm. **(B)** Quantification of TAg-positive, VP1-positive, and TAg/VP1-double-positive cells from the conditions shown in (A) by using Fiji (ImageJ). **(C)** Experimental workflow for organotypic HBLAK raft cultures subjected to drug pre-treatment and archetype BKPyV Dik infection as described in Materials and Methods. **(D)** Representative confocal images of differentiated rafts mock-infected or infected with archetype BKPyV after the indicated pre-treatment. Tissues were stained for TAg, VP1, F-actin, and DAPI. Scale bar = 50 μm. **(E)** Archetype BKPyV genome abundance in organotypic cultures subjected to pre-infection or post-infection treatment with the indicated agents. Data represent the mean ± SD from three independent experiments. Statistical analysis was performed using one-way ANOVA using Prism v10 software (GraphPad).

To assess whether this observed elevated BKPyV replication phenomenon extends to a multilayered, more physiologically relevant context, and to determine whether archetype BKPyV behaves similarly, we generated differentiated organotypic cultures of HBLAKs. Once full differentiation was confirmed by microscopy (presence of basal and apical layers) (data not shown), cultures were pre-treated for 24 hours with couple of representative immune-conditioning drugs (cyclophosphamide and fludarabine) and chemotherapeutic agents (etoposide and cisplatin) at their IC_10_ concentrations, washed, and inoculated with archetype BKPyV for 17 days (**Fig. 2C**). In a parallel set of experiment, after differentiation, cells were infected with archetype BKPyV for 10 days, followed by drug treatment for 48 hours, and media replacement and retention for 6 days in media/urine conditions. After that, tissues were fixed and processed for confocal imaging (**Supplementary Fig 4A**).

Confocal Z-stack microscopy revealed minimal TAg or VP1 expression in untreated, infected cultures. In contrast, cyclophosphamide pre-treated cultures exhibited TAg positivity, with extensive strong VP1 signal across basal and spinous layers. Fludarabine pre-treated yielded a similar pattern, with TAg staining and widespread VP1 signal. Retrospective experiments with chemotherapeutic agents also validated the elevated archetype BKPyV replication in the 3D system (**Fig 2D**). This observation confirms that archetype BKPyV, typically non-replicative in monolayers (**Supplementary Fig. 1**), can replicate robustly under therapeutic conditioning within a 3D differentiation context.

qPCR quantification of archetype BK genome copies in 3D culture-derived DNA corroborated the microscopy findings. In the drug pre-treatment conditions, cyclophosphamide-conditioned culture harbored nearly 2.3–fold higher viral copy versus untreated, infected cultures (P < 0.001), whereas fludarabine, etoposide, and cisplatin pre-treated rafts exhibited nearly 3.5-fold (P < 0.001), 2.1-fold (P < 0.001), and 3.9-fold (P < 0.0001) increases (**Fig. 2E**). Post-drug treatment conditions also exhibited enhancement of archetype BKPyV replication (**Supplementary Fig 4B, Fig. 2E**). These values support the immunofluorescence data and demonstrate that archetype BKPyV replication is also significantly augmented by genotoxic immune conditioning in organotypic cultures.

### Therapy-associated BKPyV amplification occurs without viral genomic adaptation

To determine whether the increase in BKPyV replication following therapeutic conditioning was accompanied by detectable changes in the viral genome, we performed complementary long- and short-read sequencing of BKPyV recovered from treated HBLAK cells and from longitudinal urine specimens collected from transplant recipients. Long-read sequencing was used to assess larger structural alterations in the viral genome, whereas short-read sequencing was used to examine single-nucleotide substitutions and small insertion/deletion variants. Clinical sampling was interpreted in relation to transplantation and, where available, conditioning exposure, allowing viral genomic changes to be evaluated in the context of longitudinal changes in BKPyV burden.

In HBLAK cells infected with rearranged BKPyV and exposed to sublethal immune-conditioning, chemotherapeutic, or antiviral agents, Oxford Nanopore sequencing revealed no a reproducible treatment-associated increase in large structural variants, including deletions, insertions, duplications, inversions, or microhomology-associated events, relative to untreated infected controls (**Supplementary Fig. 5A**).

Short-read Illumina sequencing of the corresponding viral DNA likewise showed no recurrent treatment-specific shift in the spectrum of single-base substitutions or emergence of a common mutational pattern across the different therapeutic conditions (**Supplementary Fig. 5B**). Although individual low-frequency sequence variants were detected, their distribution was not consistent with the acquisition of a reproducible treatment-associated BKPyV mutational signature. Thus, the enhanced viral replication observed following sublethal therapeutic exposure was not accompanied by detectable recurrent genetic adaptation of the viral genome under the conditions examined.

We next evaluated BKPyV dynamics in longitudinal clinical samples. Cohort 1 consisted of eight transplant recipients enrolled at the NIH Clinical Center, with urine specimens collected at multiple time points spanning the conditioning and transplantation period. BKPyV DNA was quantified by qPCR to follow changes in urinary viral burden over time (**Supplementary Fig. 6A**). In several individuals, BKPyV levels increased substantially during the post-transplant period, providing a clinical setting in which viral expansion could be compared with viral genome sequence stability.

To determine whether this pattern was also observed in an independent population, we analyzed longitudinal urine specimens from transplant recipients who developed BKPyV-associated hemorrhagic cystitis at the Cincinnati Children’s Hospital and the Children’s Hospital of Philadelphia (cohort 2). Among 147 patients, 31 (21.1%) exhibited substantial increases in urinary BKPyV during the clinical course, with some patients showing increases of several orders of magnitude relative to earlier available measurements (**Supplementary Fig. 6B**). The timing and magnitude of viral shedding varied among individuals, and these observational data do not establish conditioning as the direct cause of reactivation. Rather, the two cohorts demonstrate that marked BKPyV expansion can occur during the conditioning/transplantation period and provide a clinical framework in which to ask whether viral amplification is accompanied by detectable viral genomic evolution. **Supplementary Tables 2 and 3** summarize patient characteristics, treatment information, transplantation dates, and available urine collection time points.

Therefore, we examined longitudinal BKPyV sequence variation in the clinical specimens. Short-read sequencing of serial samples from the clinical cohorts showed variable low-frequency substitutions among individual patients but no consistent enrichment of a specific substitution class or common treatment-associated mutational signature (**Supplementary Figs. 7 and 8**). In several longitudinal comparisons, substantial changes in viral burden occurred without the appearance of a corresponding new substitution pattern. Long-read sequencing of cohort 1 provided an orthogonal assessment of viral genome architecture and similarly did not reveal reproducible emergence or expansion of large deletions, insertions, duplications, inversions, or microhomology-associated structural variants in later samples relative to earlier patient-specific specimens (**Supplementary Fig. 9**).

Finally, we asked whether the increase in BKPyV DNA following therapeutic exposure was associated with a detectable change in viral DNA topology. Total DNA from infected HBLAK cells was treated with Plasmid-Safe ATP-dependent DNase, which preferentially degrades linear double-stranded DNA while preserving intact circular double-stranded DNA, and the remaining BKPyV DNA was quantified by qPCR. The Plasmid-Safe-resistant fraction was therefore used as an estimate of circular viral DNA, whereas the nuclease-sensitive fraction provided an estimate of linear viral DNA. Cyclophosphamide- and etoposide-treated samples showed circular and linear DNA fractions comparable to those of untreated infected controls. Cidofovir-treated samples showed a higher mean proportion of Plasmid-Safe-resistant viral DNA and a correspondingly lower nuclease-sensitive fraction, although this effect was variable between biological replicates. Across treatments, we did not observe a consistent increase in the proportion of linear BKPyV DNA (**Supplementary Fig. 10**). Thus, these findings therefore do not support a major treatment-associated accumulation of linear viral genomes or linear/concatemeric replication intermediates under the conditions examined, although this assay does not by itself distinguish among specific mechanisms of viral DNA replication.

### Transcriptomic profiling reveals shared host gene expression changes across conditioning regimens

To explore how chemotherapeutic, immune-conditioning, and antiviral treatments reshape host transcriptional responses, both in the presence and absence of BKPyV infection, we performed bulk total RNA sequencing on HBLAK cells under all treatment conditions. Principal component analysis (PCA) of normalized transcript counts showed that principal component 1 (PC1) accounts for 51% of the total variance, cleanly separating untreated, mock-infected samples from all other groups (**Fig. 3A**). Notably, untreated BKPyV-infected cells clustered closely with treated, mock-infected cells along PC1, indicating that drug conditioning induces a transcriptional program that partially overlaps with infection-driven changes. In contrast, treated, infected samples shifted further along PC2 (which explains 18% of variance), reflecting additional gene-expression alterations resulting from the combination of conditioning plus viral infection. These PCA patterns suggest that sublethal drug exposures alone elicit a host response that creates and infection-like state, thereby potentially facilitating BKPyV replication.

**Figure 3.**
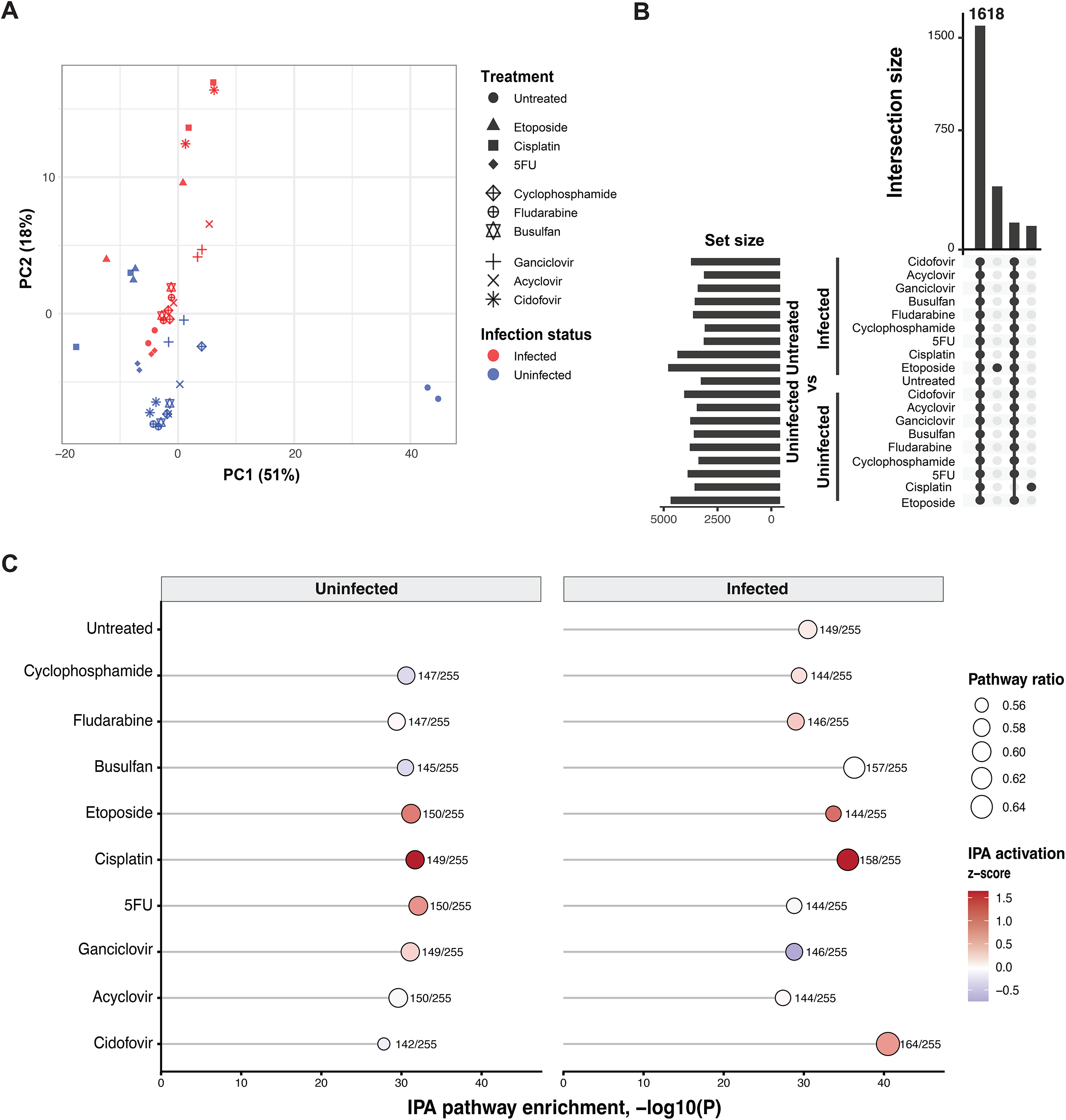
Therapeutic exposures induce a shared host transcriptional program enriched for chromatin-regulatory processes. **(A)** Principal-component analysis of normalized host RNA-seq counts from mock and rearranged BKPyV-infected HBLAK cells exposed to the indicated treatments. PC1 and PC2 account for 51% and 18% of total variance, respectively. Colors and symbols denote infection and treatment class as shown. **(B)** UpSet plot showing intersections of differentially expressed host genes across the indicated treatment and infection comparisons. 1618-gene intersection shared across all comparisons is highlighted. **(C)** Ingenuity Pathway Analysis of genes associated with chromatin organization. The x axis indicates pathway enrichment as −log_10_(P value), point size indicates pathway ratio, and color represents the predicted activation z score. RNA-seq was performed using two biological replicates per condition.

To dissect whether specific drug classes drive distinct expression signatures, we performed separate PCA for chemotherapeutic, immune-conditioning, and antiviral cohorts (**Supplementary Fig. 11A, C**, and **E**). In each subset, treated, mock-infected cells moved away from untreated, mock-infected controls along the primary principal component, confirming that each drug category remodels the transcriptome irrespective of viral exposure. Moreover, pairing these sub-analyses with the full-dataset PCA underscores that the overlapping transcriptional shifts are not limited to one drug class but rather are a general feature of genotoxic therapeutic regimens.

Next, we identified differentially expressed (DE) genes between each treatment condition (infected or mock-infected) and the untreated, mock-infected baseline. Across all regimens, regardless of chemotherapeutic, immune-conditioning, or antiviral origin, 1618 genes were consistently up- or downregulated relative to baseline (**Fig. 3B**). **Fig. 3B** shows those 1618 genes that were shared across all nineteen experimental groups, whereas flanking bars represent genes unique to individual or partial combinations of conditions. The prominence of the central intersection confirms a core transcriptional program induced by drug conditioning that persists in both infection and no-infection contexts. When split by drug class, 2029 genes are common to all chemotherapeutic treatments, 2172 are common to all immune-conditioning treatments, and 2102 are common to all antiviral treatments, emphasizing a shared host response across modalities (**Supplementary Fig. 11B, D**, and **F**).

To determine whether the transcriptional responses converged on common biological processes, we compared IPA canonical-pathway results across the 19 differential-expression contrasts. In addition to the cell cycle signatures, the IPA “Chromatin organization” pathway was recurrently and strongly enriched across both infected and mock-infected treatment conditions (**Fig. 3C**). Depending on the condition, 142– 164 of the 255 molecules assigned to this pathway were represented in the RNA-seq dataset. These molecules included components of ATP-dependent chromatin-remodeling complexes, histone-modifying enzymes, and other chromatin-associated regulators. However, the IPA activation z-scores varied across conditions, indicating that the pathway contained a mixture of upregulated and downregulated genes rather than a uniform directional response. Collectively, these findings demonstrate broad transcriptional perturbation of chromatin-regulatory processes across conditioning regimens.

Together, these transcriptomic findings suggested that genotoxic conditioning regimens, irrespective of drug class, induce a shared host gene-expression program that overlaps substantially with BKPyV infection signatures. The convergence of DE genes and pathways across all experimental groups underscores a unifying mechanism by which different conditioning agents prime urothelial cells for enhanced permissiveness to BKPyV.

### Therapeutic genotoxic stress promotes productive BKPyV transcriptional programs leading to infectious virus production

To determine whether therapy-associated genotoxic stress increases in BKPyV DNA abundance reflected productive viral transcription, we analyzed viral RNA abundance from RNA sequencing of infected HBLAK cells exposed to representative chemotherapeutic, immune-conditioning, and antiviral agents.

Total BKPyV transcript levels were markedly elevated following exposure to genotoxic chemotherapeutics relative to untreated infected controls, with cisplatin producing the strongest increase and etoposide showing a consistent but more moderate effect (**Fig. 4A**). In contrast, immune-conditioning agents induced smaller increases, while antiviral treatments restrained viral transcription relative to infected controls.

**Figure 4.**
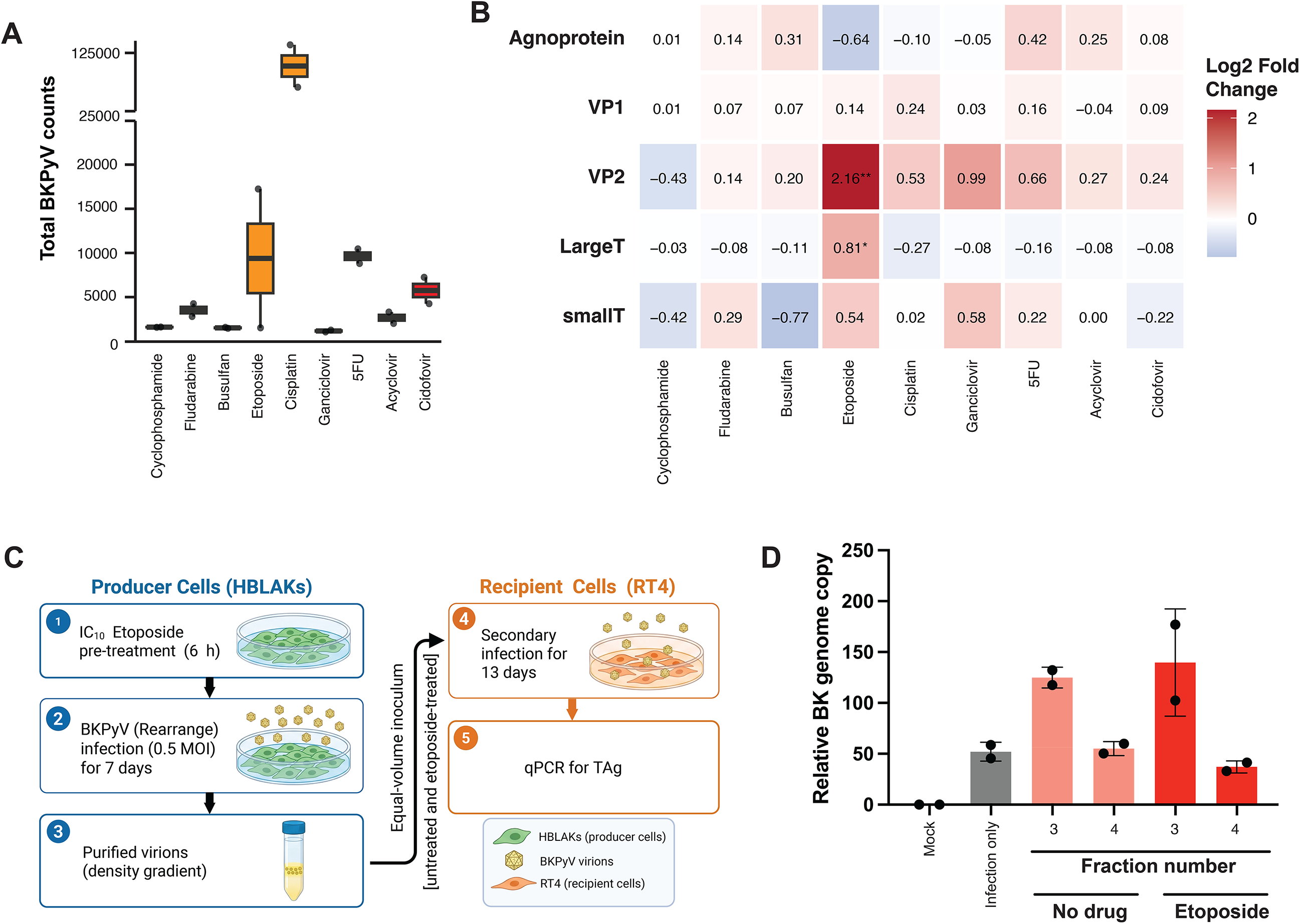
Genotoxic therapy increases BKPyV transcriptional output and production of infectious progeny. **(A)** Total normalized BKPyV transcript abundance in infected HBLAK cells exposed to the indicated treatment. Values represent DESeq2-normalized counts. **(B)** Heatmap showing log2 fold changes in BKPyV transcript abundance relative to untreated infected cells. Rows denote viral transcripts and columns denote treatment conditions. **(C)** Experimental strategy for the secondary-infection assay. HBLAK producer cells were treated with untreated or etoposide IC_10_ for 6 hours, washed, infected with rearranged BKPyV Gardner at MOI 0.5, and harvested at 7 dpi. Density-gradient virus purification was performed and fractions from cell lysates were used at equal volume to infect naive RT4 recipient cells, which were analyzed 13 days later. **(D)** BKPyV genome abundance in recipient RT4 cells after equal-volume inoculation with particle-enriched fraction 3 or 4 obtained from untreated or etoposide-treated producer HBLAKs. Data represent the mean ± SD from two independent preparations.

Analysis of BKPyV transcript composition demonstrated that genotoxic therapeutics altered viral gene expression in a gene- and treatment-specific manner. Etoposide produced the strongest transcriptional response, characterized by preferential induction of VP2 and increased large T-antigen expression (**Fig. 4B**). This selective pattern suggests that genotoxic stress does not simply increase global viral transcription but instead reshapes the viral transcriptional program toward specific replication-associated outputs. In comparison, immune-conditioning and antiviral treatments produced more modest and heterogeneous changes, without coordinated induction of either early or late viral genes. Collectively, these findings indicate that BKPyV transcriptional responses are shaped by the specific therapeutic context, with chemotherapeutics producing the most pronounced gene-selective activation and immune-conditioning conditions exerting comparatively limited effects on viral transcript composition.

To determine whether the therapy-associated increase in BKPyV DNA and viral transcription resulted in production of authentic infectious progeny, viral particles were purified from untreated and etoposide-treated BKPyV-infected HBLAK cultures at 7 days post-infection. Particle-containing fractions isolated from cell lysates and culture supernatants were subsequently used to infect naive RT4 cells, which were maintained for an additional 13 days before analysis (**Fig. 4C**). Recipient cells exposed to purified preparations exhibited BKPyV genome accumulation with viral T antigen expression, demonstrating that the recovered material contained infectious BKPyV particles capable of initiating secondary infection (**Fig. 4D**). Thus, genotoxic therapy does not simply increase intracellular viral DNA abundance but promotes completion of the BKPyV replication cycle and production of infectious virus.

### Chromatin landscape remodeling by conditioning regimens revealed by ATAC-seq

RNA-seq pathway analysis had highlighted altered expression of cell-cycle–related genes across conditioning regimens. To determine whether these transcriptomic signals corresponded to actual cell-cycle arrest or accumulation, we revisited the flow-cytometry–based cell-cycle profiling on HBLAKs treated with IC_10_ cyclophosphamide or etoposide (with and without BKPyV infection). As shown in **Fig. 1B** and **Supplementary Fig. 3A**, neither drug induced a significant shift in the proportion of cells in G_0_/G_1_, S, or G_2_/M phases compared to untreated controls. All drug treated infected HBLAKs also showed a nearly identical distribution. These negative findings confirm that enhanced BKPyV replication under sublethal conditioning is not driven by overt cell-cycle arrest or S-phase enrichment, motivating a focus on chromatin remodeling as the likely mechanism.

Comparative RNA-seq pathway analysis revealed recurrent enrichment of the chromatin organization pathway across infected and mock-infected treatment conditions. Although this result demonstrated extensive transcriptional perturbation of chromatin-associated genes, RNA-seq alone could not establish whether the accessibility of regulatory DNA elements was altered. We therefore performed ATAC-seq to directly determine whether conditioning regimens reshape the host chromatin-accessibility landscape. We performed assays on HBLAK cells subjected to representative drugs from each category, cyclophosphamide, fludarabine, and busulfan (immune conditioning); etoposide (chemotherapy); and cidofovir (antiviral), which previously showed pronounced enhancement of BKPyV replication.

Principal component analysis (PCA) of genome-wide ATAC-seq signals revealed a striking separation between drug-treated and untreated samples, with principal component 1 (PC1) accounting for 66% of the total variance (**Fig. 5A**). Unlike our RNA-seq results, untreated infected and untreated mock-infected samples clustered tightly together, indicating that the most prominent chromatin accessibility changes were treatment-induced rather than infection-driven. This again highlights that the sublethal doses of therapeutic regimens are sufficient to initiate a chromatin state resembling early infection or permissiveness.

**Figure 5.**
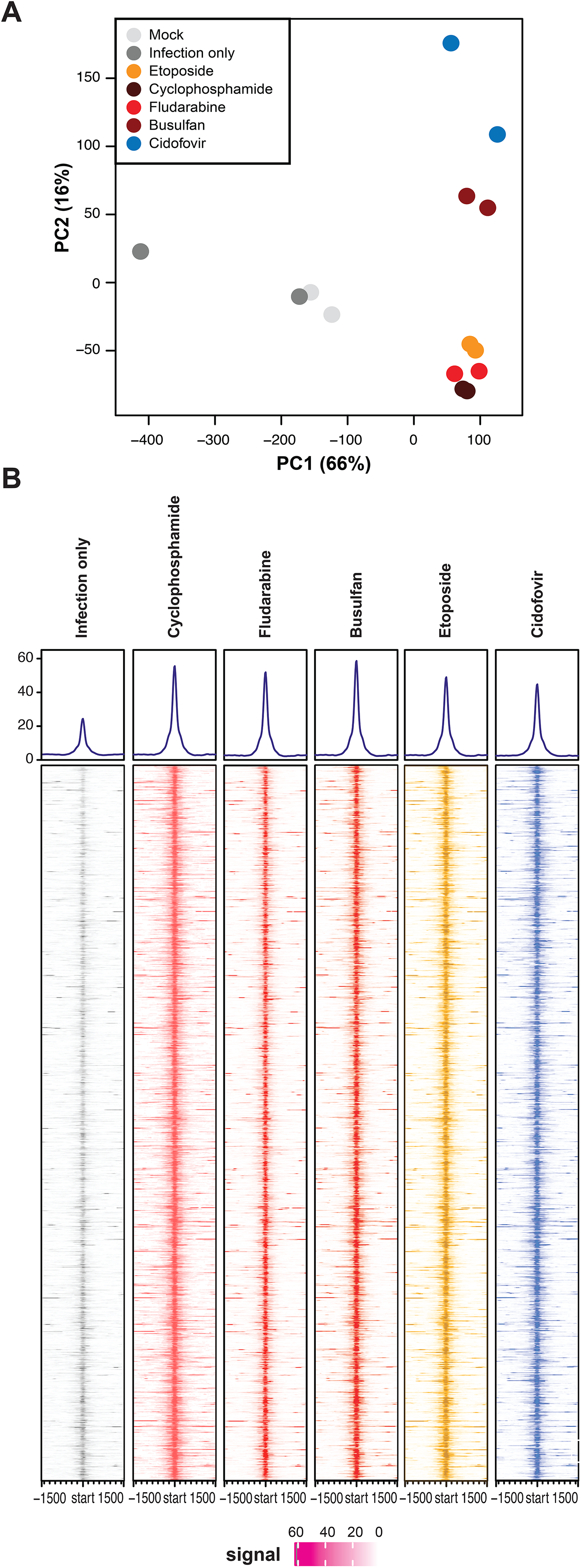
Therapeutic exposures remodel the host chromatin-accessibility landscape. **(A)** Principal-component analysis of normalized ATAC-seq signal from untreated, infected-control, and treatment-exposed HBLAK cells. PC1 and PC2 account for 66% and 16% of total variance, respectively. **(B)** Average normalized ATAC-seq signal and heatmaps centered on consensus accessible-region summits ±1.5 kb for the indicated conditions. Consensus peaks were displayed using a common signal scale across all conditions from two biological replicates per condition.

To visualize global chromatin remodeling trends, we generated a heatmap with average profile plots centered on all accessible peaks (**Fig. 5B**). All treatment conditions displayed elevated peak intensities relative to untreated controls, reinforcing that sublethal drug exposure leads to a more open chromatin state. This elevated accessibility may facilitate BKPyV replication upon infection, consistent with the increased viral genome load observed previously.

### Integration of chromatin accessibility and transcriptomic responses identifies shared promoter-associated genes and regulatory motifs

To determine which genes exhibit both increased promoter accessibility and altered expression under conditioning, ATAC-seq peaks were annotated using ChIPseeker, focusing exclusively on promoter regions defined as ±2 kb around transcription start sites (TSS). Intersection analysis across all five treatment conditions, revealed 2163 promoter-associated genes whose regulatory regions were consistently more accessible in every drug-treated sample (**Fig. 6A**). In addition to this, 76 genes were promoter-associated in all the treated and untreated infected conditions as compared to mock-infected untreated control (**Fig. 6A**). This core set of promoters likely represents a common chromatin signature by which diverse agents prime HBLAKs for BKPyV replication.

**Figure 6.**
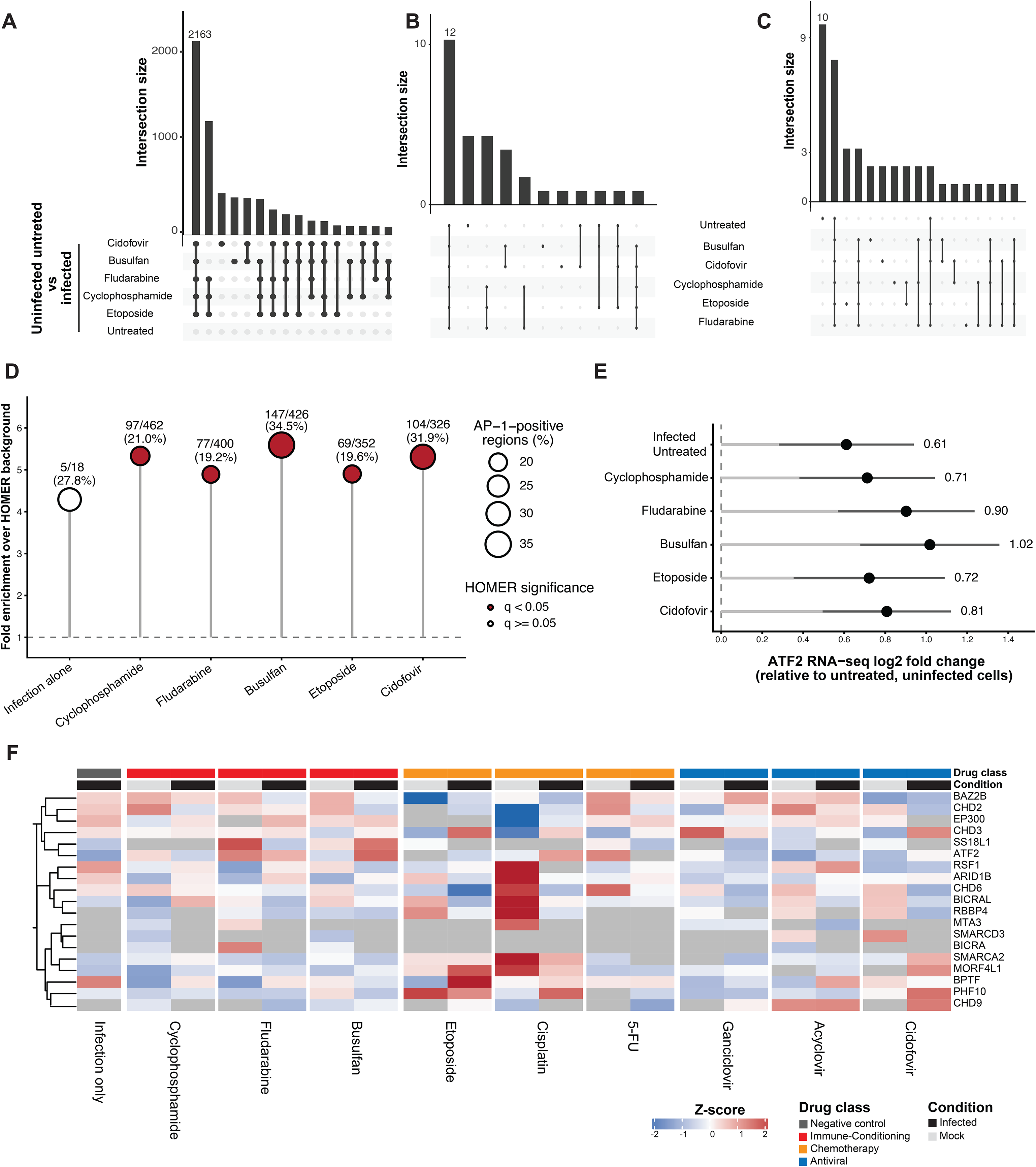
Integrated transcriptomic and chromatin-accessibility analyses nominate AP-1/ATF2 as a common regulator of the treatment-associated host state. **(A)** UpSet plot of promoter-associated genes linked to treatment-enriched ATAC-seq peaks. 2163-gene intersection shared across all the drug-treated conditions as highlighted, whereas 76-gene intersection shared across all comparisons. **(B)** Intersection of transcription-factor motifs significantly enriched in promoter-associated peaks among the shared treatment-responsive genes. 12 motifs common to all indicated conditions are highlighted. **(C)** Integrated analysis of RNA-seq differentially expressed genes and ATAC-seq promoter-associated genes identified 78 common genes. Motif enrichment within their associated accessible regions identified eight shared motifs. **(D)** AP-1 motif enrichment relative to HOMER background regions across the indicated comparisons. Point size denotes the number or proportion of AP-1-containing regions, and filled symbols indicate statistically significant enrichment. **(E)** ATF2 expression across the corresponding RNA-seq contrasts, shown as log2 fold change relative to the untreated mock-infected control.

Motif enrichment analysis using HOMER on the promoter peaks these 76 genes uncovered 12 transcription factor motifs that were significantly overrepresented in all six conditions (**Fig. 6B**). Strikingly, the majority belong to the basic leucine zipper (bZIP) family, including AP-1, Atf3, Bach, BATF, Fos, Fosl2, Fra1, Fra2, Jun-AP1, and JunB, and CTCFL (Zinc finger, ZF). The universal enrichment of AP-1 family motifs suggests that these factors may serve as central nodes in chromatin opening, coordinating a gene-expression program that mimics early viral infection and bolsters permissiveness. The presence of CTCFL (a CTCF paralog) hints at additional, perhaps epigenetic, regulation of chromatin loops at these promoters.

To refine this list to high-confidence functional targets, we overlaid the promoter-accessible ATAC peaks with differential gene-expression results from RNA-seq. Of the 2163 promoter-associated genes, 78 were also significantly differentially expressed in all the drug treatment conditions compared to untreated, mock-infected controls (**Supplementary Fig. 12**). These 78 genes thus represent loci where increased chromatin accessibility correlates directly with up- or downregulation of mRNA, pinpointing them as likely mediators of the drug-induced transcriptional priming observed earlier.

HOMER motif analysis on the subset of 78 differentially expressed, promoter-accessible genes identified 8 motifs that were enriched all the treated and untreated infected conditions as compared to mock-infected untreated control (**Fig. 6C**). Once again, bZIP family motifs, AP-1, Atf3, BATF, Fos, Fra1, Fra2, and JunB, dominated, along with CTCF (ZF). The recurrence of AP-1 and CTCF motifs among these 78 high-confidence genes underscores a model in which conditioning agents collectively activate an AP-1–driven enhancer–promoter network while CTCF may coordinate higher-order chromatin architecture to facilitate transcriptional responsiveness. The canonical AP-1 motif was significantly enriched in all five drug-treated datasets, with enrichment ranging from 4.89- to 5.59-fold over background. AP-1 motifs were detected in 19.2-34.5% of the integrated promoter regions, including 97 of 462 cyclophosphamide-associated regions, 77 of 400 fludarabine-associated regions, 147 of 426 busulfan-associated regions, 69 of 352 etoposide-associated regions and 104 of 326 cidofovir-associated regions. The infected-untreated dataset showed a comparable enrichment magnitude of 4.29-fold, with AP-1 motifs in 5 of 18 regions but did not reach significance after multiple-testing correction because the analysis was based on a small promoter set. The reproducible enrichment of AP-1 across the drug-treated conditions therefore nominated AP-1 as a candidate common regulator for subsequent chromatin-occupancy studies (**Fig. 6D**).

Among AP-1/ATF-family regulators, ATF2 was significantly upregulated across all six RNA-seq contrasts contributing to the integrated analysis, with log2 fold changes ranging from 0.61 to 1.02 (adjusted p<0.05) (**Fig. 6E**). While not consistently upregulated or downregulated, other components of the SWI/SNF complex known to be directed by AP-1 family transcription factors and involved in the regulation of the chromatin of other DNA viruses are expressed in HBLAK cells (**Fig. 6F**). Based on these observations and the convergence of our transcriptomic and accessibility datasets on chromatin remodeling, we asked whether SWI/SNF directly associates with the BKPyV chromosome under replication-enhancing treatment conditions. We performed CUT&RUN for the SWI/SNF ATPase BRG1/SMARCA4 and the BAF subunit ARID1A, together with ATF2 and the transcriptional coactivator p300 to determine whether these proteins associated with the BKPyV NCCR under replication-enhancing treatment conditions.

### Therapy promotes changes in BKPyV genome chromatin accessibility corresponding to SWI/SNF component binding

We next evaluated changes in chromatin accessibility under the treatment conditions. We observed the known nucleosome-free region over the NCCR as well as increased open chromatin over the first exon of Large T antigen (LT exon 1) in all conditions and a general increased accessibility of the late region and LT exon 1 only under drug treatment (**Fig. 7A**). Closer inspection of the NCCR and LT exon 1 region did show overlap with predicted AP-1-family member binding sites, consistent with the global enrichment observed in the human genome (**Fig. 7B**). Based on the combined AP-1-family motif enrichment, ATF2 expression and chromatin-organization pathway results, we examined whether ATF2 and associated chromatin-regulatory proteins occupied the BKPyV NCCR. CUT&RUN was performed for ATF2, p300, BRG1/SMARCA4 and ARID1A under control and a subset of replication-enhancing treatment conditions (**Fig. 7C**). Consistent binding of the differentially accessible region in LT exon 1 was observed, with no significant change in binding between conditions. However, and increase in H3K27 acetylation associated with gene enhancers was observed in this region and is one of the outcomes of SWI/SNF remodeling suggesting increased activity under etoposide treatment.

**Figure 7.**
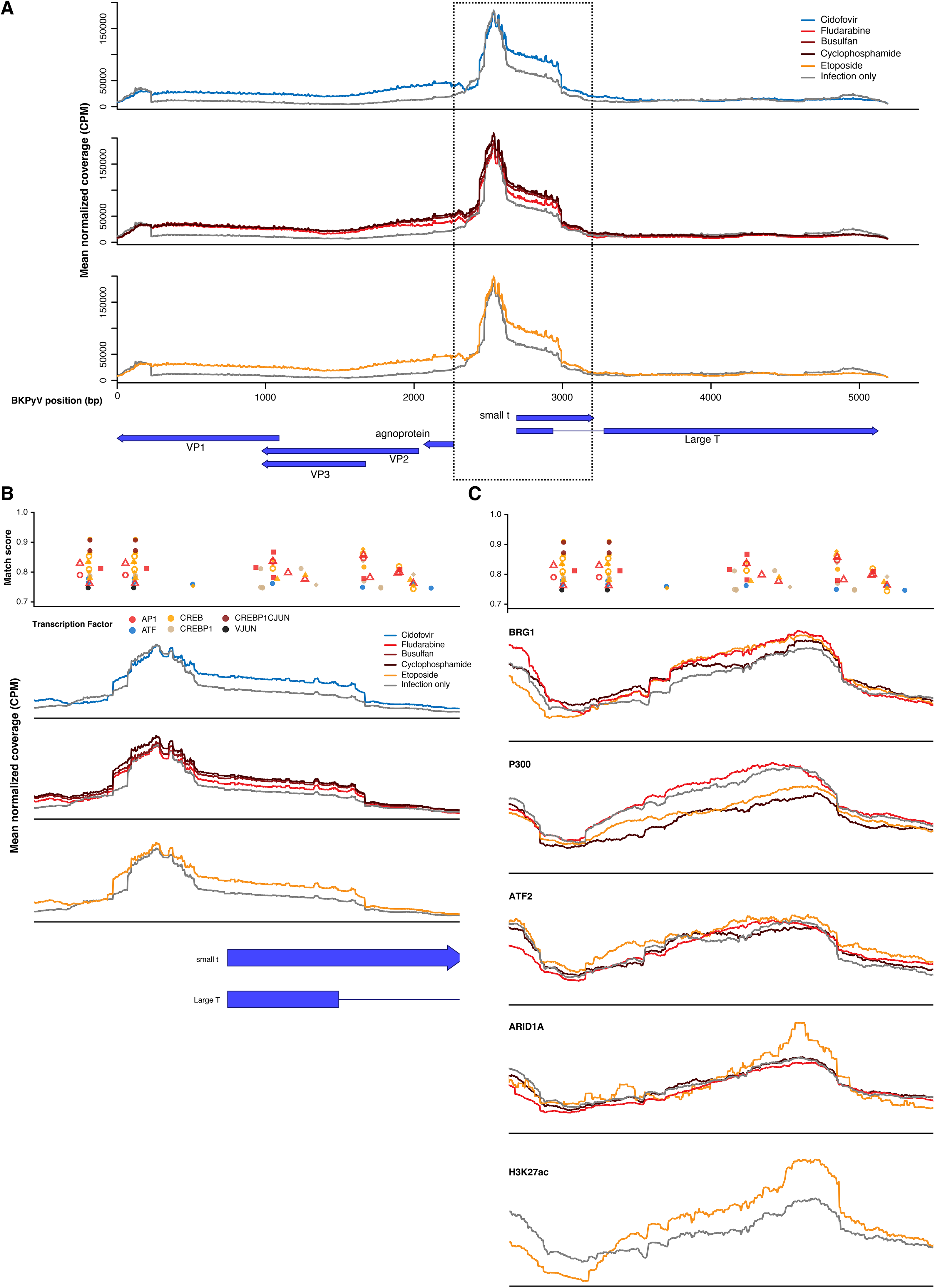
Therapeutics alter BKPyV chromatin accessibility, transcription factor binding, and histone modifications. **(A)** ATAC-seq chromatin accessibility of the BKPyV genome by drug treatment category with genes shown below (average of two biological replicates). **(B)** ATAC-seq chromatin accessibility of the region highlighted in A with a dotted box. Predicted transcription factor binding sites and match scores from TFBIND are indicated above. Alternative binding matrices for the same transcription factor are shown by different shapes. **(C)** CUT&RUN for SWI/SNF components, related transcription factors, and H3K27ac histone modifications of the same region as B. Predicted transcription factor binding sites and match scores from TFBIND are indicated above.

### SWI/SNF perturbation suppresses therapy-associated BKPyV replication

Having identified BRG1 and ARID1A association with the BKPyV NCCR, we next tested whether SWI/SNF function contributed to therapy-enhanced viral replication. HBLAK cells were pretreated with preclinical SWI/SNF ATPase degrader AU-15330 for 72 h before a 6-h exposure to the indicated genotoxic agent, followed by BKPyV infection and quantification of viral genome abundance 5 days later (**Fig. 8A**). This sequence was designed to establish SWI/SNF perturbation before genotoxic conditioning and thereby test whether SWI/SNF activity was required for development of the cellular state that supports enhanced BKPyV replication.

**Figure 8.**
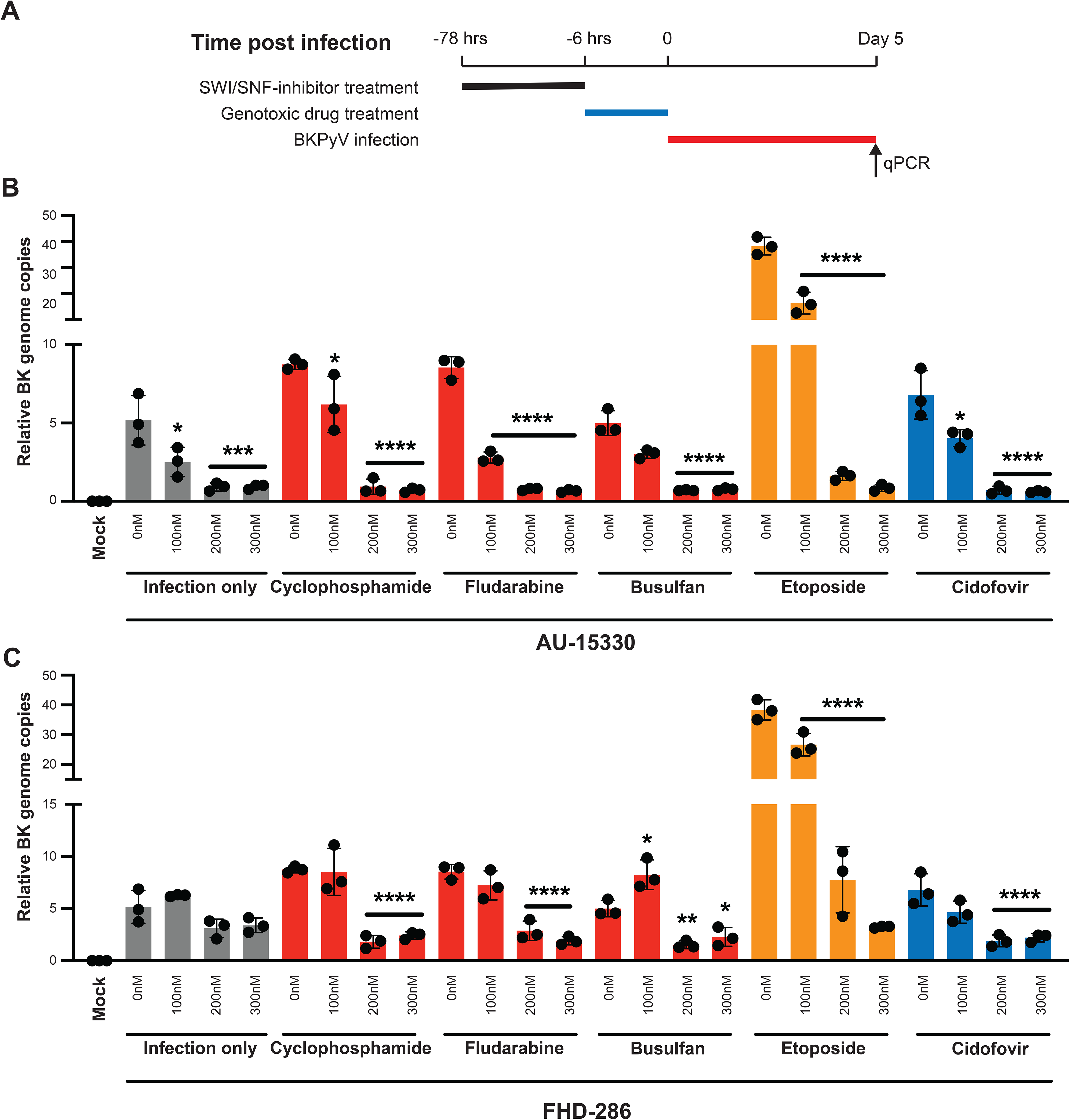
SWI/SNF perturbation suppresses therapy-associated BKPyV replication. **(A)** Experimental workflow for SWI/SNF perturbation: HBLAK cells were treated with SWI/SNF inhibitor for 72 hours, followed by washout and 6-hour exposure to the indicated therapeutic agent. Cells were washed, infected with rearranged BKPyV at MOI 0.5, and harvested at 5 dpi for qPCR analysis. BKPyV genome abundance in cells treated with untreated or 100 nM, 200 nM, or 300 nM AU-15330 (**B**) or FHD-286 (**C**) under the indicated conditions.. Data represent the mean ± SD from three independent biological experiments.

AU-15330 reduced BKPyV genome abundance at 100, 200, and 300 nM relative to the corresponding genotoxic-drug-only conditions (**Fig. 8B**). A similar reduction was observed with the mechanistically distinct SWI/SNF-directed compound FHD-286 (a clinical-stage BRG1/BRM inhibitor) (**Fig. 8C**).

Independent cell-viability profiling demonstrated that these concentrations were minimally toxic in HBLAK cells. AU-15330 maintained approximately 89% viability across the 100–300 nM working range, whereas FHD-286 maintained approximately 93% viability over the corresponding range (**Supplementary Fig. 13A-B)**. Thus, suppression of BKPyV replication occurred at concentrations that were not accompanied by extensive loss of cellular viability. Together, these pharmacologic results support a contribution of SWI/SNF activity to therapy-associated enhancement of BKPyV replication.

## Discussion

Polyomavirus-associated HC remains a major source of morbidity, and there are currently no proven interventions or prophylactic strategies that reliably prevent disease. The relative contribution of BKPyV replication versus drug-mediated urothelial injury has long been debated. Our findings suggest that these processes are not independent. Rather, clinically relevant therapeutic exposures appear to reprogram urothelial cells into a state that is more permissive for BKPyV transcription and replication.

BKPyV enters cells as a tightly packed, chromatinized episome. Prior single-cell RNA-seq studies of BKPyV infection models have shown that only a subset of exposed cells proceed to productive infection, likely reflecting differences in innate immune activation, cell-cycle state, transcription factor availability, or the capacity to remodel incoming viral chromatin [20, 21]. Consistent with the importance of epithelial cell state in vivo, recent single-cell profiling of human kidney allograft biopsies identified cellular-stress, wound-healing, and extracellular-matrix-remodeling programs in tubular epithelial cells during periods of increasing BKPyV viremia, including before overt BKPyV nephropathy [22]. Yang et al., also reported that ischemic injury promotes polyomavirus replication through a DNA-damage-response-associated program involving CDK1 in kidney cells [23]. Because many small DNA tumor viruses depend on host S-phase factors [21, 24, 25], we first asked whether sublethal conditioning altered cell-cycle distribution. Flow-cytometry profiling showed no significant shift in G₀/G₁, S, or G₂/M phases after IC_10_ cyclophosphamide or etoposide, regardless of infection status. In parallel, long- and short-read sequencing of treated HBLAK cells and clinical urine samples did not reveal consistent viral or host point mutations, mutational signature shifts, or structural rearrangements. These data argue against a model in which therapy-associated BKPyV activation is driven by simple S-phase enrichment, cell-cycle arrest, or viral genome evolution. Our observations and those by others suggest that multiple different clinically relevant tissue stresses may enhance BKPyV permissiveness and replication through tissue and context-dependent host pathways rather than through a single universal stress-response mechanism.

Our transcriptomic data support a host-state model influencing infection outcome. Bulk RNA-seq revealed that untreated BKPyV-infected cells cluster near genotoxic drug-treated, indicating that sublethal drug exposure alone induces many of the same transcriptional programs activated during early infection. Across heterogeneous conditioning regimens, 1618 genes were consistently up- or downregulated relative to untreated, mock-infected controls. This shared response suggests that diverse agents converge on a common cellular program that creates a permissive urothelial environment for BKPyV. While this did not correspond to significant increase in the total number of infected cells, it appears to drive accelerated replication dynamic as evident by virus copy numbers and virus transcriptional patterns where drug-conditioned cells showed a marked increase in late viral gene expression of the capsid proteins, VP1 and VP2.

Pathway analysis of the transcriptomic data suggested common changes in genes regulated by AP-1 (basic leucine zipper) family members, which are known to be stress responsive and can act as pioneer transcription factors by directing the SWI/SNF chromatin remodeling complex to closed chromatin [26]. Chromatin accessibility sequencing further supported this, showing untreated infected and untreated uninfected controls clustering closely together, whereas drug-treated cells acquired distinct accessibility profiles also enriched for AP-1 regulated genes. AP-1-responsive elements have previously been identified within the BKPyV regulatory region, and rearrangement of the NCCR can generate AP-1-binding enhancer modules that substantially increase early-promoter activity compared to the archetype virus [27]. CUT&RUN detected binding of BRG1/SMARCA4, p300, ARID1A, and ATF2 to the first exon of LT which corresponded to therapy-related increases in chromatin accessibility of that region as well as the entire late open reading frame. Pharmacologic perturbation further supported a functional contribution of SWI/SNF to enhanced BKPyV replication. This finding is consistent with the ability of other chromatinized DNA viruses to exploit SWI/SNF complexes to regulate transcription and replication of their episomal genomes [28, 29].

AP-1-family factors regulate papillomavirus transcription through cis-regulatory elements within the viral long control region; for HPV18, mutation of AP-1 elements can abolish early-promoter activity in keratinocytes [30]. SWI/SNF components likewise interact with HPV E2, and BRG1 ATPase activity enhances E2-dependent transcription and HPV18 DNA replication [31]. In herpesviruses (Epstein-Barr virus), BRG1 contributes directly to EBV viral-promoter activation and virion production, while recruitment of SWI/SNF to KSHV lytic promoters is required for efficient RTA-dependent reactivation [32].

While we did not observe activation of ATM/ATR from genotoxic drug concentrations used in this study, the essential role of ATM, ATR, and the DNA damage response (DDR) in BKPyV replication may still be relevant to these findings. ATM and ATR are central DDR kinases that phosphorylate broad protein networks after genotoxic stress and are strongly activated during BKPyV infection, and ATM signaling rapidly modifies chromatin surrounding DNA double-strand breaks through γH2AX-dependent recruitment of repair and remodeling factors [33]. Notably, BRG1/SMARCA4, the catalytic ATPase of SWI/SNF complexes, is a functional ATM substrate: ATM-mediated phosphorylation of BRG1 after DNA damage promotes its association with γH2AX-containing nucleosomes and supports double-strand break repair [34]. SWI/SNF activity has also been linked to ATR-associated replication-stress responses, as depletion of BRG1 and BRM attenuates full DDR activation, with a particularly prominent effect on ATR-CHK1 signaling and sensitivity to agents such as etoposide [35]. Thus, BKPyV infection induced DDR signaling may also be involved to further direct SWI/SNF activity and remodel host and viral chromatin in ways that favor BKPyV amplification and productive infection.

Our findings are also supported by the previous clinical observation that cyclophosphamide and cisplatin exposure specifically is associated with HC development [1, 3]. Considering that urinary excretion of chemotherapeutic agents and metabolites can increase local urothelial exposure, further work should define how other conditioning regimens, urinary metabolite levels, antiviral exposures, and host immune states interact to determine HC risk. These studies will be important for distinguishing patients who primarily require viral surveillance from those who may benefit from modified conditioning, intensified hydration, reduced genotoxic exposure when clinically feasible, or future host-directed therapies. More broadly, this work highlights the need to consider the urothelium not only as a passive target of drug toxicity, but as a dynamic tissue in which therapeutic stress can reshape viral susceptibility.

We acknowledge several limitations of this study. HBLAK monolayers and organotypic raft cultures recapitulate key features of human urothelium and enabled controlled mechanistic dissection, but they lack the immune, vascular, and stromal components that shape viral control in vivo. We did not directly interrogate innate or adaptive immune contributions, an important axis for future study. While clinical urine samples acutely showed increased BKPyV loads after conditioning, supporting clinical relevance, we cannot yet determine whether the chromatin remodeling observed in vitro occurs with the same timing or magnitude in patients due to limited access to bladder specimens in this patient population. In addition, although we tested representative conditioning agents and clinically relevant combination contexts, we did not exhaustively survey all regimens, doses, schedules, hydration protocols, or urinary drug concentrations used in practice.

In conclusion, common immune-conditioning, antivirals, and chemotherapy-associated exposures can reprogram urothelial chromatin and gene expression through an AP-1–SWI/SNF-associated axis, establishing a host permissive state that accelerates productive BKPyV replication without detectable global cell-cycle redistribution, sustained canonical ATM/ATR activation, or recurrent viral genetic adaptation. This host-directed mechanism provides a potential explanation for delayed HC after transplant conditioning and for HC in non-transplant patients receiving intensive chemotherapy, where genotoxic exposure, epithelial regeneration, viral amplification, and immune reconstitution all are variables in disease development. By identifying a targetable chromatin remodeling pathway as a possible core mechanism, our findings support expanded evaluation of BKPyV surveillance, risk stratification, reduced genotoxic exposure when feasible, optimized urothelial protection, and the development of new host-directed interventions as testable strategies to prevent disease.

## Author Contributions

Conceptualization, S.C. and G.J.S.; Methodology, S.C.; Investigation, S.C.; Formal Analysis, S.C. and G.J.S.; Data Curation, S.C.; Visualization, S.C.; Resources, G.J.S., B.L., J.T.B.; Writing – Original Draft, S.C.; Writing – Review & Editing, S.C., G.J.S., and all authors; Project Administration, S.C. and G.J.S.; Supervision, G.J.S.; Funding Acquisition, G.J.S, J.TB., B.L..

## Acknowledgements

The authors’ research is supported by the Center for Cancer Research, National Cancer Institute, National Institutes of Health Intramural Research Program project number ZIA BC 011894. The contributions of the NIH author are considered works of the United States Government. The findings and conclusions presented in this paper are those of the author and do not necessarily reflect the views of the NIH or the U.S. Department of Health and Human Services. We would like to thank Lorenzo Walker for sample retrieval and the DLM Microbiology laboratory for the initial testing. This work utilized the computational resources of the NIH HPC Biowulf cluster (https://hpc.nih.gov). NIDDK grant R01 DK125418 to JTB and BL.

## Materials and Methods

### HBLAK cell culture

Human immortal urothelial cells (HBLAK cells) were cultured in CnT-Prime epithelial proliferation medium (CELLnTEC) according to the manufacturer’s recommendations. Cells were maintained at 37°C in a humidified incubator with 5% CO₂ and were passaged before full confluence. All experiments used cells within a defined passage range, and cultures were routinely screened for mycoplasma (none were detected).

### HBLAK organotypic culture

For organotypic culture, 1 × 10^5^ HBLAK cells were seeded onto 0.4 µm polycarbonate membrane inserts (Millicell Cell Culture Inserts, 24-Well Hanging Inserts, 0.4 µm PET, Millipore). Cells were expanded to confluence and then differentiated in CnT-Prime 3D barrier medium (CELLnTEC) and 1:1 ratio of male and female urine. Barrier formation and epithelial coverage were verified in representative inserts by phalloidin and DAPI staining followed by fluorescence microscopy before downstream infection or treatment studies.

### BKPyV infection

For monolayer infection experiments, 1 × 10^5^ HBLAK cells were seeded in 12-well plates 24 h before infection. Cells were mock infected or infected with BKPyV Rearranged (Gardner) or Archetype strain at a multiplicity of infection (MOI) of 0.5. Virus was allowed to adsorb for 1 h at 4°C with gentle rocking in every 15 minutes, after which inoculum was removed, and cultures were maintained in fresh medium for the indicated postinfection intervals. Where indicated, cells were pretreated with chemotherapeutic, immune-conditioning, antiviral, or inhibitor compounds before infection.

In organotypic system, after differentiation, 0.5 MOI BKPyV (Archetype stain) was performed in the urine surface of the PET inserts either as pre-infection or post-infection as indicated.

### Genomic DNA isolation

Genomic DNA was isolated at indicated days of post-infection. Culture medium was aspirated and cells were lysed directly in the well with 150 µL tissue lysis solution consisting of 125 µL Buffer ATL (Qiagen) and 25 µL proteinase K (Qiagen). After 5 min incubation at room temperature, lysates were transferred to microcentrifuge tubes and incubated for 5 min at 65°C. DNA binding was achieved by adding 490 µL binding buffer PM together with 10 µL 3 M sodium acetate and loading the mixture onto spin columns (Zymo-Spin columns). Columns were washed once with 750 µL Buffer PE and once with 750 µL 80% ethanol, then dried by centrifugation and eluted in 50 µL preheated 10% Buffer EB (1 mM Tris-Cl, pH 8.5). After that the concentrations were estimated using Qubit (Thermo Fisher Scientific) and nanodrop.

### BKPyV genome copy quantification by qPCR

BKPyV genome copies were quantified by quantitative PCR using viral T-antigen primer sets [36]. Viral DNA levels were normalized to human mitochondrial DNA.

### Drug treatments and viability assays

To establish sublethal treatment conditions, HBLAK cells were exposed to serial dilutions of the indicated therapeutic agents, control compounds, or mechanistic inhibitors. Cell viability was quantified using CellTiter-Glo 2.0 (Promega) according to the manufacturer’s instructions. Luminescence values were normalized to the corresponding untreated or vehicle-treated controls, defined as 100% viability, and dose-response profiles were generated from replicate measurements.

For chemotherapeutic, immune-conditioning, and antiviral agents used to establish the therapy-associated BKPyV phenotype, IC_10_ and IC_20_ concentrations were estimated from the corresponding viability curves and used where indicated. For mechanistic and control compounds, working concentrations were selected from sublethal regions of the corresponding dose-response profiles rather than being restricted to a single calculated IC value. AU-15330 and FHD-286 were used at 100–300 nM in the BKPyV replication experiments, concentrations associated with approximately 89% and 93% viability, respectively, across the working range. Paclitaxel was used at 2.5–5 nM, concentrations falling within a minimal toxicity range.

### Cell cycle analysis

For cell-cycle analysis, cells were harvested, fixed according to the indicated experimental protocol, and washed thoroughly to remove residual fixative. Samples were adjusted to 1 × 10^6^ cells/mL in PBS and 1 µL FxCycle Violet Stain (Thermo Fisher Scientific; cat. no. F10347) was added per 1 mL of cell suspension. Samples were incubated for 30 min at room temperature protected from light and analyzed without washing by flow cytometry using manufacturer’s instructions.

### Fluorescence microscopy

Fluorescence images were acquired using a BioTek Cytation 9 Cell Imaging Multimode Reader (Agilent). For endpoint imaging, samples were blocked and then stained with the T antigen and VP1 primary antibodies, followed by alexa fluor-conjugated secondary antibodies, counterstained with DAPI, and imaged using specific channels, and matched exposure settings across experimental conditions. Image analysis was performed in FIJI, with identical processing parameters applied to all samples within an experiment.

### Confocal microscopy

For confocal imaging, HBLAK-derived organotypic cultures were used. At the indicated time points, the organotypic tissue was gently removed from the insert membrane, transferred to glass microscope slides (Fisher Scientific), and fixed. Tissues were then permeabilized, blocked, and incubated overnight with primary antibodies against BKPyV T antigen and VP1 together with an F-actin. After washing, samples were incubated with Alexa Fluor-conjugated secondary antibodies (AF488, AF555, and AF647), counterstained with DAPI, mounted under glass coverslips using mounting medium, and imaged on a Zeiss LSM 880 Airyscan confocal microscope using identical acquisition settings within each experiment.

Z-stack image series were processed in FIJI (ImageJ). Channel display settings were applied identically across matched experimental groups, and representative images were shown in the figure.

### Low-molecular-weight DNA and Plasmid-Safe assay

Low-molecular-weight DNA was isolated using a modified HIRT lysis protocol as described in [37], and DNA was amplified where indicated using TempliPhi (Cytiva) according to the manufacturer’s instructions. To enrich for circular episomal DNA, Hirt DNA was treated with Plasmid-Safe ATP-dependent DNase in 1X Plasmid-Safe reaction buffer supplemented with 1 mM ATP. Reactions were incubated at 37°C for 30 min and heat-inactivated at 70°C for 30 min before downstream qPCR. Untreated and Plasmid-Safe-treated aliquots were analyzed in parallel to distinguish circular from linear viral DNA species.

### Cell lysis and Western blotting

HBLAK protein lysates for Western blot were harvested by directly lysing 1 × 10^6^ pelleted cells in 100 µL Laemmli reducing sample buffer. Samples were boiled for 10 min, and 10 µL of each lysate was resolved on a NuPAGE 4–12% Bis-Tris gel (Thermo Fisher Scientific) and transferred to nitrocellulose membranes using standard procedures. Membranes were blocked, incubated with primary antibodies overnight at 4°C, and then probed with the appropriate HRP-conjugated secondary antibodies. Blots were imaged using a GE Amersham Imager 480.

### Oxford Nanopore long-read sequencing and structural variant analysis

For long-read analysis of BKPyV structural variation, DNA from infected cell cultures and patient urine specimens was prepared using the Oxford Nanopore sequencing kit for rolling circle amplification products according to the manufacturer’s instructions and sequenced on a MinION R10 flow cell.

### Viral DNA short-read sequencing and variant analysis

Demultiplexed FASTQ files were obtained from the NCI Genomics Core after sequencing indexed viral DNA libraries from BKPyV-infected HBLAK samples under treatment conditions and from longitudinal urine specimens on an Illumina NextSeq instrument using paired-end 2×150-bp reads. Raw reads were quality filtered and adapter trimmed with fastp. A Bowtie2 index was generated from the BKPyV Gardner reference genome, and quality-controlled reads were aligned to the reference using bowtie2 in --very-sensitive-local mode. Alignments were coordinate-sorted and indexed with samtools. Single-nucleotide variants and short indels were called using LoFreq, and the resulting files were annotated with SnpEff.

### RNA sequencing and differential gene expression analysis

Total RNA was isolated using RNeasy mini kit (Qiagen) according to the manufacturer’s instructions. RNA integrity was assessed using TapeStation, and libraries were prepared. Following library preparation, indexed RNA-seq libraries were submitted to the NCI Genomics Core and sequenced on an Illumina NextSeq platform using paired-end 2×150-bp reads. FASTQ files were quality filtered and adapter trimmed with fastp, then aligned to the reference genome using STAR. Gene-level count files generated by STAR were merged across samples to create a combined count matrix for downstream analysis. Differential expression analysis was performed in R using DESeq2. Data visualization was carried out with ggplot2, and overlaps among gene sets were displayed using UpSet plots. Pathway enrichment analysis was performed using IPA (Qiagen).

### ATAC sequencing and motif enrichment analysis

Chromatin accessibility profiling was performed using the ATAC-Seq Kit (Active Motif) according to the manufacturer’s instructions. The kit contains the assembled transposome, tagmentation buffer, PCR reagents, indexed primers, and bead-based purification reagents, and is designed for low-input chromatin accessibility workflows. Following library preparation, indexed ATAC-seq libraries were sequenced at the NCI Genomics Core on an Illumina NextSeq platform.

ATAC-seq data were processed on NIH Biowulf using the ENCODE ATAC-seq pipeline, which is designed for automated end-to-end quality control and processing of ATAC-seq and is configured through a JSON input file. The pipeline was run using paired-end FASTQ inputs and the appropriate reference-genome resources available on Biowulf. Differentially accessible regions and track files generated by the pipeline were used for downstream analysis. Motif enrichment within treatment-associated accessible regions was assessed using HOMER. The signal on the BKPyV genome was normalized against the total mapping reads to the early region which had the least variability in binding signal across all conditions. Transcription factor motif binding prediction was performed using TFBIND and plotted with ggplot2.

### Cut & Run assay

Protein–chromatin occupancy was profiled using the ChIC/CUT&RUN Assay Kit (Active Motif) according to the manufacturer’s instructions. Libraries generated from CUT&RUN DNA were submitted to the NCI Genomics Core and sequenced on an Illumina NextSeq platform using paired-end 2×100-bp reads.

Sequencing data were analyzed on NIH Biowulf using CUT&RUNTools2, a pipeline designed for both CUT&RUN datasets. The workflow uses a JSON configuration file and integrates standard tools including bowtie2, samtools, and MACS2, with downstream options for motif analysis and footprint-style interpretation. The signal on the BKPyV genome was normalized against the total mapping reads to the early region which had the least variability in binding signal across all conditions.

### Statistical analysis

Data were assessed for statistical significance using Prism v10 software (GraphPad). On all graphs, statistical P or value significance is represented as follows: *<0.05, **<0.01, ***<0.001, and ****<0.0001. The abbreviation “ns” denotes that an effect was not statistically significant.

## Data availability

All raw sequencing data will be available in SRA. Full resolution microscopy images will be available in FigShare.

## Supplementary figure legends

**Supplementary Figure 1. Rearranged and archetype BKPyV replication in HBLAK monolayers.**

(A) BKPyV genome abundance at 3 and 5 dpi after infection with rearranged or archetype Dik BKPyV at MOI 0.5. (B) Archetype BKPyV genome abundance at 5 dpi following infection at the indicated MOI. Data represent the mean ± SD from three biological replicates.

**Supplementary Figure 2. Determination of sublethal treatment concentrations in HBLAK cells.**

Dose-response viability curves for (A) cyclophosphamide, (B) fludarabine, (C) busulfan, (D) etoposide, (E) cisplatin, (F)5-fluorouracil, (G) ganciclovir, (H) acyclovir, and (I) cidofovir. Dashed lines indicate the viability thresholds used to define IC_10_ and IC_20_. Values represent the mean ± SD from three experiments.

**Supplementary Figure 3. Validation of treatment-associated BKPyV enhancement across exposure conditions and epithelial models.**

(A) Cell-cycle distribution in mock-infected, archetype-infected, or rearranged-BKPyV-infected HBLAK cells treated with vehicle or etoposide IC_10_. (B) BKPyV genome abundance following pre-infection or post-infection treatment with etoposide or cisplatin. (C) BKPyV genome abundance after IC_20_ pretreatment and low-MOI infection (0.1 MOI). (D) Representative western blot analysis of cellular stress and DNA damage-response proteins in mock and BKPyV-infected HBLAK cells following the indicated therapeutic exposures. BAX, phospho-Chk1, phospho-Chk2, BCL-XL, total Chk1, ATM, and ATR were examined, with beta-actin shown as a loading control. (E) Dose-dependent effect of paclitaxel on HBLAK cell viability measured using CellTiter-Glo 2.0. Viability was normalized to untreated controls, defined as 100%; the estimated IC_10_ and IC_20_ positions are indicated. (F) BKPyV genome abundance following short (48 h) or extended (5-day) exposure to cyclophosphamide or fludarabine in rearranged-BKPyV-infected HBLAK cells. (G) BKPyV genome abundance following short (48 h) or extended (5-day) etoposide exposure in rearranged-BKPyV-infected HBLAK cells. (H) BKPyV genome abundance following exposure to etoposide at IC_10_, IC_20_, or IC_30_ concentrations. (I) BKPyV genome abundance in renal proximal tubular epithelial cells (RPTECs) following exposure to the indicated immune-conditioning agents, chemotherapeutics, or antivirals. Data represent the mean ± SD from three experiments.

**Supplementary Figure 4. Post-infection therapeutic exposure enhances archetype BKPyV replication in organotypic urothelial cultures.**

(A) Experimental workflow for post-infection treatment of differentiated HBLAK rafts.

(B) Representative confocal images of archetype-BKPyV-infected rafts exposed to the indicated treatment after infection and stained for TAg, VP1, F-actin, and DAPI.

**Supplementary Figure 5. Therapeutic exposure does not produce recurrent BKPyV structural variants or mutational signatures in HBLAK cells.**

(A) Frequencies of duplications, insertions, inversions, and microhomology-associated events detected by Nanopore long-read sequencing across treatment conditions. (B) Trinucleotide-context substitution profiles derived from short-read BKPyV sequencing.

**Supplementary Figure 6. Urinary BKPyV loads in independent transplant cohorts.**

(A) Urinary BKPyV loads measured by clinical qPCR in eight transplant recipients enrolled at the NIH Clinical Center. Plotted value represents the highest measured value (I guess!) [Need to add more information?] (B) Longitudinal urinary BKPyV loads at baseline, transplantation/day 0, and approximately day 14 in the independent multicenter cohort. Each line represents one patient with available paired measurements. Viral loads are shown on a log_10_ scale. [More information?]

**Supplementary Figure 7. Longitudinal BKPyV substitution spectra in the NIH Clinical Center cohort.**

Trinucleotide-context substitution profiles for BKPyV sequences recovered from longitudinal urine samples from each NIH Clinical Center participant. Each panel represents one patient, and sampling times are shown relative to transplantation. No recurrent time-dependent substitution pattern was observed across participants.

**Supplementary Figure 8. Baseline-subtracted BKPyV substitution spectra in the multicenter hemorrhagic cystitis cohort.**

Longitudinal trinucleotide-context substitution profiles for BKPyV sequences recovered from the multicenter cohort. For each patient, substitution frequencies at later time points were compared with or subtracted from the earliest available baseline sample indicated in the panel. No recurrent treatment-associated mutational signature was shared across patients.

**Supplementary Figure 9. Longitudinal BKPyV structural-variant profiles in transplant recipients.**

Frequencies of duplications, insertions, inversions, and microhomology-associated structural events detected by long-read sequencing in longitudinal urine specimens. Panels represent individual patients and collection times relative to transplantation. No reproducible structural-variant trajectory was observed across participants.

**Supplementary Figure 10. Therapeutic exposure does not induce a reproducible accumulation of linear BKPyV DNA species.**

(A) Schematic of the Plasmid-Safe ATP-dependent DNase assay. Total DNA was analyzed without enzyme, after Plasmid-Safe treatment, or after linearization with SphI followed by Plasmid-Safe treatment. (B) Estimated Plasmid-Safe-resistant circular and Plasmid-Safe-sensitive linear BKPyV DNA fractions in infected-control, cyclophosphamide-treated, etoposide-treated, and cidofovir-treated cells. The assay provides an estimate of genome topology. Data represent the mean ± SD from two experiments.

**Supplementary Figure 11. Shared host differentially expressed genes within therapeutic classes.**

Principal-component analysis of normalized host RNA-seq counts from mock and rearranged BKPyV-infected HBLAK cells exposed to chemotherapeutics (A), immune-conditioning (C), and anti-virals (E). UpSet plots showing intersections of differentially expressed host genes across (B) chemotherapeutic, (D) immune-conditioning, and (F) antiviral treatment comparisons in mock and BKPyV-infected HBLAK cells.

**Supplementary Figure 12. Integration of host RNA-seq and ATAC-seq identifies a shared treatment-responsive gene set.**

UpSet plot of genes that were both differentially expressed by RNA-seq and associated with treatment-enriched promoter accessibility by ATAC-seq. The common intersection contains 78 genes.

**Supplementary Figure 13. Cell viability following SWI/SNF-directed pharmacologic perturbation in HBLAK cells.**

HBLAK cells were exposed to increasing concentrations of AU-15330 (A) or FHD-286 (B), and cellular viability was measured using CellTiter-Glo 2.0. Luminescence values were normalized to untreated controls, defined as 100% viability. The 100–300 nM concentration range used for BKPyV replication experiments is indicated. Data represent the mean ± SD from three replicate wells per concentration. The

300-nM concentration was not directly measured in the viability assay and lies within the range bounded by the measured 200- and 500-nM concentrations.

## Supplementary table titles

**Supplementary Table 1. Sublethal concentrations of therapeutic agents used in HBLAK experiments.**

