## Supplementary figures for "Therapeutic-induced chromatin remodeling via AP-1/SWI-SNF enhances BK Polyomavirus replication in urothelial cells"

**Supplementary Figure 1.**

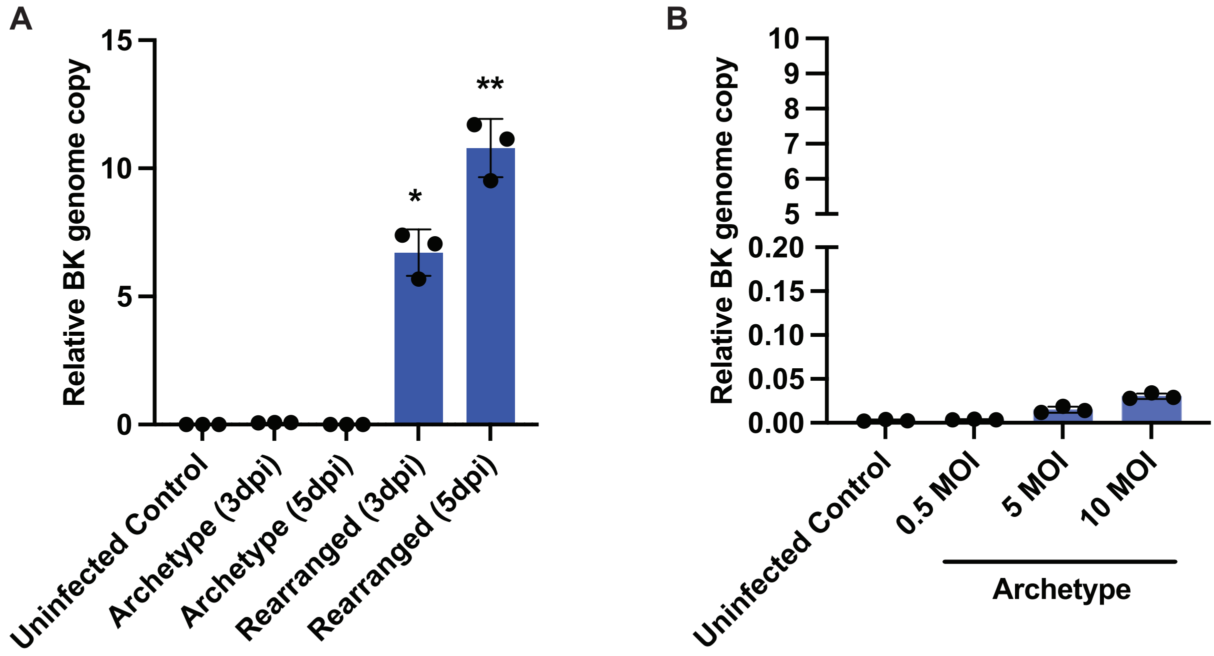

#### Supplementary Figure 2.

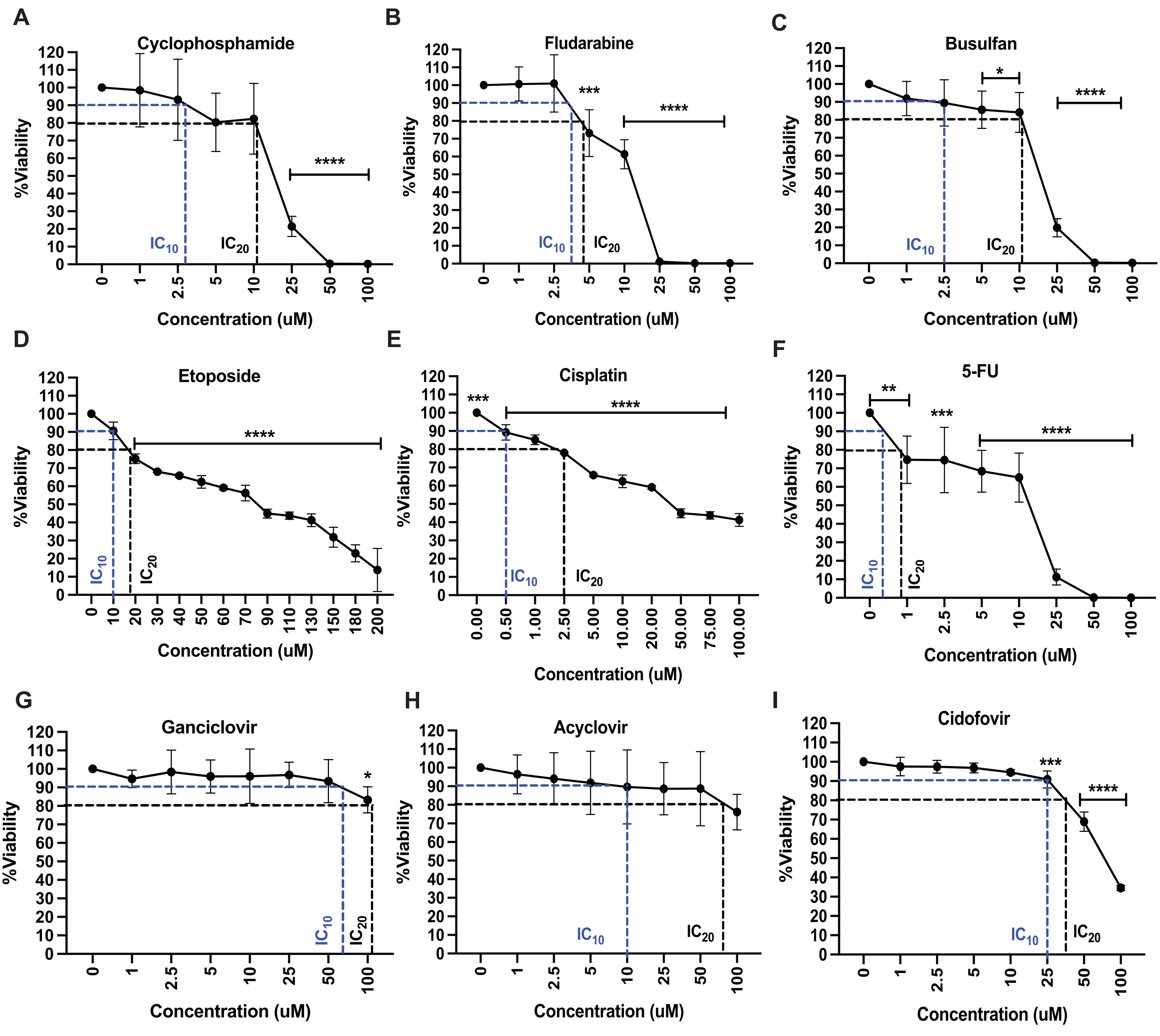

#### Supplementary Figure 3.

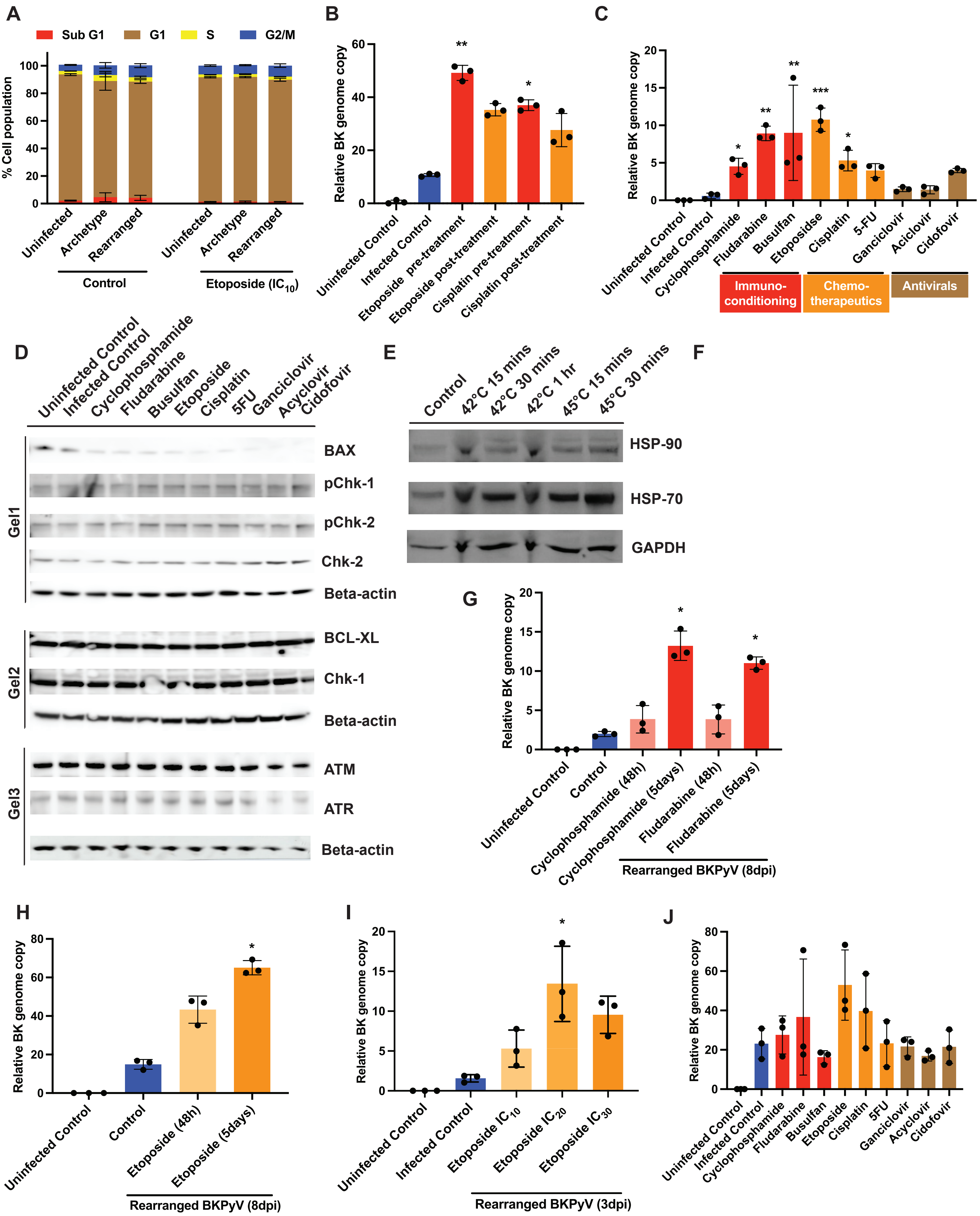

#### Supplementary Figure 4

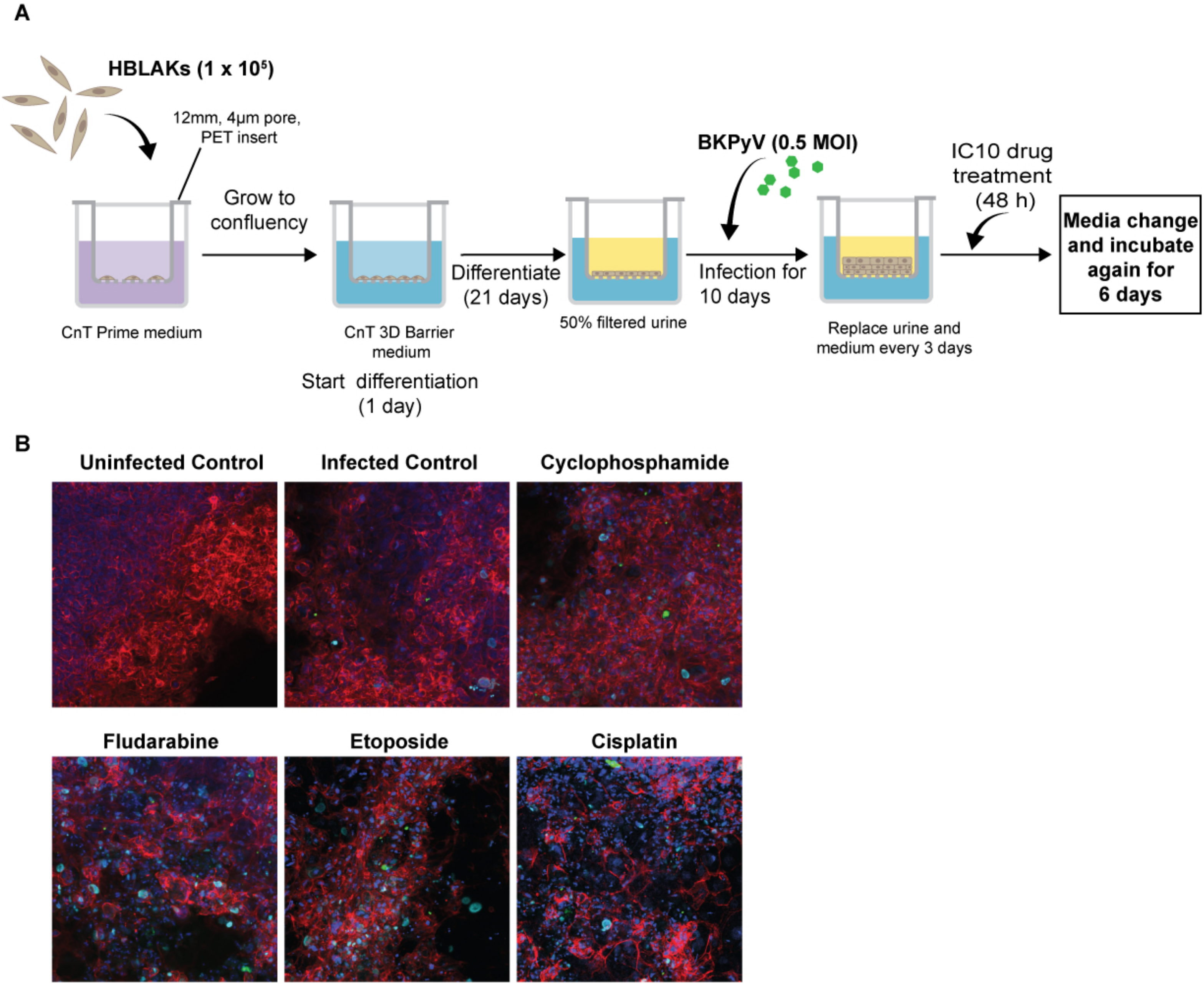

#### Supplementary Figure 5

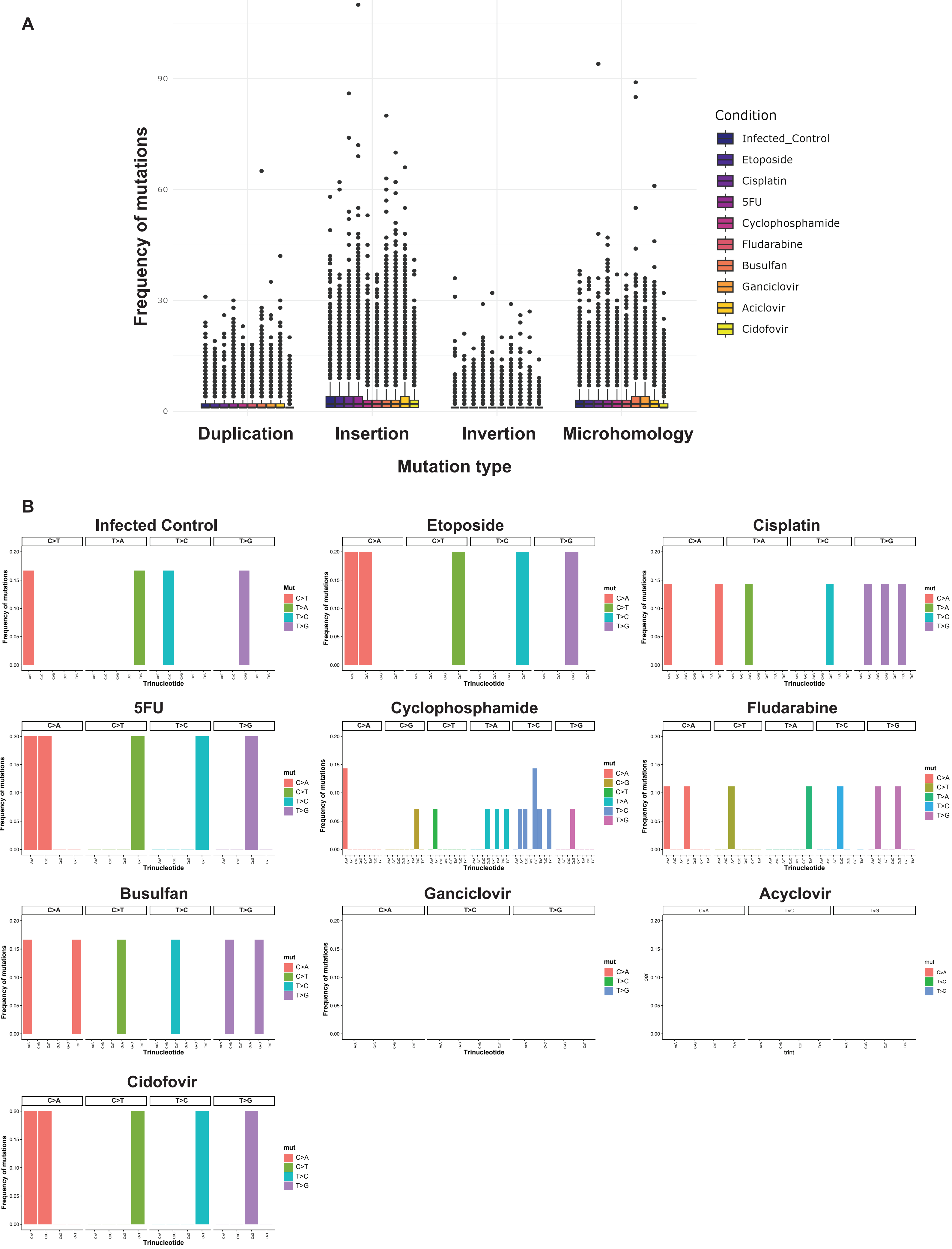

#### Supplementary Figure 6

**A**

**B**

### Supplementary Figure 7

#### Cohort-1

##### Patient 1 [Conditioning start Day -6, Transplant Day 0]

###### Day 0

###### Day +7

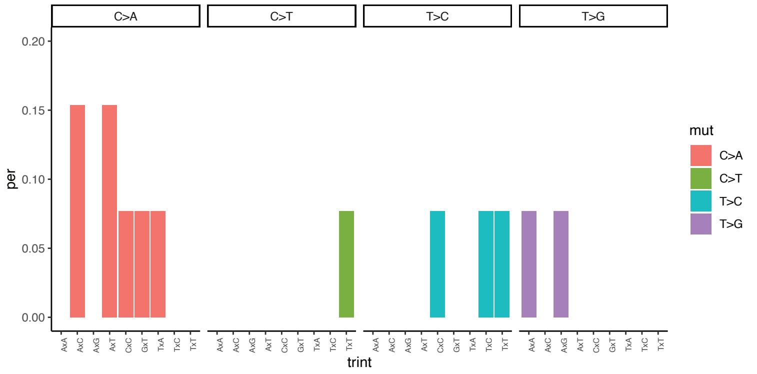


###### Day +14

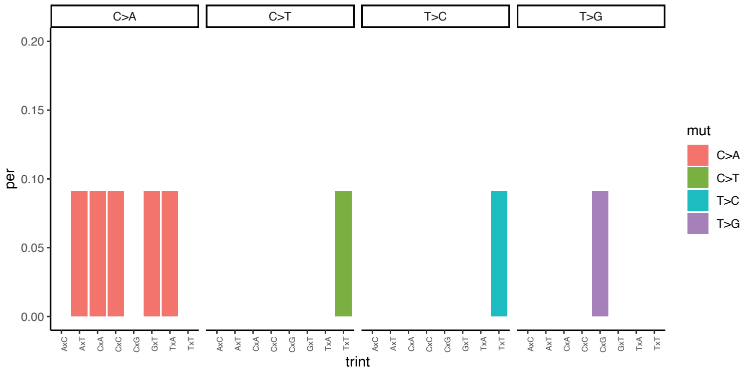


##### Patient 2 [Conditioning start Day -14, Transplant Day 0]

###### Day -14

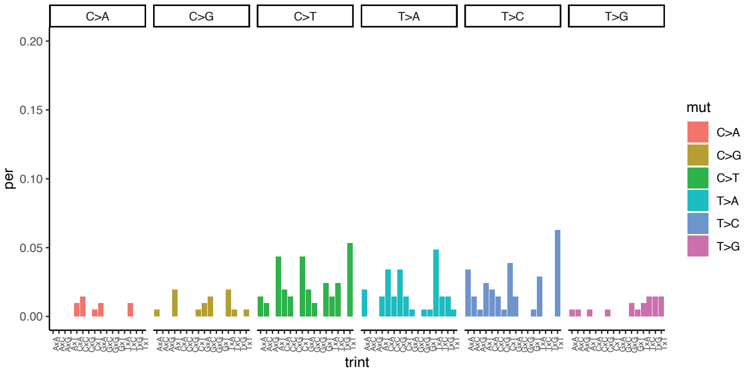


###### Day -7

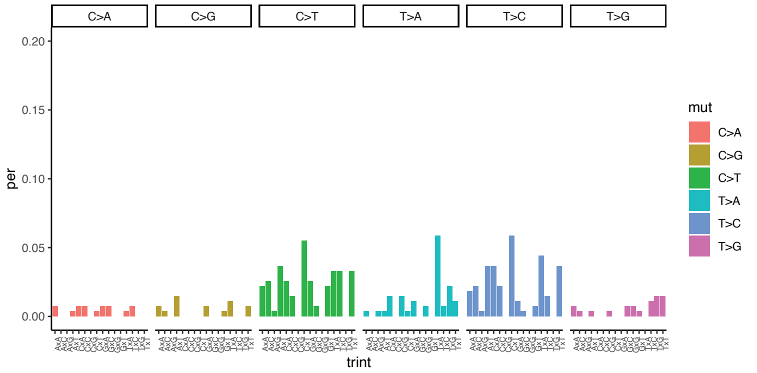


###### Day +28

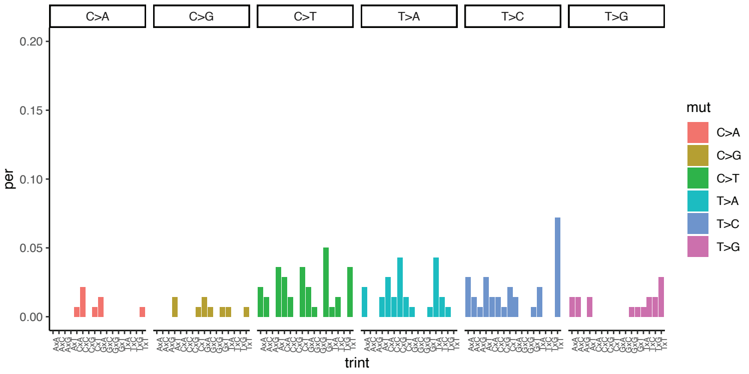


###### Day +35

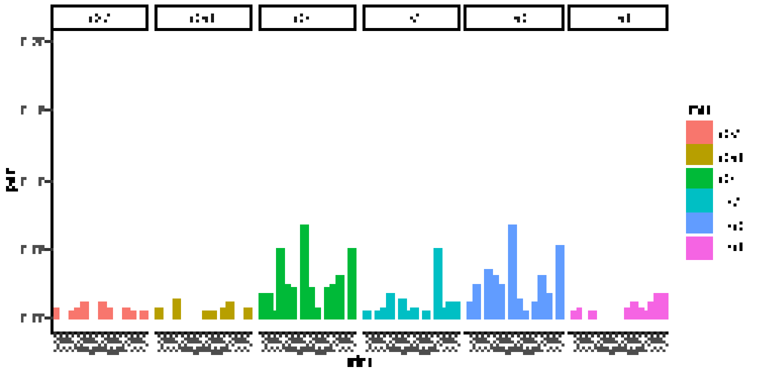

### Cohort-1

#### Patient 4 [Conditioning start Day -6, Transplant Day 0]

##### Day +41

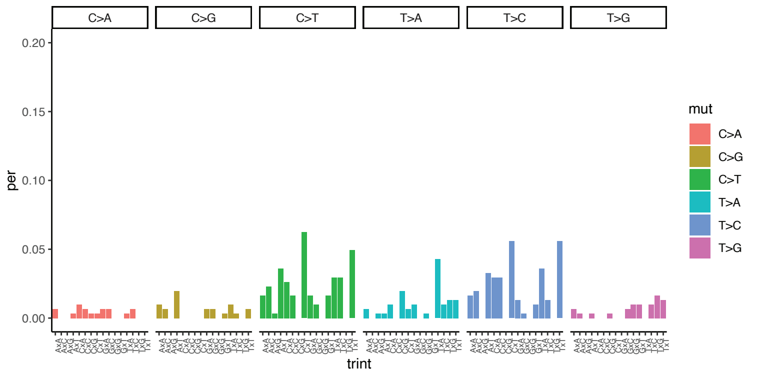


##### Day +62

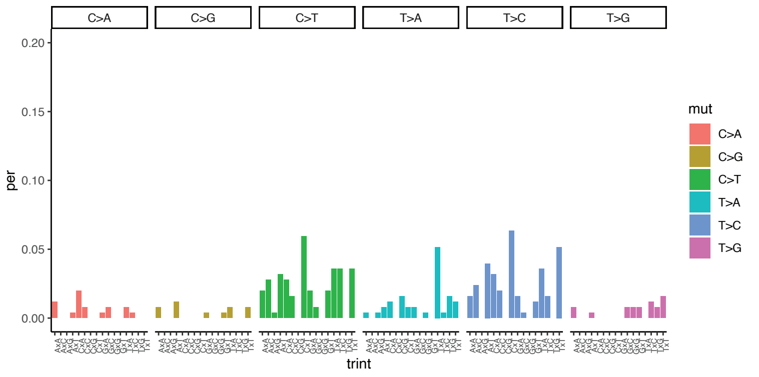


##### Day +69

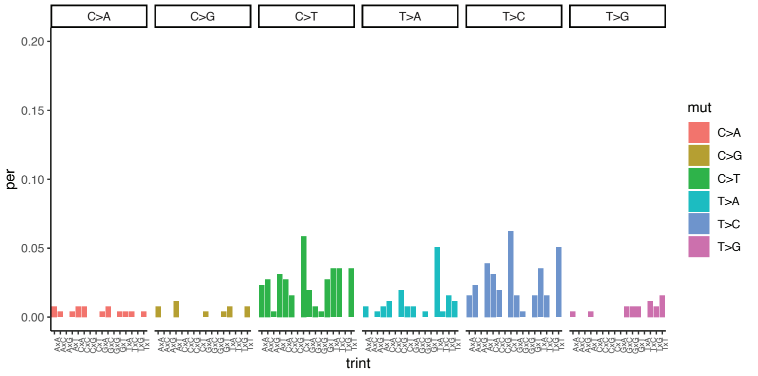


##### Day +76

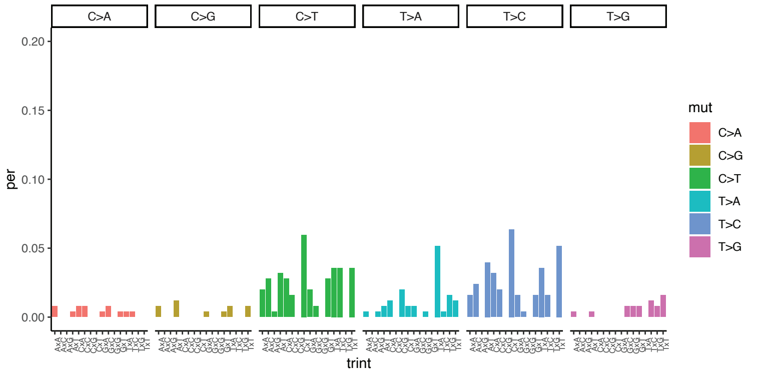


#### Patient 5 [Conditioning start Day -117 Transplant Day 0]

##### Day -97

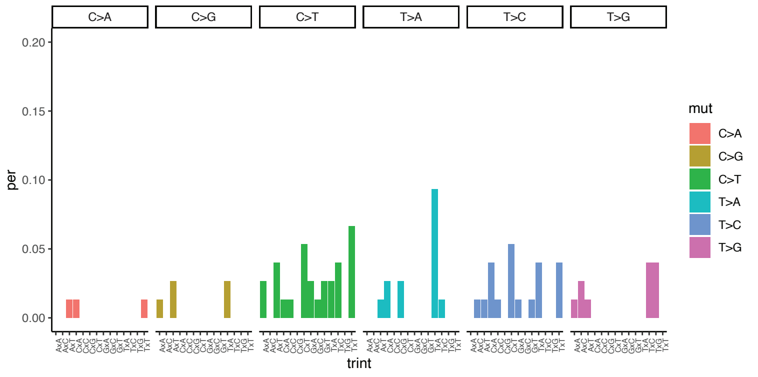


##### Day -90

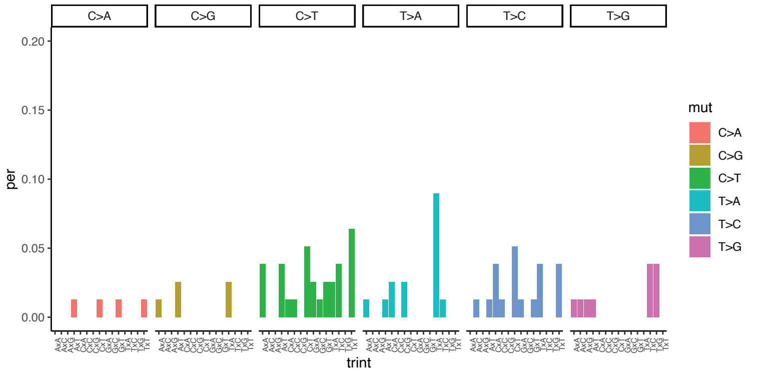


##### Day -84

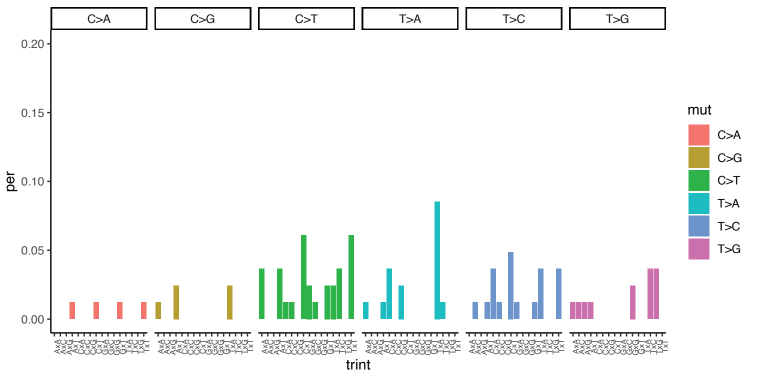


##### Day -77

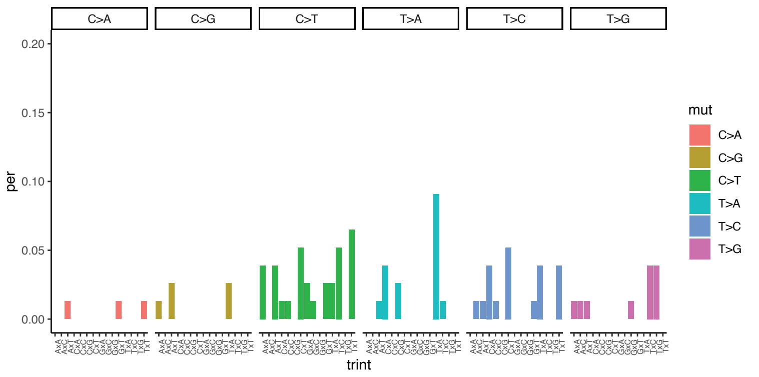

### Cohort-1

#### Patient 6 [Conditioning start Day -6, Transplant Day 0]

##### Day +63

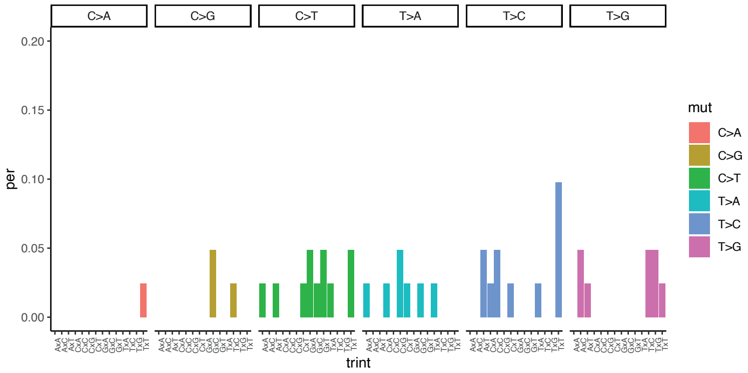


##### Day +70

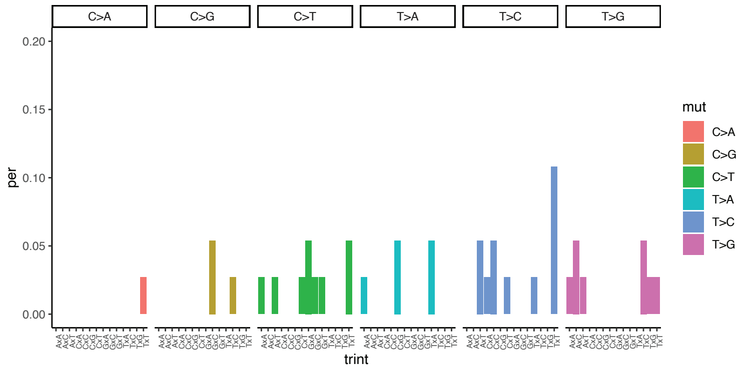


##### Day +77

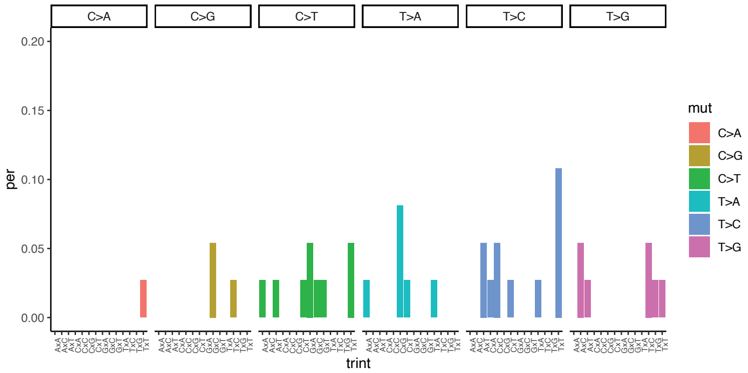


#### Patient 7 [Conditioning start Day -6 Transplant Day 0]

##### Day +34

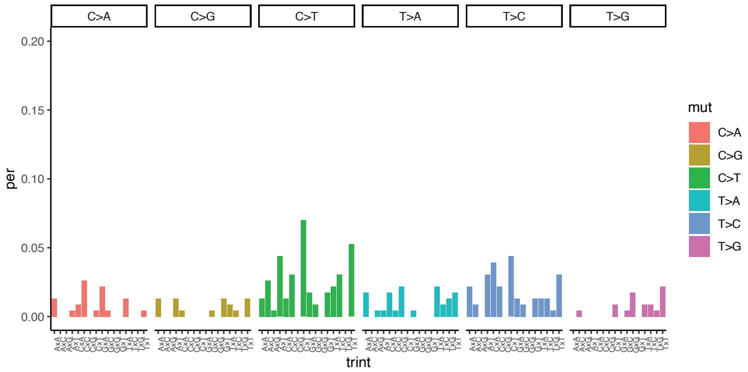


##### Day +41

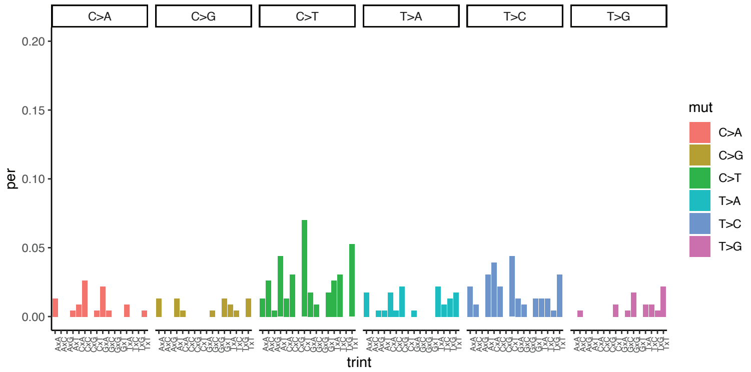


##### Day +48

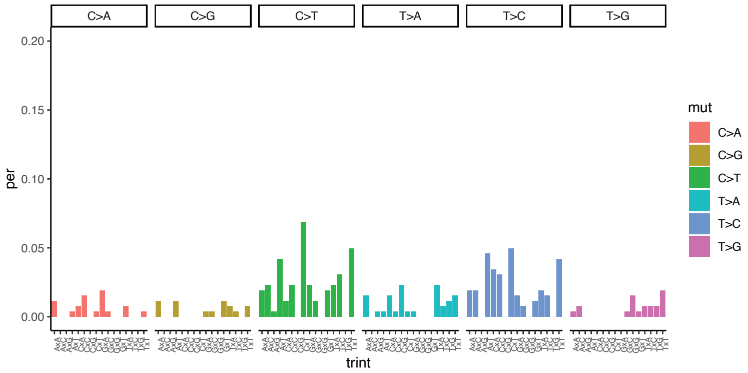


##### Day +55

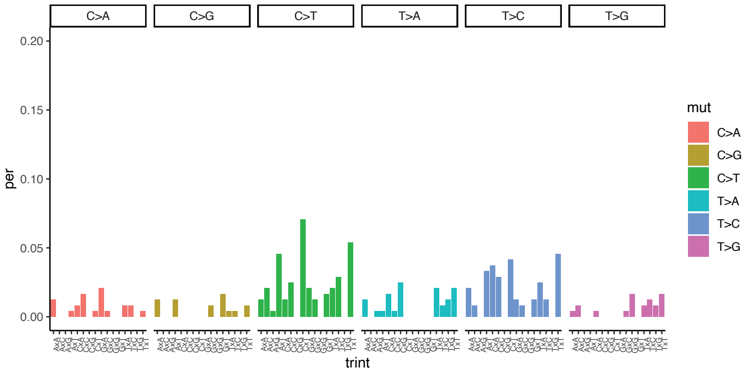

### Cohort-1

#### Patient 9 [Conditioning start Day -5 Transplant Day 0]

##### Day +14

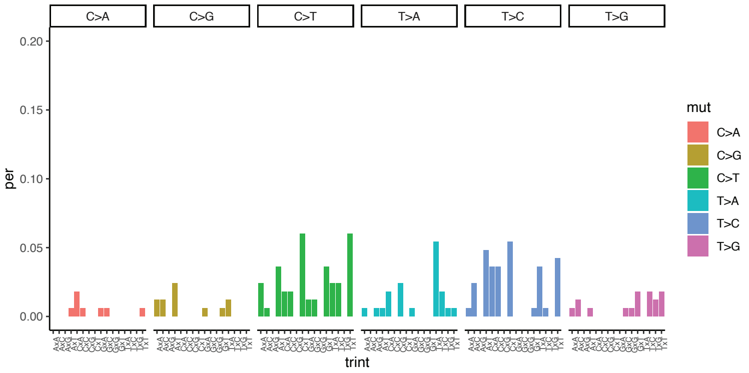


##### Day +21

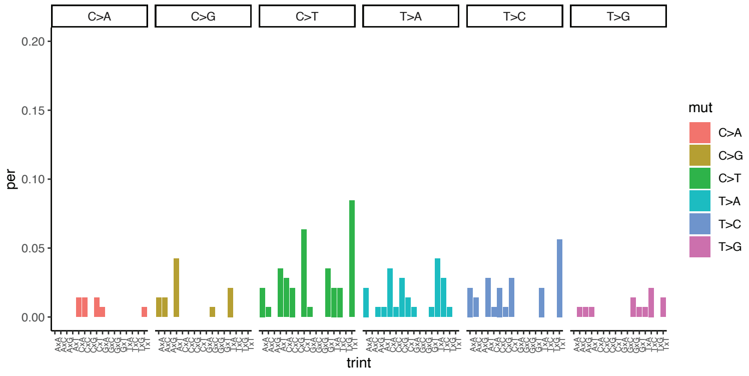


##### Day +28

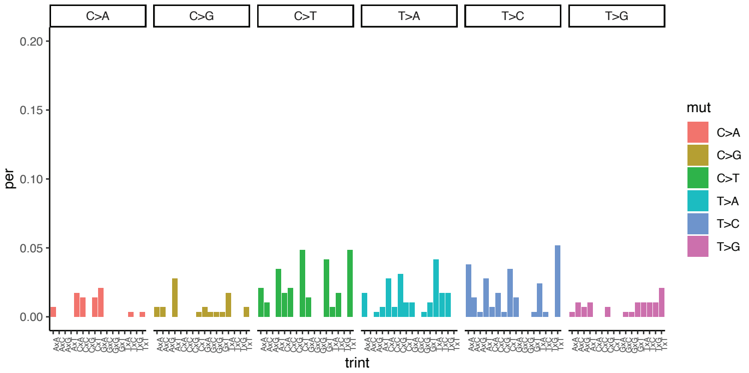


##### Day +35

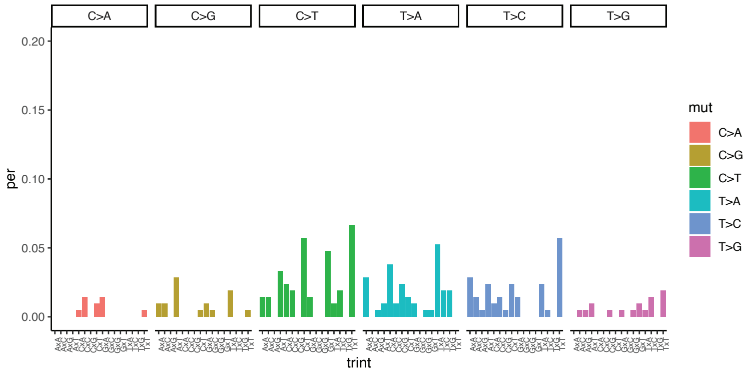


#### Patient 10 [Conditioning start Day -6 Transplant Day 0]

##### Day +34

##### Day +41

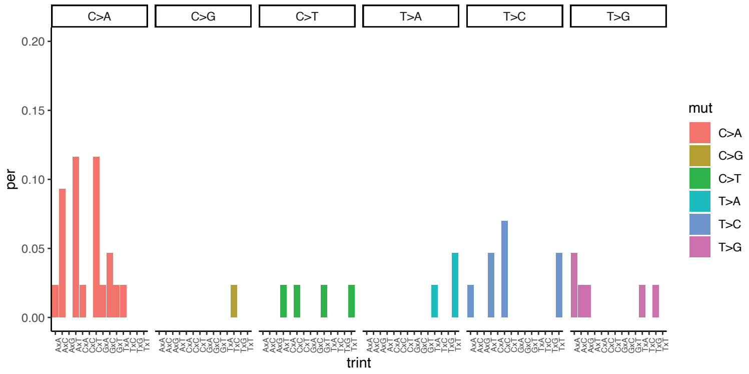


##### Day +55

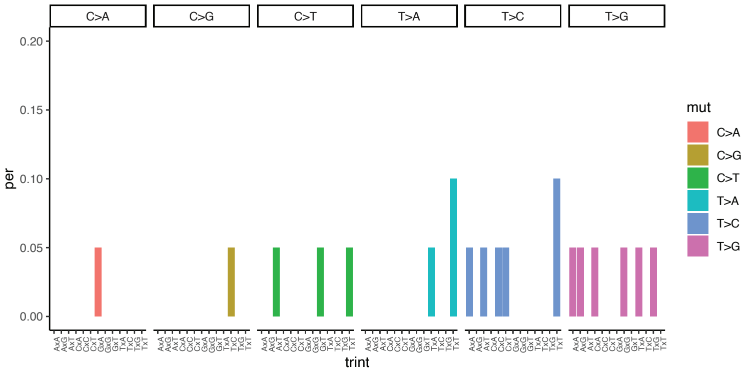

### Supplementary Figure 8

#### Cohort-2

##### Patient 1 (substracted from Day -16)

Day -2

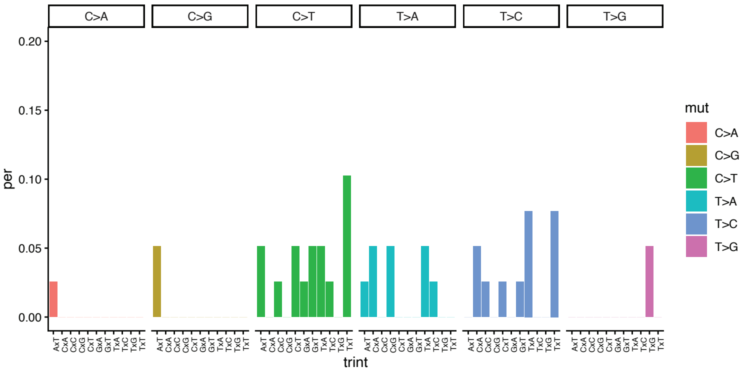


Day +6

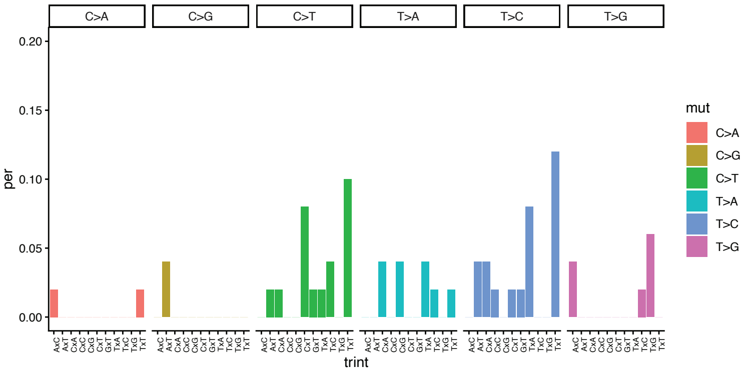


##### Patient 3 (substracted from Day -10)

(No mutation on Day -3 and +4)

Day +18

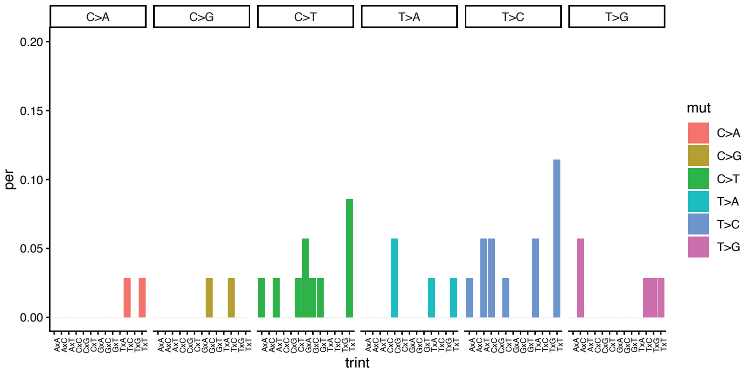


##### Patient 4 (substracted from Day -4)

Day +4

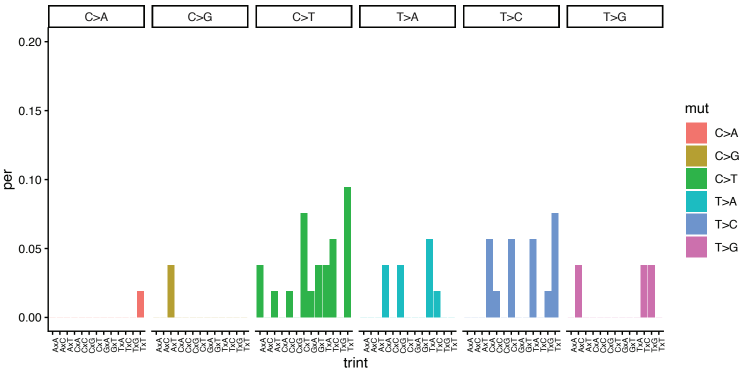


Day +11

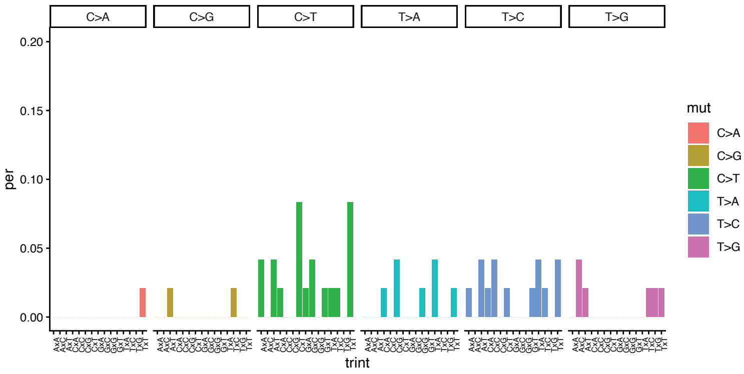

### Cohort-2

#### Patient 5 (substracted from Day -7)

##### Day +18

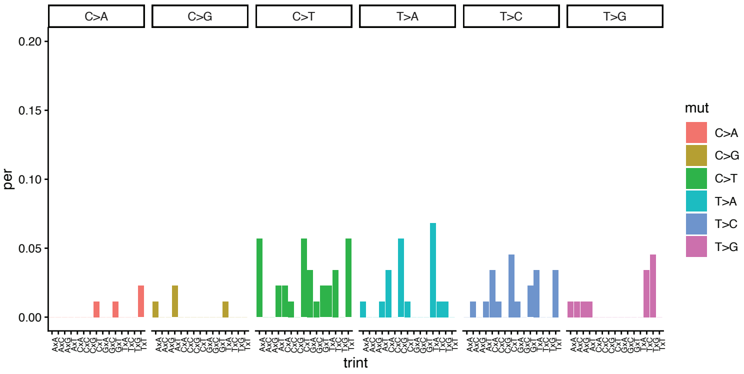


#### Patient 6 (substracted from Day -22)

##### Day -2

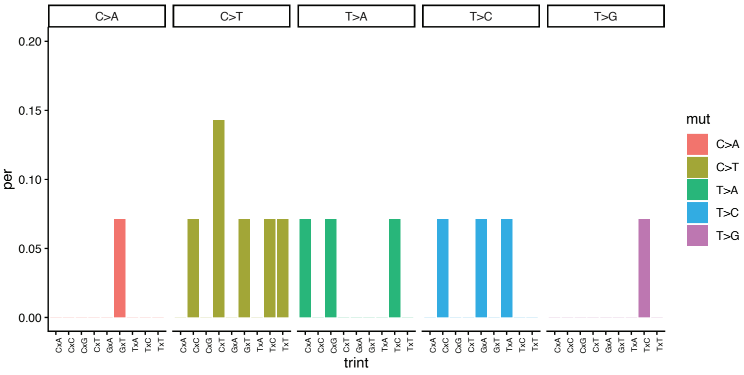


##### Day +5

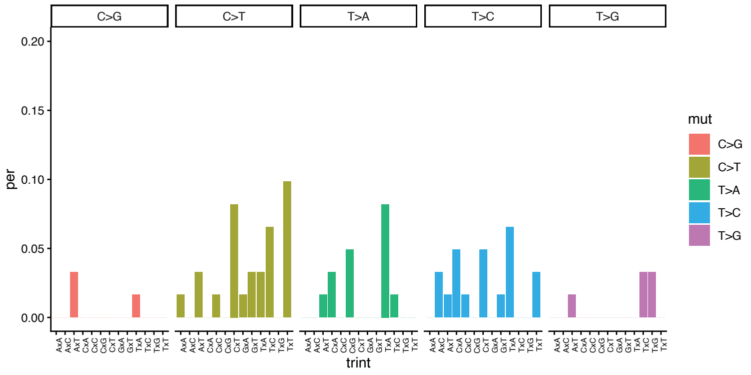


##### Day +18

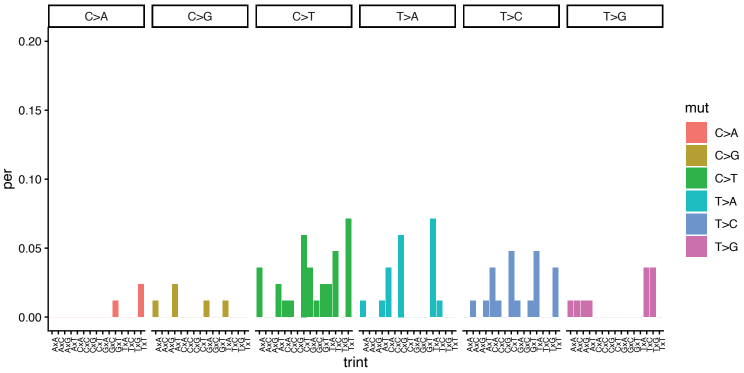


#### Patient 7 (substracted from Day -10)

##### Day +18

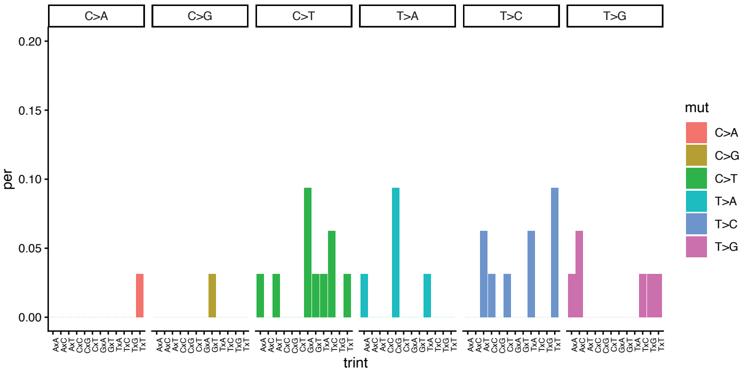

### Cohort-2

#### Patient 8 (substracted from Day -17)

##### Day -3

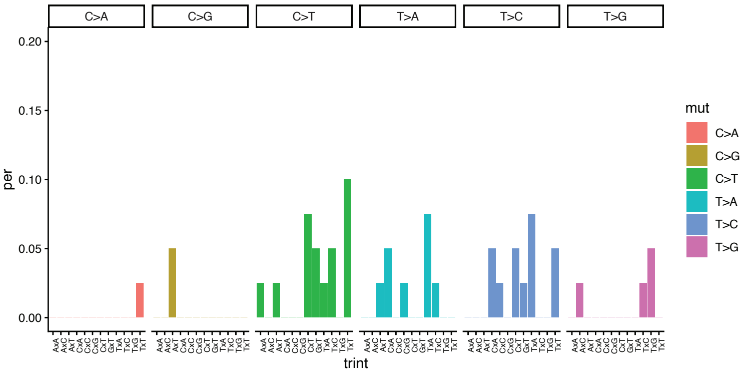


##### Day +4

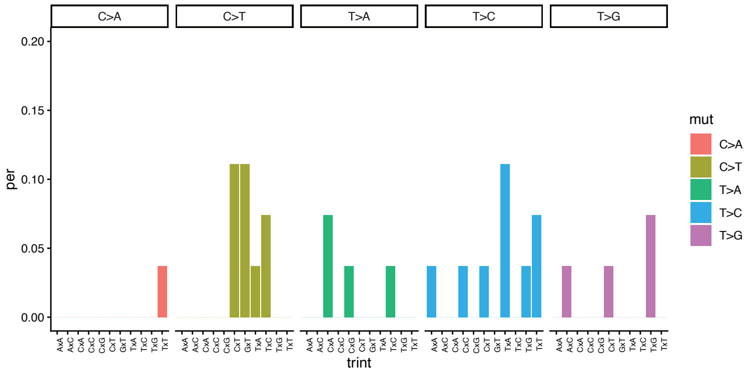


##### Day +11

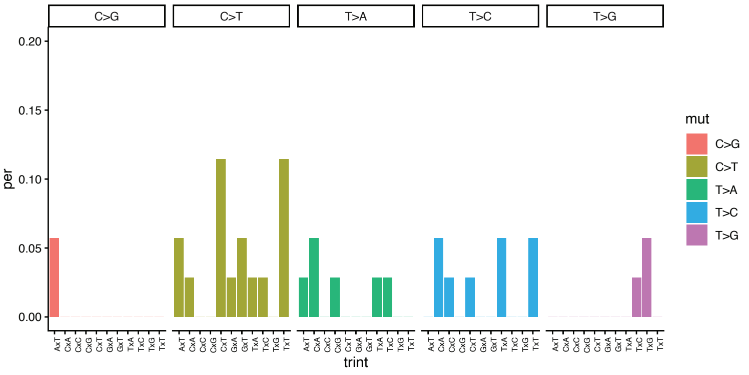


#### Patient 9 (substracted from Day -10)

##### Day -1

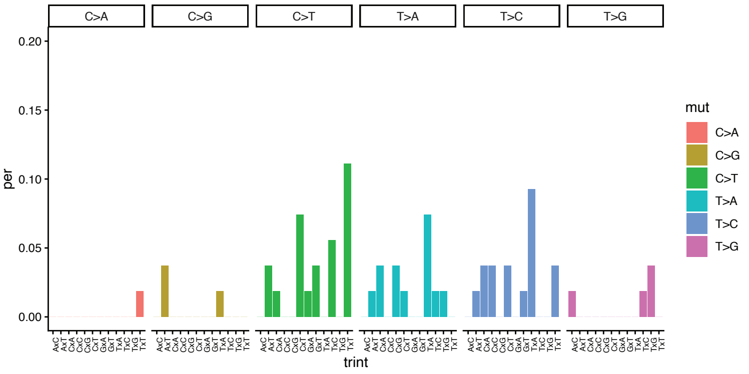


##### Day +6

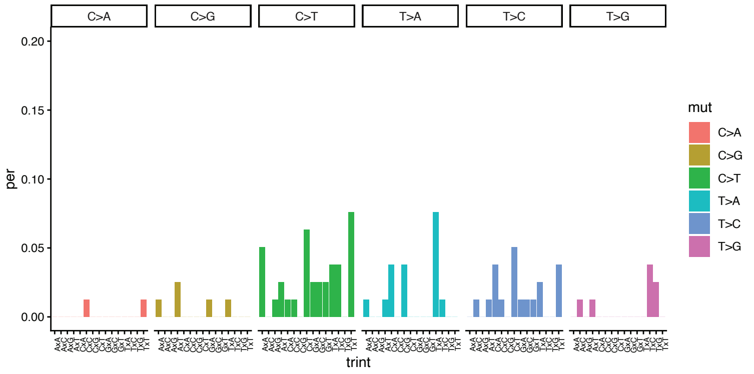


#### Patient 10 (substracted from Day -10)

##### Day +18

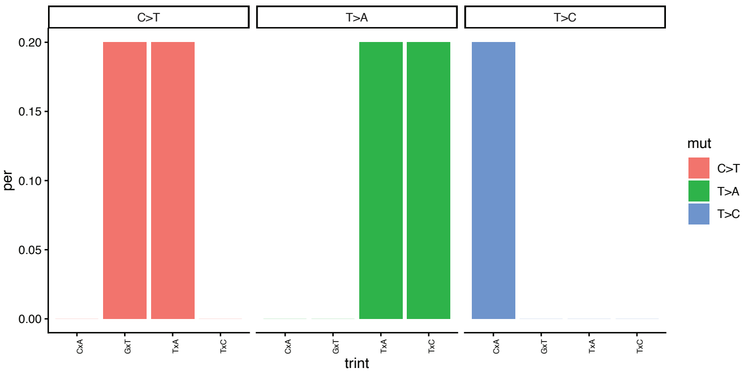

### Cohort-2

#### Patient 11 (substracted from Day -7)

##### Day +2

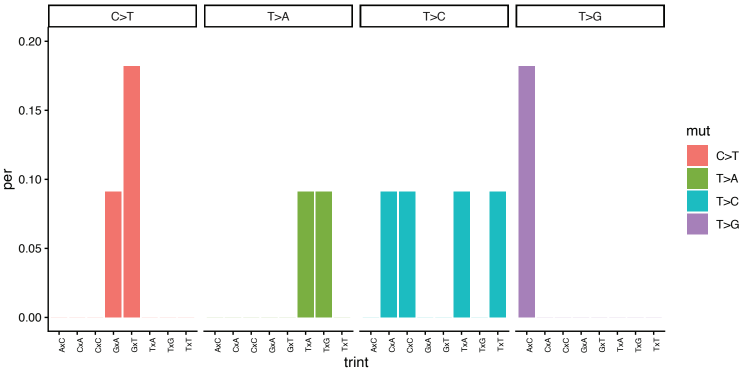


##### Day +7

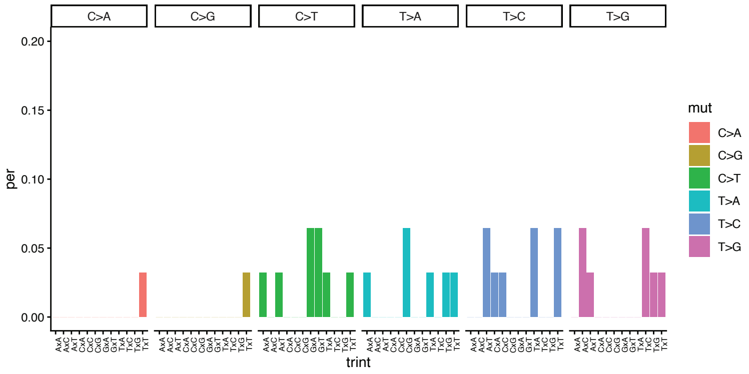


#### Patient 12 (substracted from Day -8)

##### Day +7

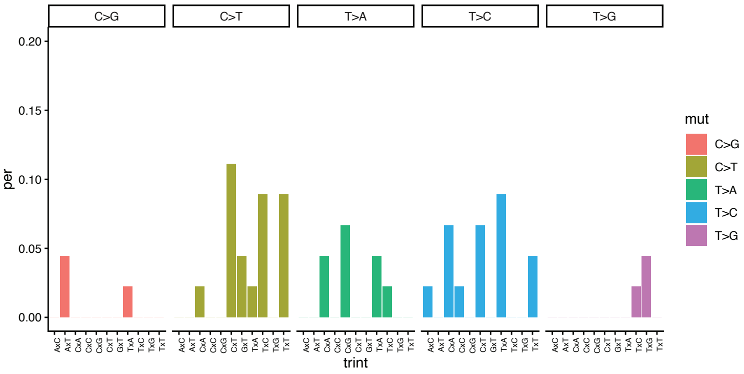


#### Patient 14 (substracted from Day -7)

##### Day -1

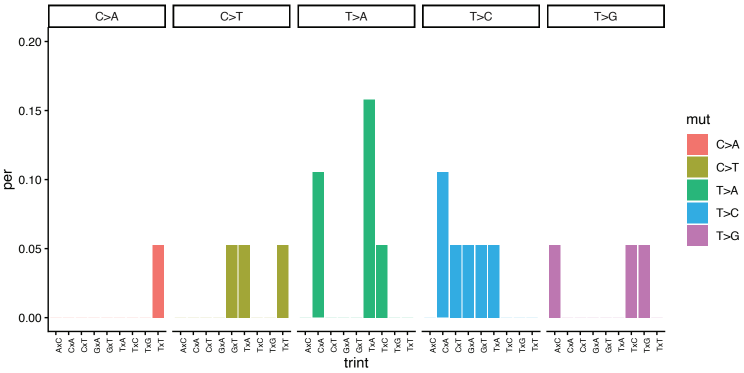


##### Day +6

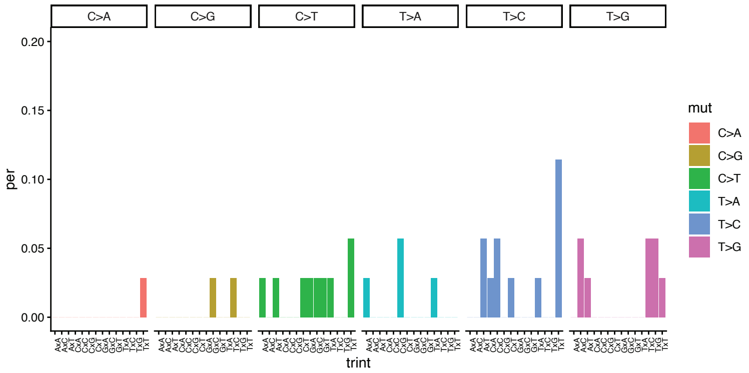

### Cohort-2

#### Patient 15 (substracted from Day -10)

##### Day-2

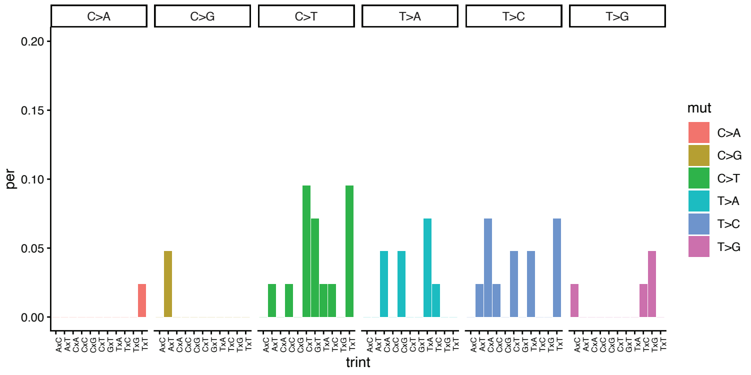


##### Day +5

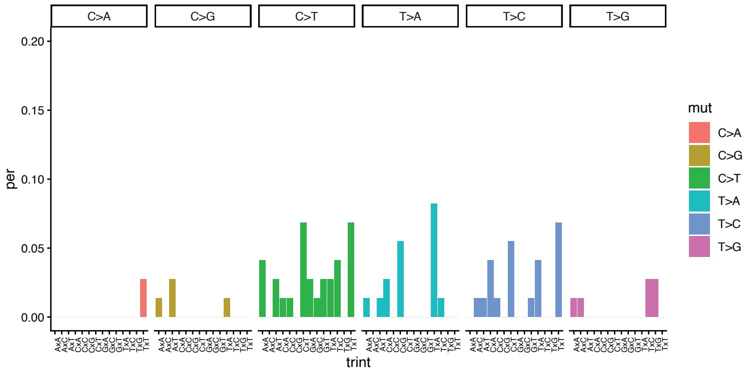


##### Day +12

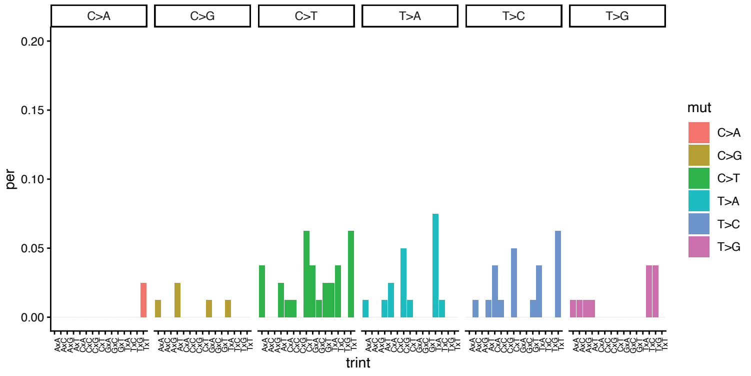


#### Patient 16 (substracted from Day -7)

##### Day +7

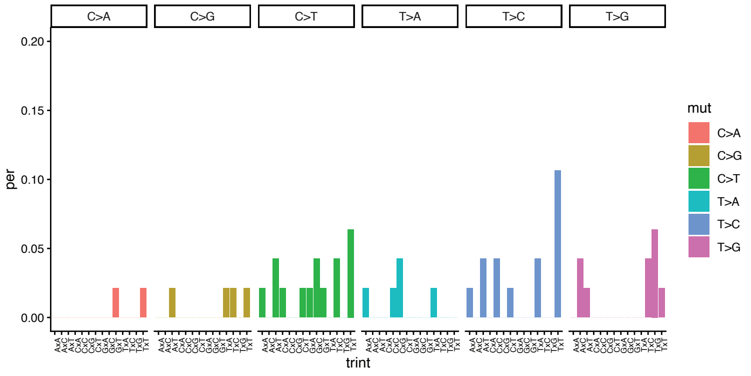


#### Patient 17 (substracted from Day -8)

##### Day +5

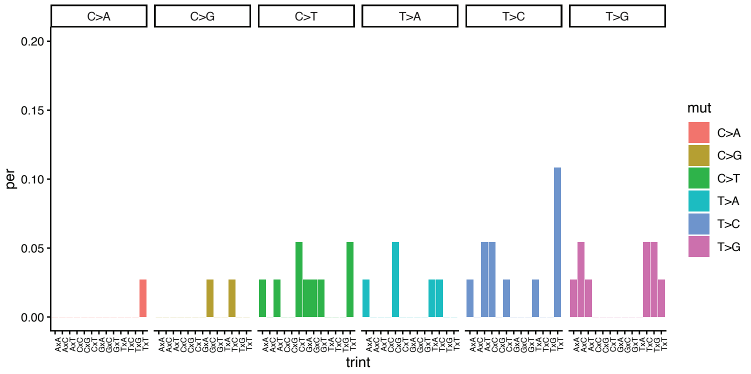

### Cohort-2

#### Patient 18 (substracted from Day -16)

No mutation at Day +1

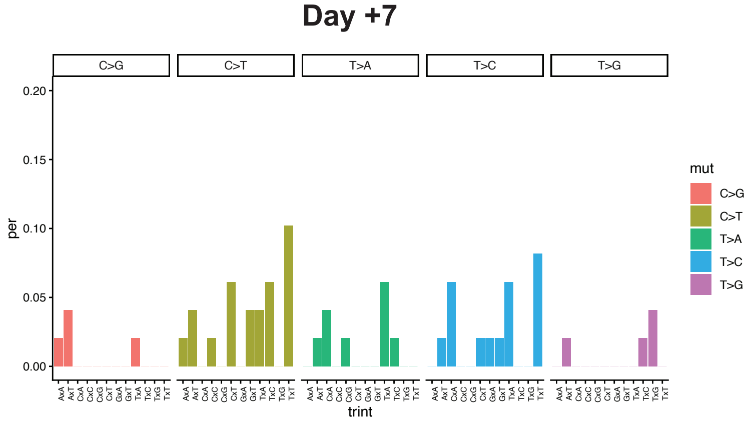


#### Patient 21 (substracted from Day -11)

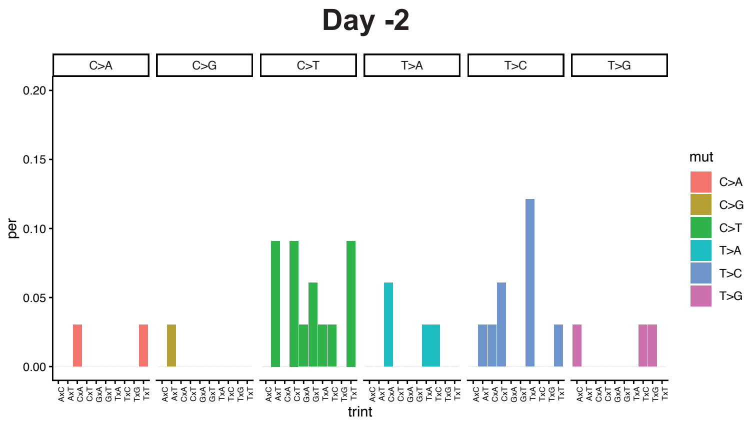


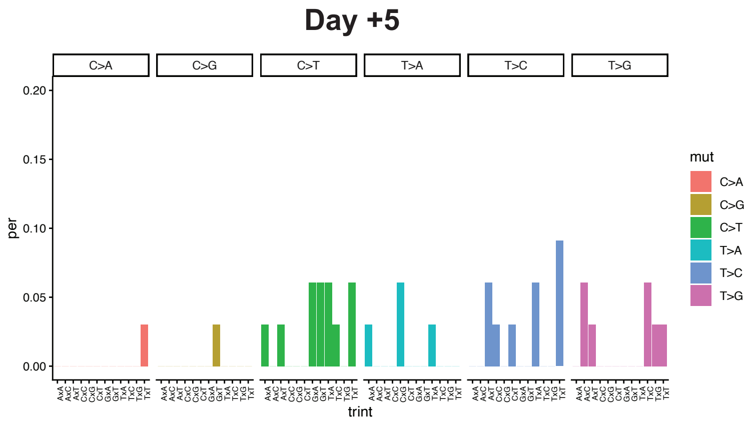


No mutations on Patient 2, 13, 19, and 20

### Supplementary Figure 9

#### Cohort-1

##### Patient 1 [Conditioning start Day -6, Transplant Day 0]

Day 0

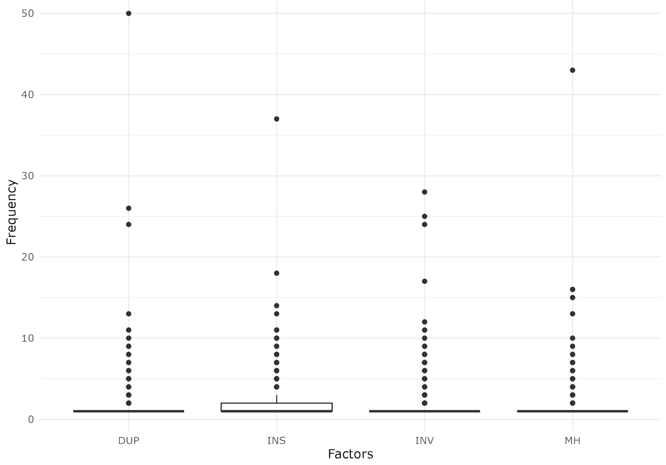


Day +7

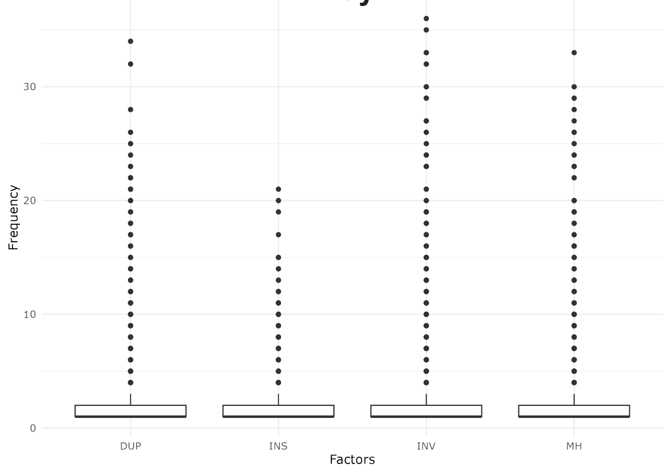


Day +14

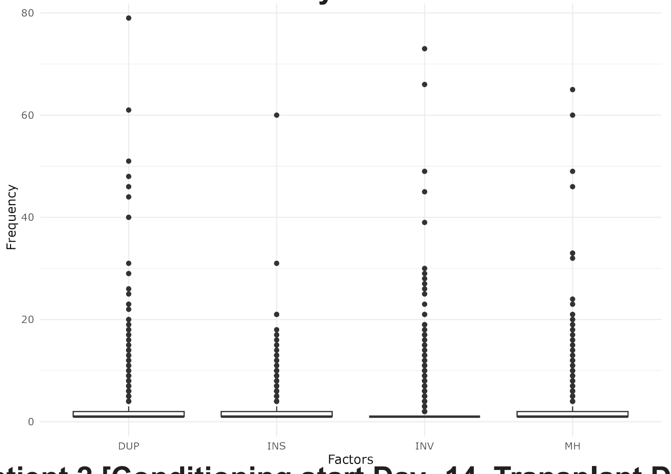


##### Patient 2 [Conditioning start Day -14, Transplant Day 0]

Day -14

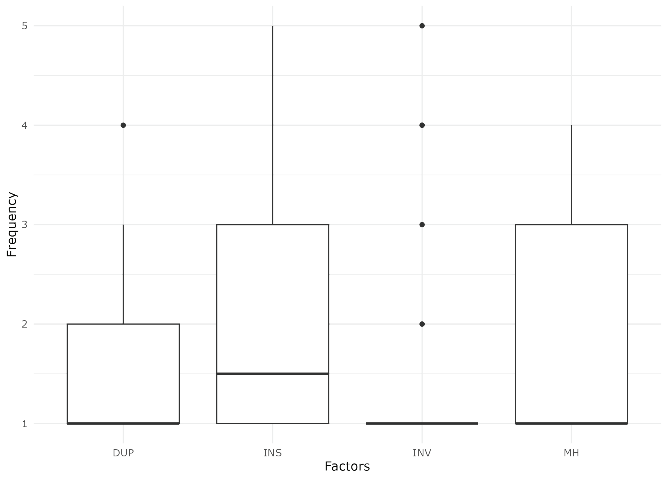


Day -7

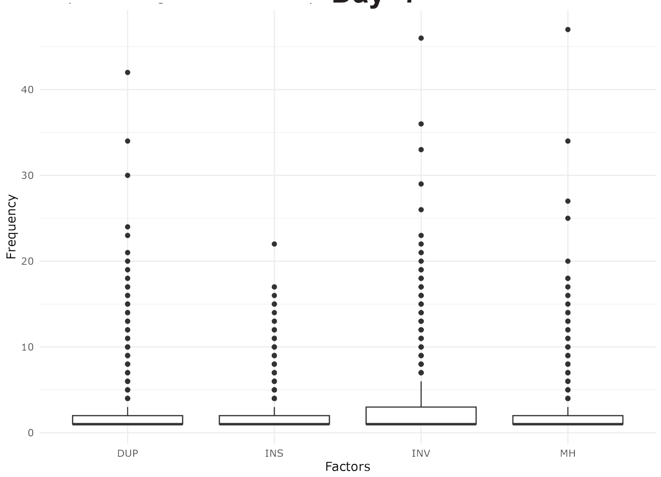


Day +28

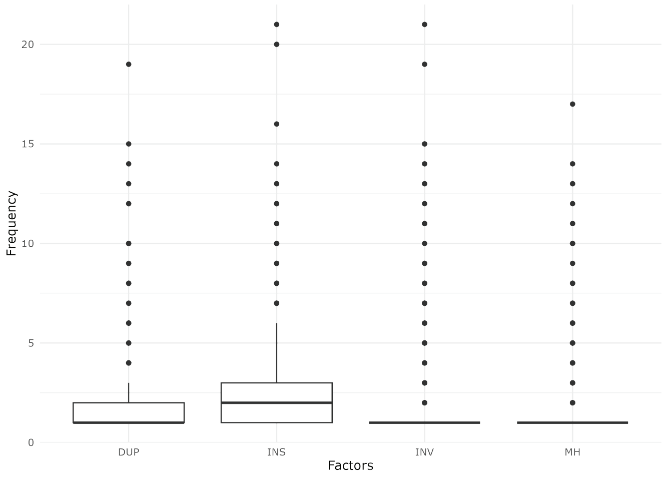


Day +35

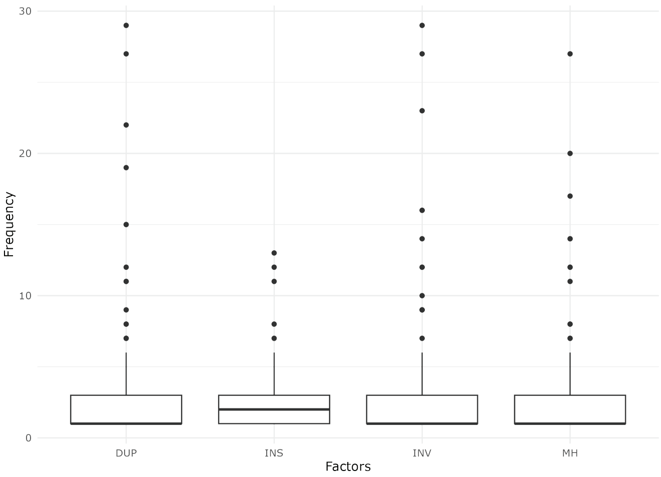

### Cohort-1

#### Patient 4 [Conditioning start Day -6, Transplant Day 0]

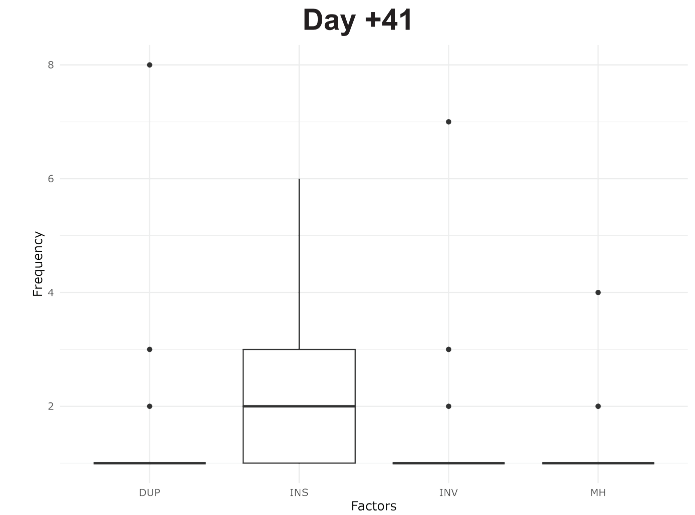


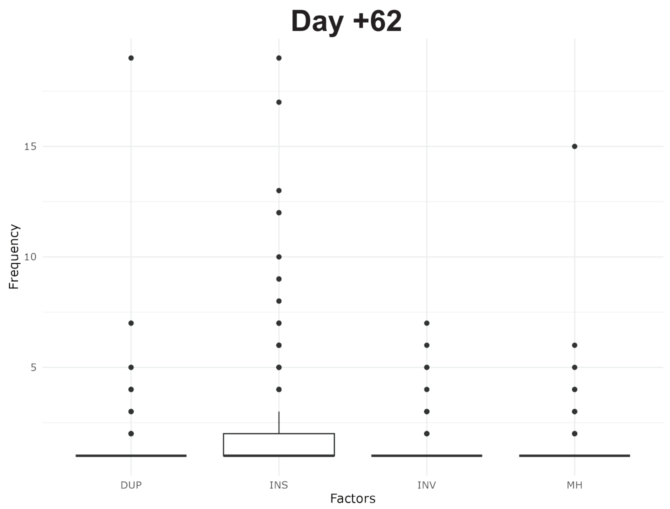


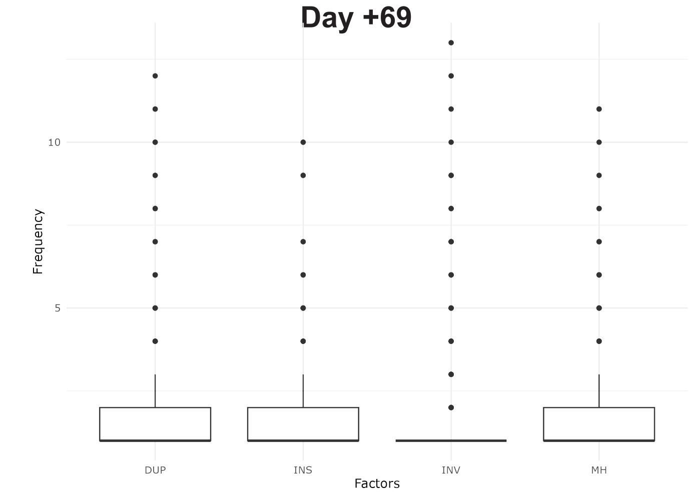


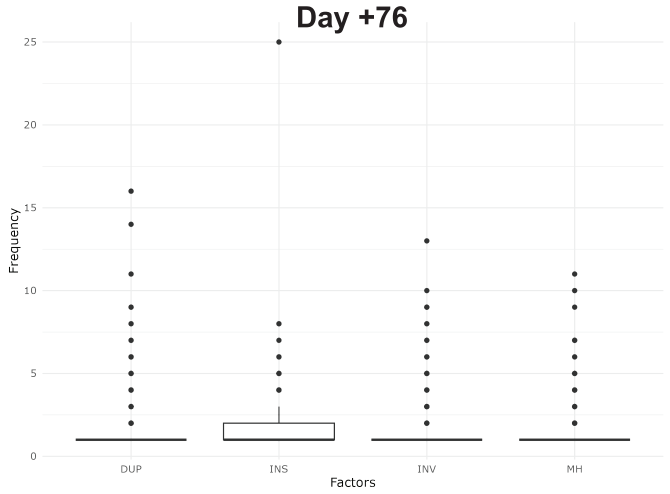


#### Patient 5 [Conditioning start Day -117 Transplant Day 0]

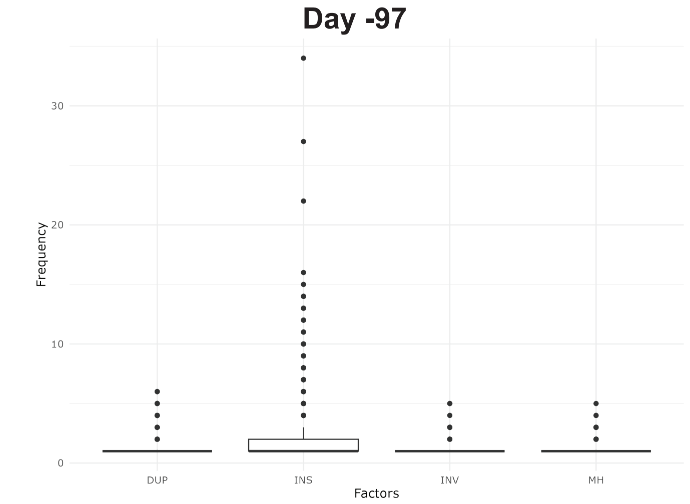


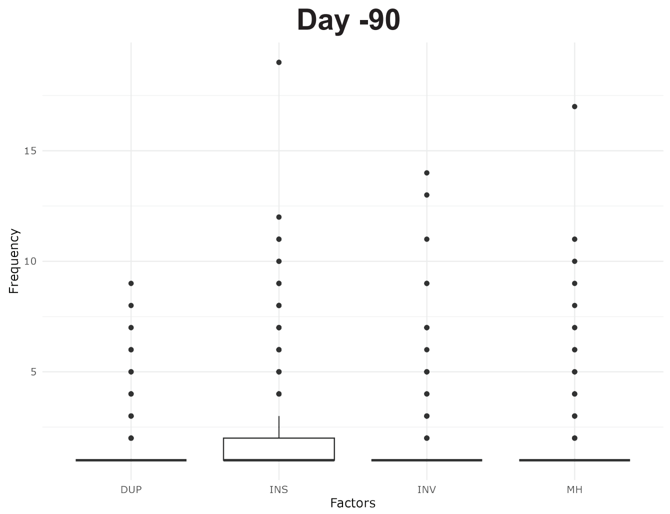


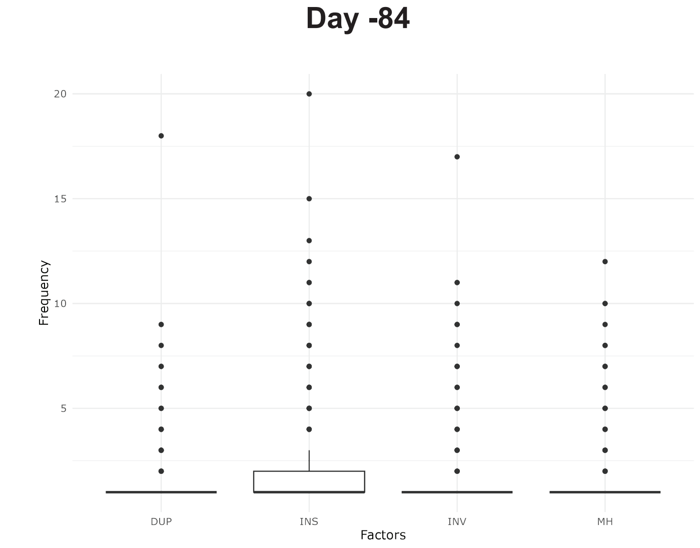


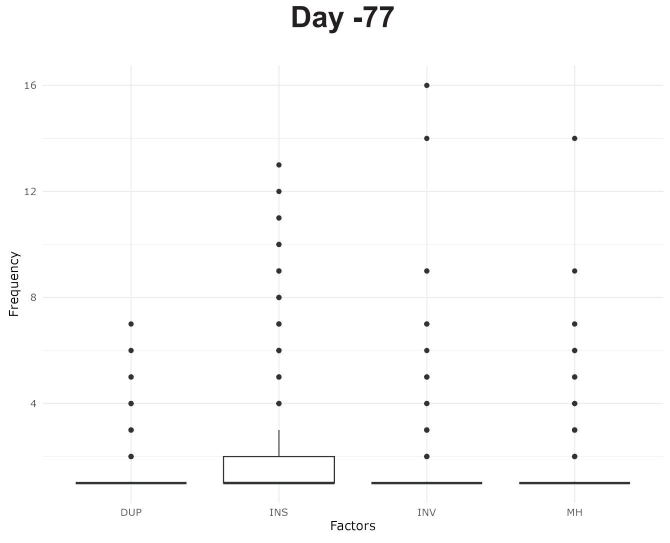

### Cohort-1

#### Patient 6 [Conditioning start Day -6, Transplant Day 0]

##### Day +63

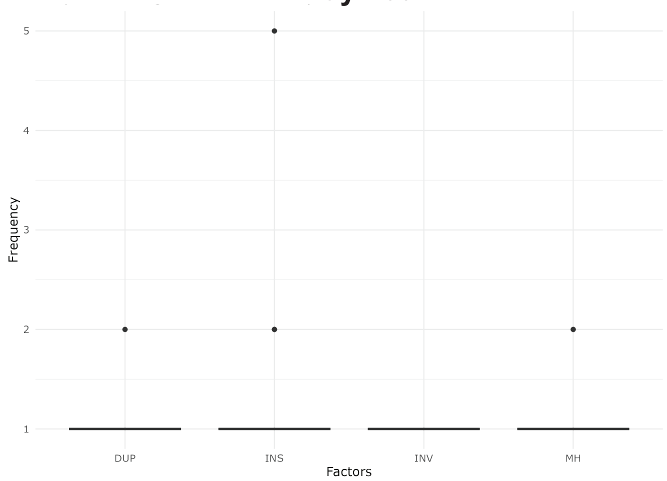


##### Day +70

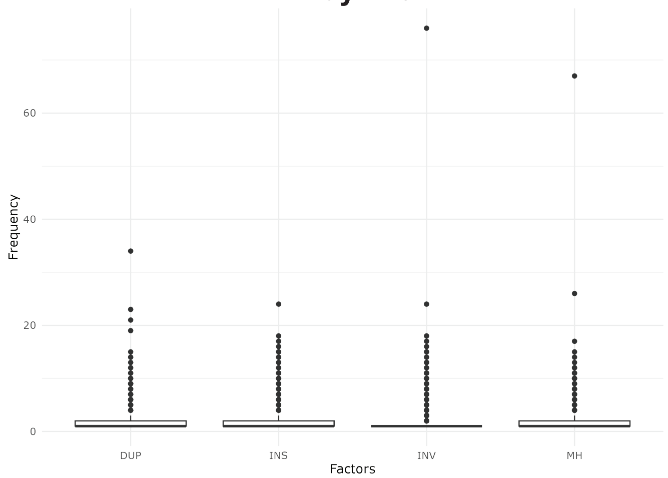


##### Day +77

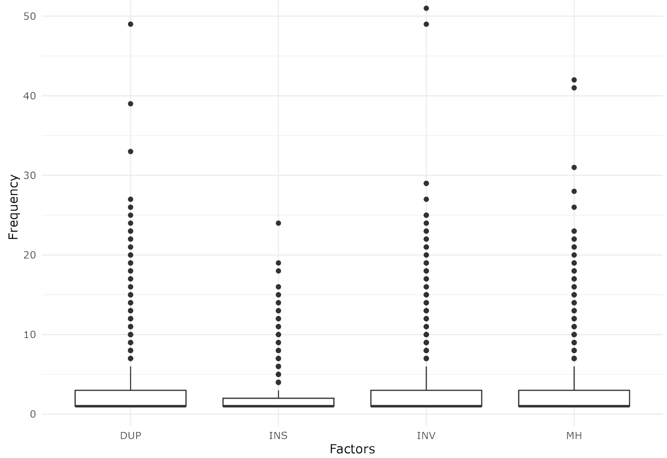


#### Patient 7 [Conditioning start Day -6 Transplant Day 0]

##### Day +34

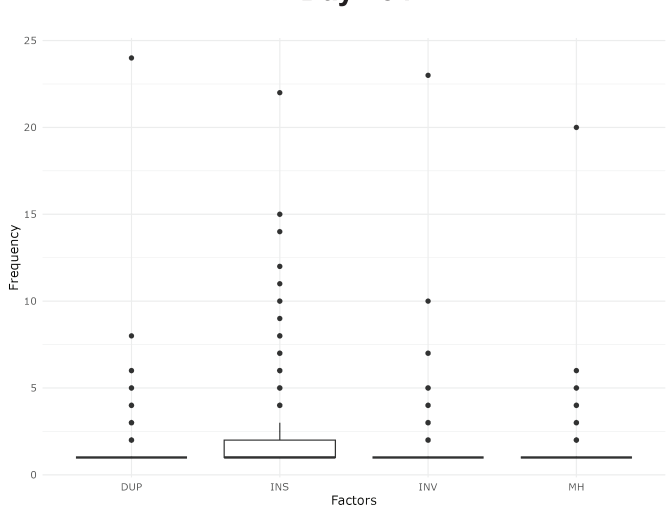


##### Day +41

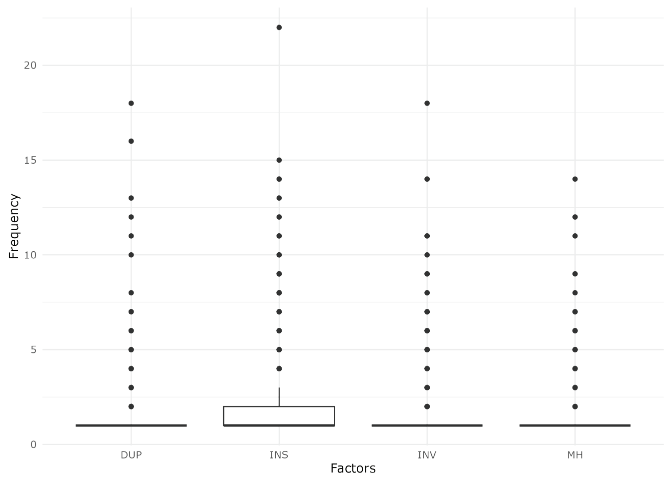


##### Day +48

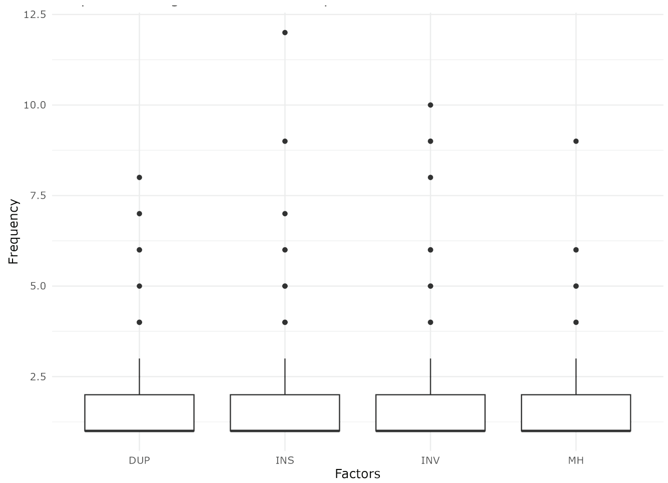


##### Day +55

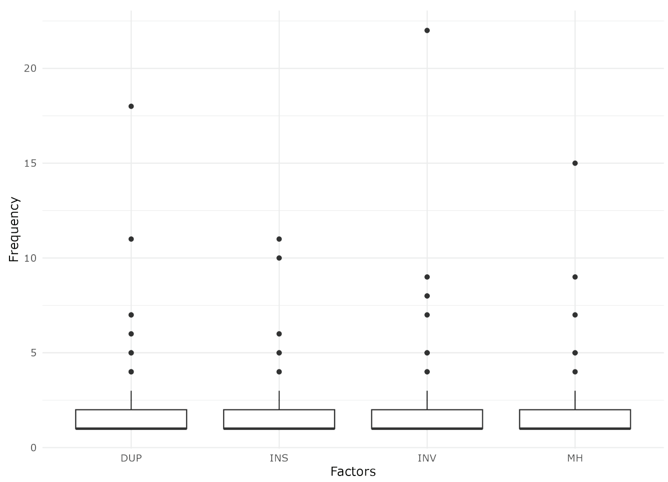

### Cohort-1

#### Patient 9 [Conditioning start Day -5 Transplant Day 0]

##### Day +14

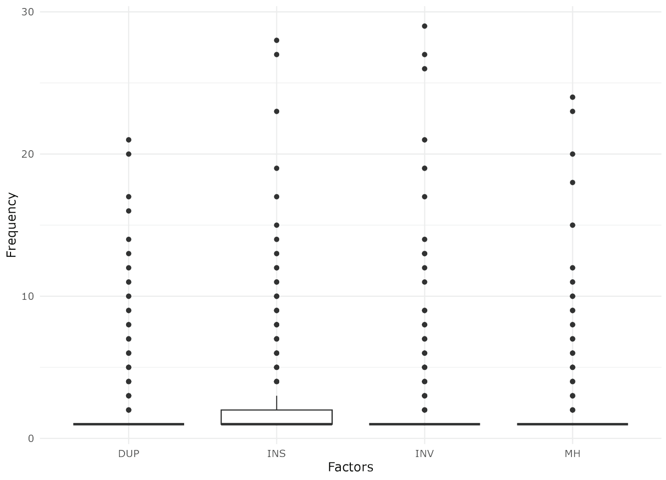


##### Day +21

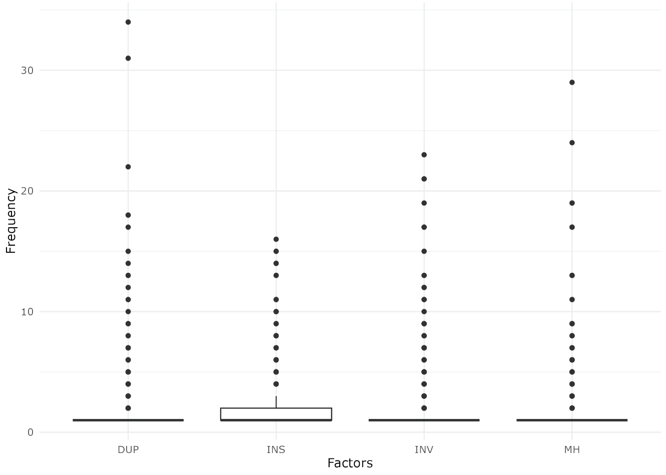


##### Day +28

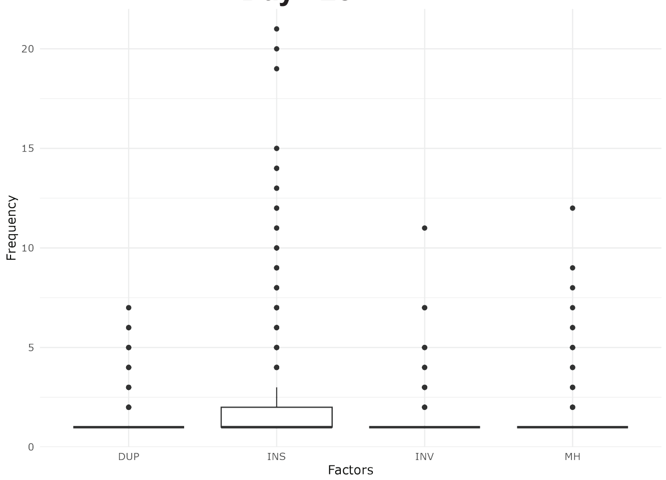


##### Day +35

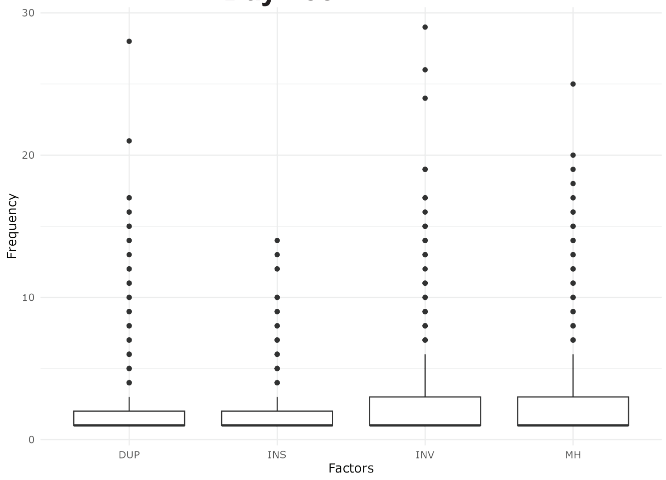


#### Patient 10 [Conditioning start Day -6 Transplant Day 0]

##### Day +34

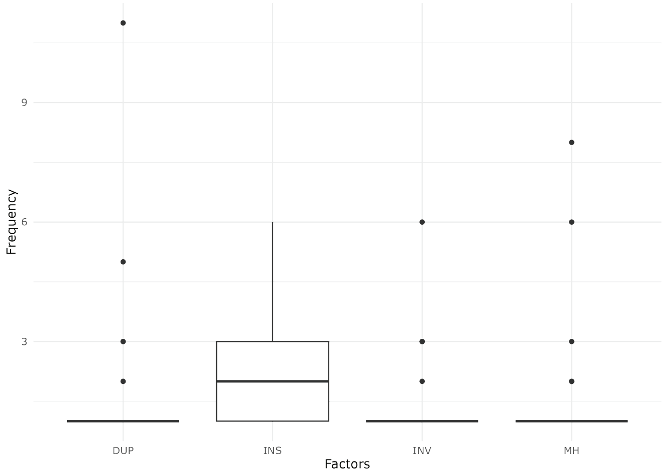


##### Day +41

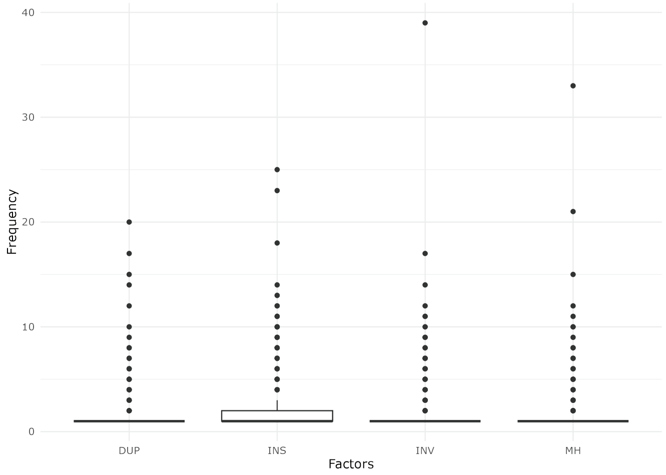


##### Day +55

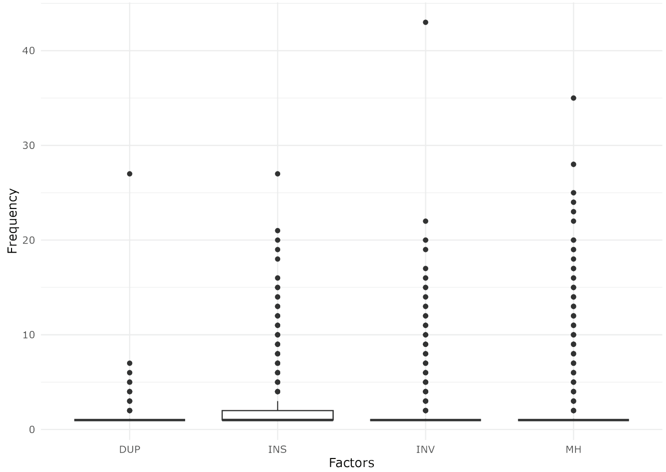

**Supplementary Figure 10**

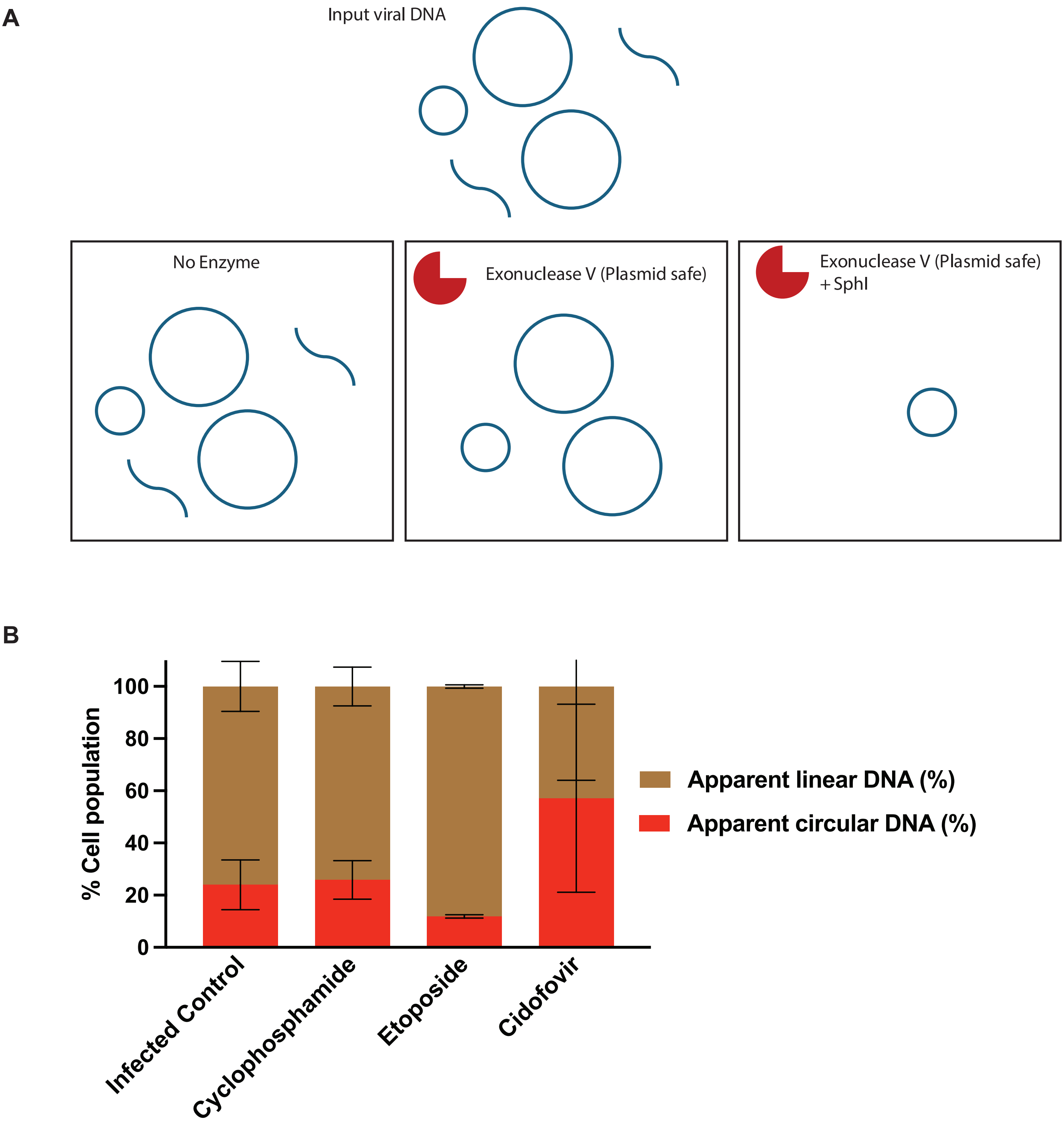

#### Supplementary Figure 11

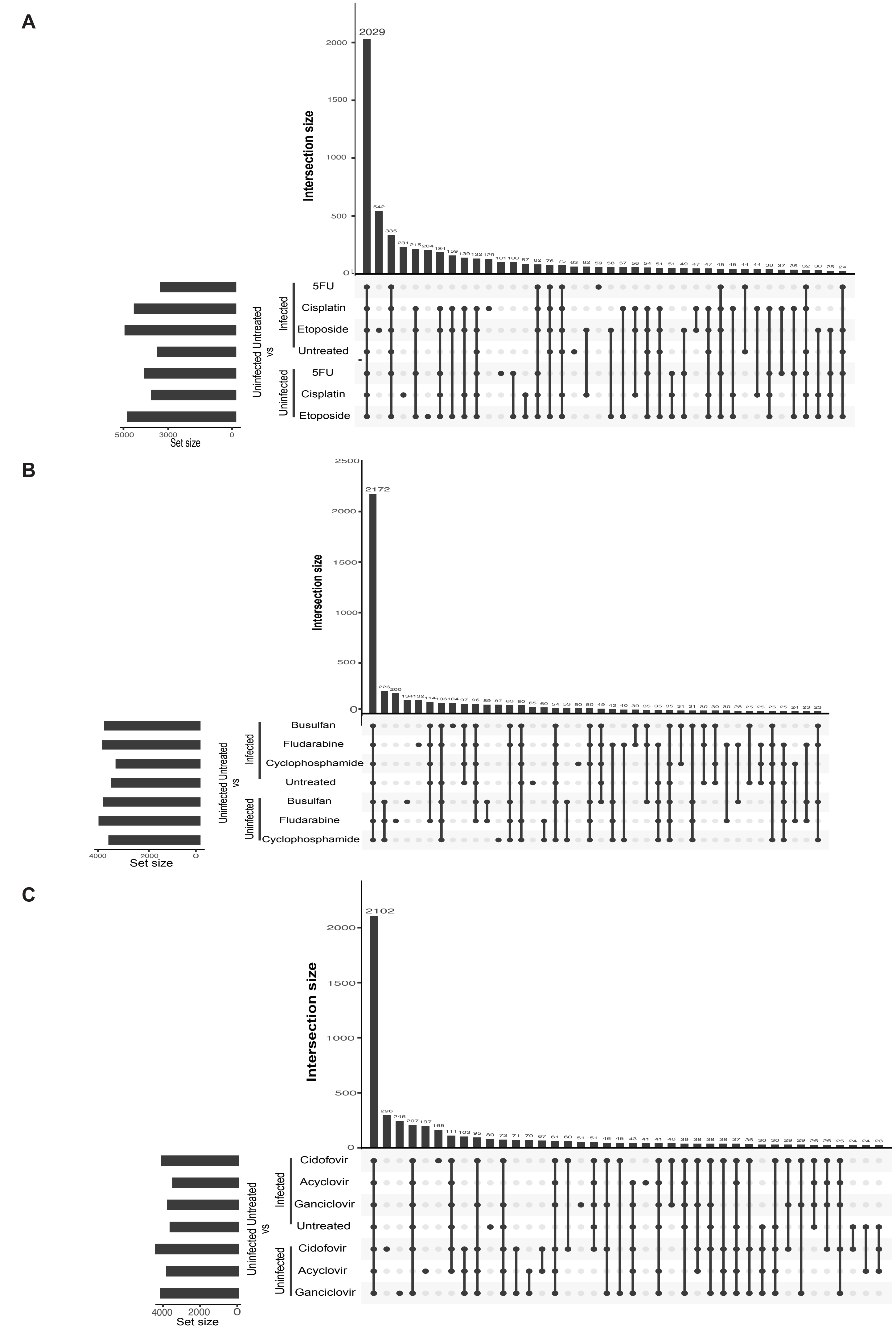
