## Supplementary Table 1 for "Therapeutic-induced chromatin remodeling via AP-1/SWI-SNF enhances BK Polyomavirus replication in urothelial cells"

| **Drugs** | **IC_10_ value** | **IC_20_ value** |
| --- | --- | --- |
| Cyclophosphamide | 3 μM | 10 μM |
| Fludarabine | 4 μM | 5 μM |
| Busulfan | 2.5 μM | 10 μM |
| Etoposide | 10 μM | 18 μM |
| Cisplatin | 0.5 μM | 2.5 μM |
| 5-FU | 0.5 μM | 1 μM |
| Ganciclovir | 70 μM | 105 μM |
| Acyclovir | 10 μM | 75 μM |
| Cidofovir | 25 μM | 37 μM |
