## Supplementary Table 2 for "Therapeutic-induced chromatin remodeling via AP-1/SWI-SNF enhances BK Polyomavirus replication in urothelial cells"

Clinical-event and urine-sampling timeline for Cohort 1

| **Patient ID** | **Conditioning start relative to transplantation** | **Transplantation** | **Relative urine sample collection day(s)** |
| --- | --- | --- | --- |
| Patient 1 | Day -6 | Day 0 | Day 0, Day +7, Day +14 |
| Patient 2 | Day -14 | Day 0 | Day -14, Day -7, Day +28, Day +35 |
| Patient 4 | Day -6 | Day 0 | Day +41, Day +62, Day +69, Day +76 |
| Patient 5 | Day -117 | Day 0 | Day -97, Day -90, Day -84, Day -77 |
| Patient 6 | Day -6 | Day 0 | Day +63, Day +70, Day +77 |
| Patient 7 | Day -6 | Day 0 | Day +34, Day +41, Day +48, Day +55 |
| Patient 9 | Day -5 | Day 0 | Day +14, Day +21, Day +28, Day +35 |
| Patient 10 | Day -6 | Day 0 | Day +34, Day +41, Day +55 |
