## Supplementary Table 3 for "Therapeutic-induced chromatin remodeling via AP-1/SWI-SNF enhances BK Polyomavirus replication in urothelial cells"

Clinical-event and urine-sampling timeline for Cohort 2

| **Patient ID** | **Conditioning start relative to transplantation** | **Transplantation** | **Relative urine sample collection day(s)** |
| --- | --- | --- | --- |
| Patient 1 |  | Day 0 | Day -16, Day -2, Day +6 |
| Patient 2 |  | Day 0 | Day -10, Day -5 |
| Patient 3 |  | Day 0 | Day -10, Day -3, Day +4, Day +18 |
| Patient 4 |  | Day 0 | Day -4, Day +4, Day +11 |
| Patient 5 |  | Day 0 | Day -7, Day +18 |
| Patient 6 |  | Day 0 | Day -22, Day -2, Day +5, Day +18 |
| Patient 7 |  | Day 0 | Day -10, Day +18 |
| Patient 8 |  | Day 0 | Day -17, Day -3, Day +4, Day +11 |
| Patient 9 |  | Day 0 | Day -10, Day -1, Day +6 |
| Patient 10 |  | Day 0 | Day -10, Day +18 |
| Patient 11 |  | Day 0 | Day -7, Day +2, Day +7 |
| Patient 12 |  | Day 0 | Day -8, Day +7 |
| Patient 13 |  | Day 0 | Day -15, Day +17 |
| Patient 14 |  | Day 0 | Day -7, Day -1, Day +6 |
| Patient 15 |  | Day 0 | Day -10, Day -2, Day +5, Day +12 |
| Patient 16 |  | Day 0 | Day -7, Day +7 |
| Patient 17 |  | Day 0 | Day -8, Day +5 |
| Patient 18 |  | Day 0 | Day -16, Day +1, Day +7 |
| Patient 19 |  | Day 0 | Day -15, Day -1, Day +6 |
| Patient 20 |  | Day 0 | Day -14, Day +1, Day +7 |
| Patient 21 |  | Day 0 | Day -11, Day -2, Day +5 |
